# One-carbon metabolism controls colorectal cancer cell state and chemosensitivity through chromatin remodeling

**DOI:** 10.64898/2026.09.30.755690

**Authors:** Laura Vízkeleti, Csaba Kiss, Viktória Tisza, Anna Zichy, Arnold Markovics, Alexandra Kalmar, Pál Szabó, István Csabai, Lőrinc Pongor, Sándor Spisák, Béla Molnár

## Abstract

Folate is co-administered with fluoropyrimidines in colorectal cancer to potentiate thymidylate synthase inhibition, yet its therapeutic effects are paradoxically inconsistent across disease and treatment settings. Here we show that one-carbon metabolism remodels epithelial identity before treatment and thereby alters fluoropyrimidine response. Folate restoration shifted colorectal cancer cells along a polarized epithelial axis from a stem-like toward a differentiated state, whereas folate deprivation favored the opposite configuration. These pretreatment states differed markedly in 5-fluorouracil sensitivity. Folate restoration increased sensitivity across colorectal cancer cell lines and patient-derived organoids, whereas folate deprivation or methotrexate increased therapy-induced senescence. State-dependent sensitivity was not explained by proliferation alone, as stem-like cells were least responsive under the same conditions in which their S-phase occupancy was highest. Mechanistically, folate restoration increased methylation potential, supported EZH2-dependent H3K27me3 remodeling, reorganized chromatin accessibility and transcription-factor engagement, and reduced canonical WNT activity. In human colorectal tissue, stem-like program activity was higher and differentiation program activity lower in carcinoma than in normal mucosa, with adenomas intermediate, a pattern opposite to folate-induced remodeling. Folate is therefore not merely a cofactor for thymidylate synthase inhibition but an instructive metabolic regulator of the epithelial state that stratifies fluoropyrimidine response.

## Introduction

5-fluorouracil (5-FU) remains a cornerstone of treatment for colorectal cancer (CRC), forming the backbone of therapy in both adjuvant and metastatic disease, most commonly as infusional 5-FU/leucovorin (the de Gramont schedule) or as the fluoropyrimidine component of combination regimens such as FOLFOX (1,2). In clinical practice, 5-FU is routinely combined with folinic acid (leucovorin), based on the longstanding model that folate enhances fluoropyrimidine efficacy by providing the reduced folate cofactor required to stabilize the ternary complex between FdUMP, thymidylate synthase and 5,10-methylenetetrahydrofolate, thereby prolonging target inhibition (1). However, folate participates in multiple interconnected one-carbon pathways that extend beyond thymidylate synthesis. Folate-dependent one-carbon metabolism transfers one-carbon units between biosynthetic pathways and, through its connection to the methionine cycle, supports the generation of S-adenosylmethionine (SAM), the principal methyl donor for cellular methylation reactions. Thus, in addition to supporting nucleotide production, folate availability can influence cellular methylation capacity, providing a potential link between nutrient availability, chromatin regulation and epithelial cell identity (3–6). These broader functions provide a plausible basis for the complex and sometimes paradoxical effects of folate availability on tumor behavior and therapeutic response observed across experimental and clinical settings (7,8). Whether folate-dependent regulation of methylation capacity and chromatin state contributes to fluoropyrimidine response has remained unresolved.

Despite its widespread clinical adoption, the therapeutic benefit of folinic acid co-administration has remained difficult to predict across disease settings. A meta-analysis of metastatic colorectal cancer trials demonstrated improved response rates and a modest overall survival benefit with 5-FU/leucovorin compared with 5-FU alone (9). By contrast, randomized adjuvant trials have yielded conflicting results, with several randomized studies reporting minimal or no improvement in disease-free or overall survival following the addition of folinic acid (10,11). Together, these observations indicate that the therapeutic value of folate modulation is strongly context-dependent and is not readily predicted by the canonical thymidylate synthase model alone, raising the possibility that additional one-carbon-dependent processes influence treatment response.

A potential source of such context dependence is epithelial cell state. Colorectal tumorigenesis is frequently initiated by loss of APC, driving constitutive Wnt signaling and destabilizing epithelial identity (12,13). Rather than simply blocking differentiation, APC loss promotes epithelial plasticity, enabling tumor cells to reversibly switch between an aberrant, stem-like state and a differentiated state (14,15). Seminal studies spanning intestinal organoid models, genetically engineered mouse models, and single-cell analyses of primary human tumors have shown that epithelial cells reversibly acquire regenerative, fetal-like, and stem-like transcriptional programs in response to oncogenic signaling, tissue injury, and therapeutic stress (16–20). These states are not equivalent to the LGR5-marked homeostatic stem compartment of the normal crypt. In APC-mutant epithelium, SOX9-associated stem-like programs can be aberrantly sustained outside the canonical homeostatic stem-cell hierarchy (14,15,19,21). These transitions are not phenotypically neutral, since movement toward less differentiated states is associated with metastatic competence and the emergence of drug-tolerant populations that contribute to disease persistence and relapse (22,23). Together, these studies shifted the conceptual framework of colorectal cancer from a static hierarchical model toward one in which epithelial identity remains continuously remodelable throughout tumor progression and therapeutic adaptation. Yet the upstream metabolic signals governing movement within this epithelial state space have remained largely unknown.

Identifying these upstream metabolic regulators has direct implications for fluoropyrimidine therapy. More differentiated CRC cells are generally more sensitive to 5-FU, whereas less differentiated and more plastic states exhibit reduced responsiveness (23–25). Since epithelial states also differ in proliferation, it remains unclear whether this difference in sensitivity simply reflects proliferation or is tied to epithelial identity itself, a distinction with direct consequences for how folate is expected to act. Recent work has shown that the early adaptive response to 5-FU, including rapid mitochondrial remodeling in surviving cells, is distinct from the later emergence of senescent or persistently arrested states (26). This distinction raises the possibility that the cellular state established before treatment may influence not only acute fluoropyrimidine sensitivity, but also the subsequent fate of treatment-surviving cells. Whether metabolic inputs such as folate availability shape both phases through their effects on epithelial state remains unknown. Despite the routine incorporation of folinic acid into fluoropyrimidine-based chemotherapy, its use is not stratified according to tumor metabolic state or epithelial plasticity and is therefore applied broadly rather than guided by the cellular context in which treatment acts. Building on our previous work showing that folate availability influences gene expression and DNA methylation in colorectal cancer cells and patient tissues (27–29), we asked whether one-carbon metabolism acts as an instructive regulator of epithelial plasticity and thereby establishes the cellular context in which fluoropyrimidine response varies. Addressing this question required experimental systems capable of distinguishing genuine state remodeling from changes in population composition and of resolving the regulatory events that precede the emergence of a new epithelial-state distribution.

To address this, we combined the HT29 dual endogenous reporter (DER) system, in which fluorescent cassettes integrated at the endogenous SOX9 and KRT20 loci enable live identification and prospective isolation of SOX9-associated stem-like (SCL) and KRT20-associated differentiated (DIFF) epithelial states (30), with time-resolved single-cell transcriptome and chromatin-accessibility profiling across ten successive time points. This framework allowed us to reconstruct the molecular trajectory connecting reporter-defined states and to resolve the regulatory events preceding their population-level redistribution. We show that folate restoration drives progressive remodeling toward differentiated epithelial states, whereas folate deprivation favors SCL-associated and adaptive intermediate states. Mechanistically, this transition is coupled to increased methylation potential and EZH2-dependent H3K27me3 remodeling. Time-resolved regulatory profiling further revealed extensive enhancer and transcription-factor reorganization accompanied by attenuation of canonical WNT signaling. This remodeling occurs along a polarized SCL–DIFF axis that stratifies fluoropyrimidine sensitivity, while the same component programs reorganize in the opposite direction across human normal mucosa, adenoma and carcinoma. Together, these findings identify one-carbon metabolism as an upstream regulator of epithelial plasticity and reveal a metabolic-epigenetic axis linking folate availability, epithelial state and fluoropyrimidine response.

## Results

### Folate remodels stem-like and differentiation-linked epithelial states in colorectal cancer models

To determine whether folate availability directs epithelial identity in colorectal cancer, we used the HT29 dual endogenous reporter (DER) system. In this system, fluorescent reporter cassettes were integrated at the endogenous *SOX9* and *KRT20* loci, allowing mKate2 to report stem-like (SCL) activity and GFP to report differentiated (DIFF) activity in live cells (30) (**Fig. 1A**). Folate restoration (FAᴸ, 2.25 μM folic acid for 48 h) shifted the distribution observed under folate starvation (FA⁰), reducing the SCL fraction by 10.8 percentage points and increasing the DIFF fraction by 12.5 percentage points (both *p* = 0.0004, two-sided Student’s *t*-test, *n* = 3 biological replicates; **Fig. 1B**). Representative cytometry and the full four-quadrant analysis are shown in **Fig. S1A,B**.

**Figure 1.**
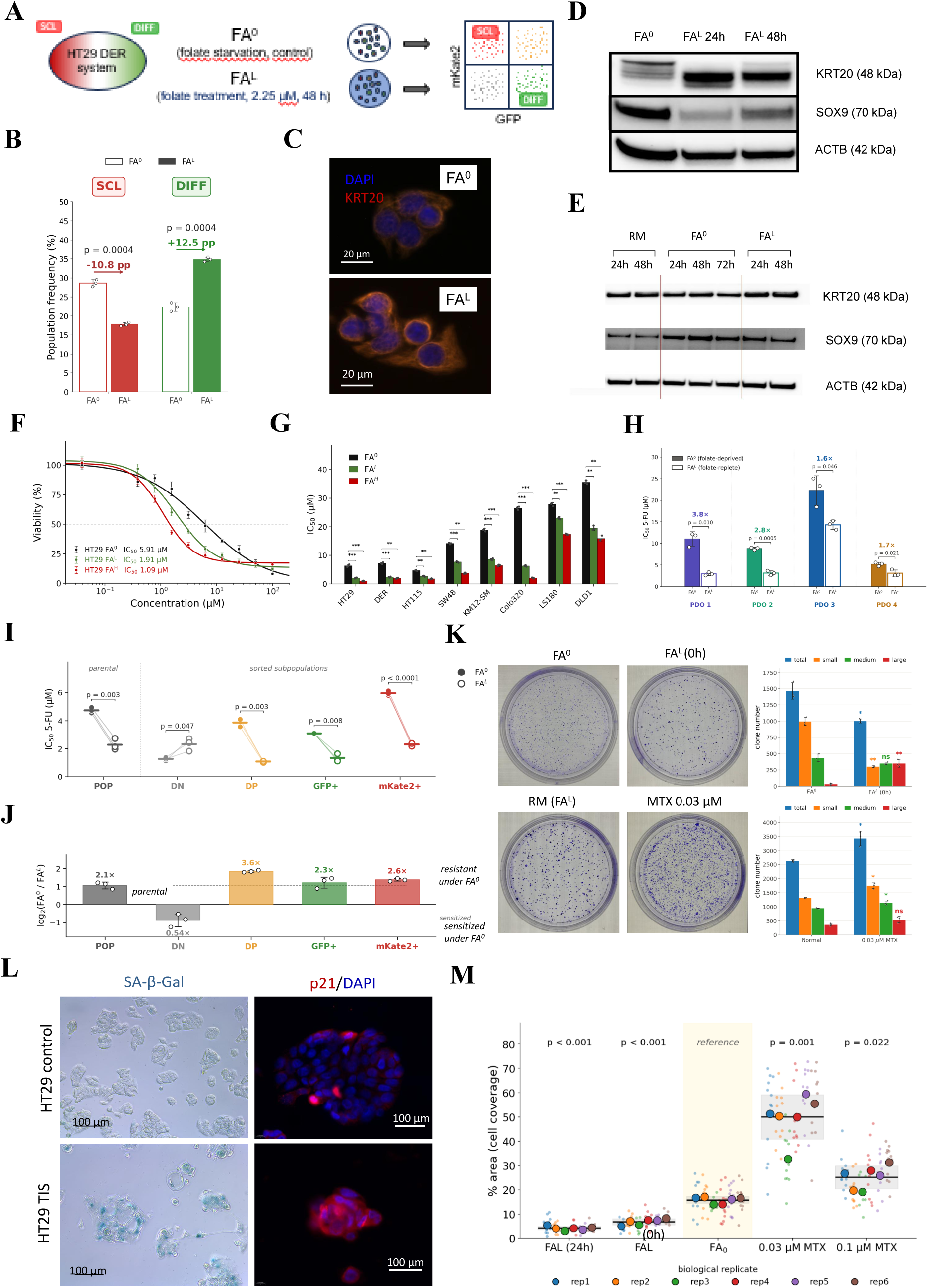
Folate shifts colorectal cancer epithelial state composition toward differentiation and increases fluoropyrimidine sensitivity. **A**, Schematic of the experimental design using the HT29 dual endogenous reporter (DER) system, in which mKate2 reports the SOX9-associated stem-like (SCL) state and GFP reports the KRT20-associated differentiated (DIFF) state. Cells were cultured under folate-starved (FA⁰) or folate-replete (FAᴸ; 2.25 μM folic acid for 48 h) conditions prior to flow cytometric analysis. **B**, Quantification of SCL and DIFF population frequencies under FA⁰ (open bars) and FAᴸ (filled bars) conditions. Percentage point (pp) changes relative to FA⁰ are indicated above the bars. Data are presented as mean ± SD (*n* = 3 biological replicates). Statistical significance was determined by two-sided Student’s *t*-test. Representative flow cytometry plots are shown in Fig. S1A, and an extended analysis including the double-positive and double-negative populations in Fig. S1B. **C**, Representative immunofluorescence images of KRT20 (red) and DAPI (blue) in HT29 cells cultured under FA⁰ and FAᴸ conditions. Increased KRT20 immunoreactivity following folate supplementation is consistent with induction of a differentiated epithelial phenotype. Scale bar, 20 μm. Images are representative of three independent experiments. **D**, Representative immunoblot analysis of SOX9 and KRT20 protein expression under FA⁰ and after 24 h and 48 h of folate supplementation (FAᴸ). SOX9 and KRT20 exhibit reciprocal regulation. ACTB served as a loading control. Corresponding transcript-level time-course data are shown in Fig. S1D. **E**, Representative immunoblot analysis of SOX9 and KRT20 protein expression in patient-derived colorectal cancer organoids cultured in regular medium (RM), under folate starvation (FA⁰), or with folate supplementation (FAᴸ) for the indicated durations, demonstrating a conserved folate-responsive differentiation program *ex vivo*. ACTB served as a loading control. **F**, Dose-response of HT29 cells to 5-FU following culture under FA⁰, FAᴸ (2.25 μM) or FAᴴ (22.5 μM) conditions. IC_50_ values were determined by nonlinear regression using a four-parameter logistic model. Data are presented as mean ± SD (n = 3 biological replicates). **G**, Comparison of 5-FU IC_50_ values across colorectal cancer cell lines cultured under FA⁰, FAᴸ, and FAᴴ conditions. IC_50_ values were derived by nonlinear regression using a four-parameter logistic model. Data are presented as mean ± SD (n = 3 biological replicates per cell line). Statistical significance was assessed by Welch’s two-sided t-test. Full dose-response curves are shown in Fig. S1E. **H**, 5-FU sensitivity of four patient-derived colorectal cancer organoid (PDO #1-4) lines cultured under FA⁰ and FAᴸ conditions. IC_50_ values were determined by nonlinear regression. Data are presented as mean ± SD (n = 3 independent experiments). Statistical significance was assessed by Welch’s two-sided t-test. **I**, 5-FU sensitivity of the unsorted DER population (POP) and DER-sorted subpopulations (DN = double negative, DP = double positive, GFP⁺ = DIFF, and mKate2⁺ = SCL) cultured under FA⁰ and FAᴸ conditions. IC_50_ values for 5-FU are shown; filled circles, FA⁰; open circles, FAᴸ, with paired conditions connected. Data are presented as mean ± SD (n = 3 biological replicates); statistical significance was assessed by Welch’s two-sided t-test (p values indicated). **J**, Fold change in 5-FU sensitivity between FA⁰ and FAᴸ for each population, shown as log_2_(FA⁰/FAᴸ). Positive values, annotated as resistant under FA⁰, indicate a higher IC_50_ under FA⁰ than FAᴸ, that is, sensitization after folate supplementation; negative values, annotated as sensitized under FA⁰, indicate increased resistance after folate supplementation. Values above each bar denote the corresponding linear fold change; the dashed line marks the unsorted DER (POP) level. Data are mean ± SD (n = 3 biological replicates). **K**, Clonogenic survival of HT29 cells treated with 20 µM 5-FU following folate or antifolate manipulation. Representative crystal violet–stained dishes are shown for FA⁰, FAᴸ (0 h before 5-FU), regular medium with 2.25 μM folate supplementation (RM (FAᴸ)), and MTX (0.03 µM), alongside automated size-resolved colony quantification for the two paired comparisons (FA⁰ versus FAᴸ 0 h; normal medium versus 0.03 µM MTX). Colony classes are small (orange), medium (green), large (red), and total (blue, the sum of the three size classes); see Fig. S1H–J for all conditions and further details. Data are presented as mean ± SD (n = 3 biological replicates), with dots showing individual values. Statistical significance was assessed per size class by Welch’s two-sided t-test against normal medium (antifolate arm) or FA⁰ (folate arm); asterisks are shown in the color of the corresponding size class. \**p* < 0.05, \*\**p* < 0.01; ns, not significant. **L**, Validation of therapy-induced senescence (TIS) in HT29 cells. Representative senescence-associated β-galactosidase (SA-β-Gal) staining (left) and p21 immunofluorescence with DAPI counterstain (right) in untreated control cells and TIS cells at day 12 (7-day recovery phase following a 5-day course of 20 μM 5-FU treatment). Scale bars, 100 μm. **M**, Quantification of TIS-associated cell coverage area (%) at day 12, following a 5-day course of 20 μM 5-FU and a 7-day recovery phase, across two experimental arms, a folate arm (FAᴸ 24 h, FAᴸ 0 h and FA⁰) and an antifolate arm (0.03 μM and 0.1 μM MTX). FA⁰ served as the reference for the folate arm and untreated normal medium for the antifolate arm. Small dots represent individual fields; large dots represent biological replicate means (*n* = 6), with statistics computed on replicate means. Horizontal lines indicate mean ± SD. Statistical significance was assessed by Welch’s two-sided *t*-test. Unless otherwise indicated, \**p* < 0.05, \*\**p* < 0.01, \*\*\**p* < 0.001, \*\*\*\**p* < 0.0001; ns, not significant.

This redistribution was accompanied by reciprocal changes in endogenous markers. KRT20 staining increased, with partial redistribution to the membrane, at single-cell resolution (**Fig. 1C**). In an independent time-course experiment, reporter-defined population frequencies shifted gradually, with no significant difference detected at 4 h, partial redistribution at 24 h and a pronounced shift at 48 h, reaching −11.9 pp for SCL and +10.8 pp for DIFF (p < 0.0001 and p = 0.0001; **Fig. S1C**). The underlying transcriptional response was considerably faster. SOX9 transcript levels declined progressively, whereas KRT20 rose rapidly, peaking at approximately 2.2-fold above baseline after 4 h (**Fig. S1D**), and immunoblotting confirmed the corresponding protein changes (**Fig. 1D**). Because the DER readout depends on synthesis, folding and maturation of the encoded fluorescent proteins, it reports epithelial state with a delay relative to transcription. The rapid transcriptional response and slower redistribution of reporter-defined populations are therefore temporally consistent with progressive folate-induced epithelial-state remodeling. Patient-derived colorectal cancer organoids showed the same opposing SOX9 and KRT20 responses (**Fig. 1E**). Folate restoration therefore favors differentiation-associated epithelial states in both HT29 cells and patient-derived organoids.

### Folate availability modulates fluoropyrimidine sensitivity across epithelial states

Having established that folate redistributes reporter-defined epithelial states, we asked whether this remodeling is accompanied by altered fluoropyrimidine sensitivity. Folate restoration increased HT29 sensitivity to 5-fluorouracil (5-FU), lowering the IC_50_ from 5.91 μM under FA⁰ to 1.91 μM under FAᴸ and 1.09 μM under high-folate conditions (FAᴴ, 22.5 μM), corresponding to 3.1- and 5.4-fold sensitization, respectively, within a single genetic background (**Fig. 1F**). The HT29 IC_50_ estimates were derived by four-parameter logistic regression from three biological replicates.

The same direction of response was observed across seven additional CRC cell-line models, including the HT29-derived DER reporter line, although its magnitude varied (**Fig. 1G**; **Fig. S1E**; **Table S1**). In patient-derived organoids, folate restoration reduced 5-FU IC_50_ values by approximately 3.8-, 2.8-, 1.6- and 1.7-fold in PDO1–PDO4 (**Fig. 1H**), and HT29 spheroids responded comparably (**Fig. S1F**).

Pharmacological inhibition of folate metabolism produced a reciprocal shift in 5-FU response. Relative to FA⁰, folate restoration lowered the 5-FU IC_50_, whereas MTX pretreatment increased it approximately sevenfold at 0.03 μM and 2.4-fold at 0.1 μM, with the smaller shift at the higher dose being consistent with increasing direct MTX cytotoxicity at approximately one third of its own IC_50_ of 0.33 μM (**Fig. S1G**). Thus, folate restoration and pharmacological folate blockade shifted fluoropyrimidine response in opposing directions.

To determine whether pretreatment epithelial state stratifies 5-FU sensitivity and whether folate alters the response within each state, we measured 5-FU responses in prospectively isolated DER populations. Under FA⁰, SCL cells were the least sensitive (IC_50_ approximately 6 μM), double-negative (DN) cells the most sensitive (approximately 1.2 μM), and DIFF and double-positive (DP) cells intermediate (approximately 3–4 μM) (**Fig. 1I**). Prospectively isolated pretreatment states displayed distinct baseline 5-FU sensitivities, which were further modified by folate restoration in a state-dependent manner, with DP cells showing the strongest sensitization and DN cells responding in the opposite direction. (**Fig. 1J**). DN cells were the exception, their IC_50_ rising under FAᴸ. Fluoropyrimidine sensitivity therefore differs between epithelial states, and the effect of folate is state dependent rather than uniform.

### Folate availability influences clonogenic recovery and therapy-induced senescence after fluoropyrimidine treatment

We first examined clonogenic recovery after treatment with 20 μM 5-FU, comparing simultaneous folate supplementation (0 h) or preconditioning for 24 or 48 h with continued folate starvation. Colonies were segmented and assigned to size classes using the automated workflow shown in **Fig. S1H**. Relative to FA⁰, simultaneous supplementation and 24 h preconditioning reduced total colony numbers and depleted small colonies while increasing larger ones (**Fig. S1I**). After 48 h, total colony numbers instead exceeded the FA⁰ reference, whereas the shift toward larger colonies persisted. Thus, the effect of folate on colony number depended on preconditioning duration, while its effect on colony-size distribution was maintained.

We then tested the reciprocal perturbation. Relative to normal medium, MTX pretreatment increased colony numbers at both concentrations tested, with increases across all three size classes but no marked shift in their relative distribution (**Fig. S1J**). The representative dishes and size-resolved quantification in **Fig. 1K** summarize both arms, with simultaneous folate supplementation reducing colony numbers relative to FA⁰ and 0.03 μM MTX increasing them relative to normal medium. Counts from three biological replicates were compared within each size class by Welch’s two-sided *t*-test.

Having shown that folate availability rapidly alters epithelial gene expression, subsequently remodels epithelial state (**Fig. 1B–E**; **Fig. S1C,D**), and modulates 5-FU sensitivity, we asked whether it might also predispose cells to different therapy-induced senescence (TIS) outcomes. In a model established by 72 h of folate deprivation before prolonged 5-FU exposure, SA-β-Gal staining and p21 expression supported a senescence-associated phenotype (**Fig. 1L**). The culture area occupied by TIS-associated cells was then quantified from brightfield images acquired across twelve predefined fields per dish, with the sampling strategy, representative images and corresponding segmentation masks shown in **Fig. S1K–M**, and additional p21 immunofluorescence in **Fig. S1N**. Quantification revealed a consistent reduction when folate was restored either 24 h before 5-FU exposure or at treatment onset, whereas MTX pretreatment increased the area occupied by these cells (**Fig. 1M**). Measurements were summarized at the biological-replicate level (n = 6). These findings link folate availability around treatment onset to the subsequent accumulation of TIS-associated cells, raising the possibility that pretreatment epithelial state contributes to this response.

Together, these results identify folate availability as a regulator of epithelial state before 5-FU exposure and of the cellular responses that follow treatment. Folate restoration favored differentiated states, increased 5-FU sensitivity and reduced the accumulation of TIS-associated cells, whereas folate deprivation favored the less sensitive SCL state. MTX increased 5-FU resistance, clonogenic recovery and TIS-associated cell coverage, extending the opposing effects of folate availability to pharmacological folate blockade. Folate pretreatment resulted in fewer but larger colonies after 5-FU treatment, whereas folate deprivation yielded more numerous but smaller colonies, separating the number of surviving clones from their subsequent expansion.

### Reporter-defined epithelial states span a continuous stem-like-to-differentiated axis

Time-resolved measurements indicated that folate-induced epithelial remodeling begins before it becomes apparent from reporter-defined population frequencies, with KRT20 transcription rising and SOX9 declining within 4 h of folate restoration (**Fig. S1D**), whereas redistribution of the SCL and DIFF populations emerged at 24 h and became pronounced by 48 h (**Fig. S1C**). To place this response in a population-level context, we first profiled HT29 cells under folate-starved (FA⁰), low-folate (FAᴸ) and high-folate (FAᴴ) conditions by bulk RNA-seq. This revealed a rapid transcriptional response that was largely shared between FAᴸ and FAᴴ at 48 h but diverged according to folate dose after prolonged exposure. The bulk response involved metabolic and cell-cycle-associated pathways but did not resolve epithelial-state-specific programs (**Fig. S2A–C; Table S2A**), prompting us to examine the response at single-cell resolution.

To define the molecular changes underlying this transition, we performed paired single-cell transcriptome and chromatin-accessibility profiling (10x Genomics Multiome ATAC + Gene Expression) of HT29 DER cells after folate starvation (FA⁰, 72 h) or 48 h of folate restoration (FAᴸ, 2.25 μM). Under each condition, we profiled the unsorted population (POP) together with prospectively isolated stem-like (SCL, top 2% mKate2⁺) and differentiated (DIFF, top 2% GFP⁺) fractions, yielding six populations spanning the principal epithelial states of the system (**Fig. 2A**).

**Figure 2.**
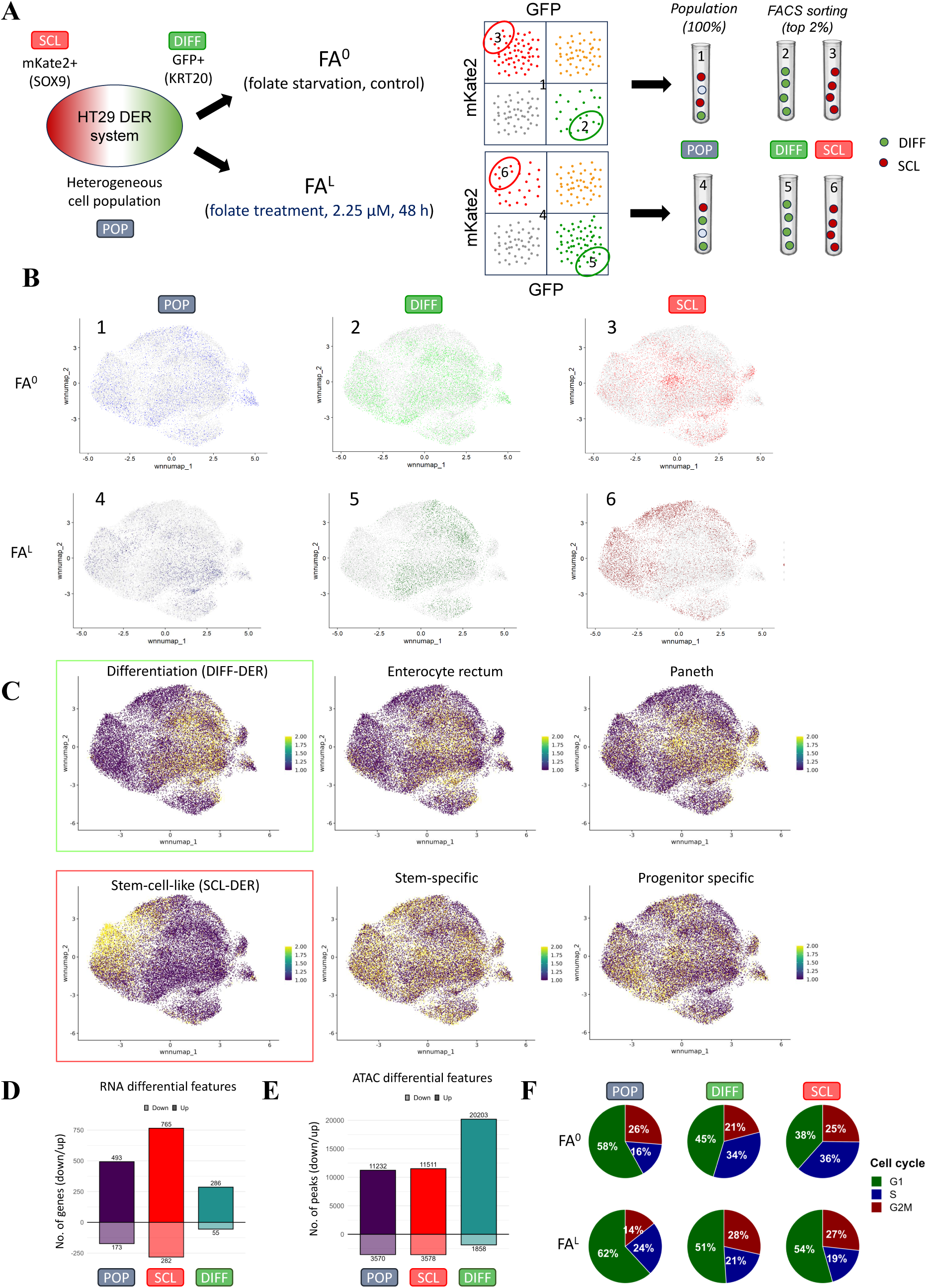
Folate reorganizes epithelial state composition through coordinated chromatin and transcriptional remodeling. **A**, Experimental design of paired single-cell chromatin accessibility and transcriptome profiling (10x Multiome) performed on DER-sorted HT29 epithelial populations under FA⁰ (folate starvation) and FAᴸ (folate supplementation, 2.25 μM, 48 h) conditions. The unsorted population (POP) and the top 2% mKate2⁺ (SCL) and GFP⁺ (DIFF) fractions were isolated by FACS, yielding six single-cell multiome datasets (1–6). **B,** Integrated multiome UMAP embedding of the six datasets, shown separately for POP, SCL and DIFF fractions under FA⁰ (panels 1–3) and FAᴸ (panels 4–6) conditions, with panel numbering corresponding to (A). Reporter-defined SCL and DIFF fractions preferentially occupied distinct regions of the epithelial manifold, whereas unsorted POP cells showed a broader and more heterogeneous distribution. **C,** Projection of lineage-associated transcriptional programs onto the integrated epithelial manifold. DER-derived differentiation (DIFF-DER (RNA)) and stem-cell-like (SCL-DER (RNA)) signatures (boxed) are shown alongside independent enterocyte, Paneth, stem-specific, and progenitor-specific gene programs. Differentiation-associated signatures localize predominantly to DIFF regions, whereas stem-like programs localize to SCL regions, consistent with a graded stem-like-to-differentiated organization of the epithelial manifold. Lineage-specific gene sets and data sources are summarized in **Tables S2C, S7C**. **D,** Differential gene expression analysis between FA⁰ and FAᴸ conditions within POP, SCL, and DIFF populations. Numbers indicate significantly upregulated (above the axis) and downregulated (below the axis) genes (|log_2_FC| > 1, adjusted *p* < 0.05). The most extensive folate-responsive transcriptional remodeling occurs within the stem-like population. **E,** Differential chromatin accessibility analysis between FA⁰ and FAᴸ conditions within POP, SCL, and DIFF populations. Numbers indicate significantly more accessible (above the axis) and less accessible (below the axis) peaks (|log_2_FC| > 1, adjusted *p* < 0.05). In contrast to the transcriptional response in (D), the differentiated population exhibits the most extensive folate-responsive chromatin accessibility remodeling. **F,** Cell-cycle phase distribution inferred from single-cell transcriptomes across POP, SCL, and DIFF populations under FA⁰ and FAᴸ conditions. Cell-cycle phases were assigned using Seurat CellCycleScoring. Folate supplementation increases the G1 fraction in all three populations. S-phase occupancy decreases in the sorted DIFF and SCL states and increases in the unsorted POP fraction, whereas the G2M fraction increases in DIFF and SCL and decreases in POP. The largest redistribution is observed in the stem-like population.

Integration of the six fractions resolved a continuous epithelial manifold organized between SOX9-associated stem-like and KRT20-associated differentiated states (**Fig. 2B; Table S2B**). Endogenous SOX9 and KRT20 expression and the previously defined SCL-DER (RNA) and DIFF-DER (RNA) signatures (30) mapped concordantly along this axis (**Fig. 2C; Fig. S2D,E**). The same reciprocal pattern was apparent across the sorted fractions, with POP cells occupying heterogeneous and intermediate positions and the folate-treated DIFF and SCL fractions showing the strongest enrichment for their corresponding programs (**Fig. S2F,G**). Curated intestinal stem, progenitor, enterocyte and Paneth-like programs localized concordantly along the same axis, supporting its interpretation as a continuous stem-like-to-differentiated organization (**Fig. S2H; Table S2C**).

We independently defined chromatin counterparts of the DER states by bulk ATAC-seq of top-2% mKate2⁺ and GFP⁺ fractions, identifying 1,643 SCL-enriched and 748 DIFF-enriched accessible regions (**Fig. S2I**). Projection of the independently derived ATAC signatures across the sorted fractions recapitulated the reciprocal state organization. DIFF cells were preferentially enriched for the DIFF-DER (ATAC) program, whereas SCL cells were preferentially enriched for the SCL-DER (ATAC) program and unsorted POP cells occupied heterogeneous and intermediate positions (**Fig. S2J**). The transcriptional and accessibility signatures therefore resolved the same stem-like-to-differentiated epithelial axis, showing that this state organization is reflected in both gene expression and chromatin accessibility.

### Folate induces state-specific transcriptional and chromatin remodeling

Folate availability preserved the overall organization of the epithelial manifold but altered where the sorted fractions were positioned within it. Under FA⁰, DIFF cells extended into manifold regions occupied by *SOX9* expression and stem- and progenitor-associated programs, whereas under FAᴸ they concentrated within the differentiated domain (**Fig. 2B**; **Fig. S2D,H**). SCL cells under FA⁰ likewise occupied broader, intermediate territory rather than concentrating at the stem-like pole. Folate deprivation therefore blurred the positional separation of both poles, and the consequence differed according to the starting epithelial state.

Within-state comparisons confirmed this asymmetry. SCL cells showed the largest transcriptional response to folate restoration, with 765 upregulated and 282 downregulated genes, compared with 493 and 173 in POP and 286 and 55 in DIFF cells (**Fig. 2D**). Chromatin remodeling followed a different hierarchy, with DIFF cells showing the greatest number of differentially accessible regions (20,203 gained and 1,858 lost), followed by SCL (11,511 gained and 3,578 lost) and POP cells (11,232 gained and 3,570 lost) (**Fig. 2E**; **Table S2D,E**). The limited overlap of responsive genes and regions among fractions further indicated that folate engages state-specific regulatory programs rather than a uniform response (**Fig. S2K–O**).

Finally, cell-cycle phase distributions inferred from scRNA-seq changed in a cell-state-dependent manner (**Fig. 2F**). Under folate restriction, both sorted states showed high S-phase occupancy (DIFF, 34%; SCL, 36%), which fell after folate restoration (21% and 19%, respectively). The displaced cells were redistributed differently, toward both G1 and G2/M in DIFF cells but predominantly toward G1 in SCL cells. Because S-phase occupancy in a static snapshot reflects both phase entry and duration, these changes do not by themselves distinguish altered cell-cycle progression from altered phase-transition rates. The heterogeneous POP fraction responded in the opposite direction, gaining S-phase and losing G2/M representation (16% to 24% and 26% to 14%, respectively). Despite the reduced S-phase occupancy of the sorted SCL and DIFF states, folate supplementation increased population-level HT29 cell density approximately twofold over five days (**Fig. S2P**), indicating that snapshot cell-cycle distributions did not translate directly into population accumulation. Notably, SCL cells displayed their highest S-phase occupancy under folate restriction, precisely the condition under which they were least fluoropyrimidine-sensitive (**Fig. 1I**), demonstrating that replication-associated cell-cycle state can be uncoupled from drug response.

Together, these findings establish the DER SCL–DIFF axis as a continuous, multimodally defined epithelial-state system and show that folate restoration remodels its constituent states through distinct transcriptional, chromatin and cell-cycle responses. The strongest transcriptional response occurred in SCL cells, whereas the largest accessibility response occurred in DIFF cells, indicating that folate-dependent remodeling is shaped by the epithelial state in which it begins.

### Folate restoration drives a progressive stem-like-to-differentiated transition

The endpoint comparison in **Fig. 2** showed that folate remodels each epithelial state differently, but it could not resolve how these transitions unfold over time or through which intermediate states. Because the earliest regulatory events underlying cell-state transitions are likely to precede establishment of a new steady state, we captured this response window at high temporal resolution by performing single-cell multiome profiling across ten time points (0, 1, 2, 3, 4, 6, 12, 24, 36 and 48 h) following folate addition and reconstructing a pseudotemporal trajectory of the response (**Fig. 3A**).

**Figure 3.**
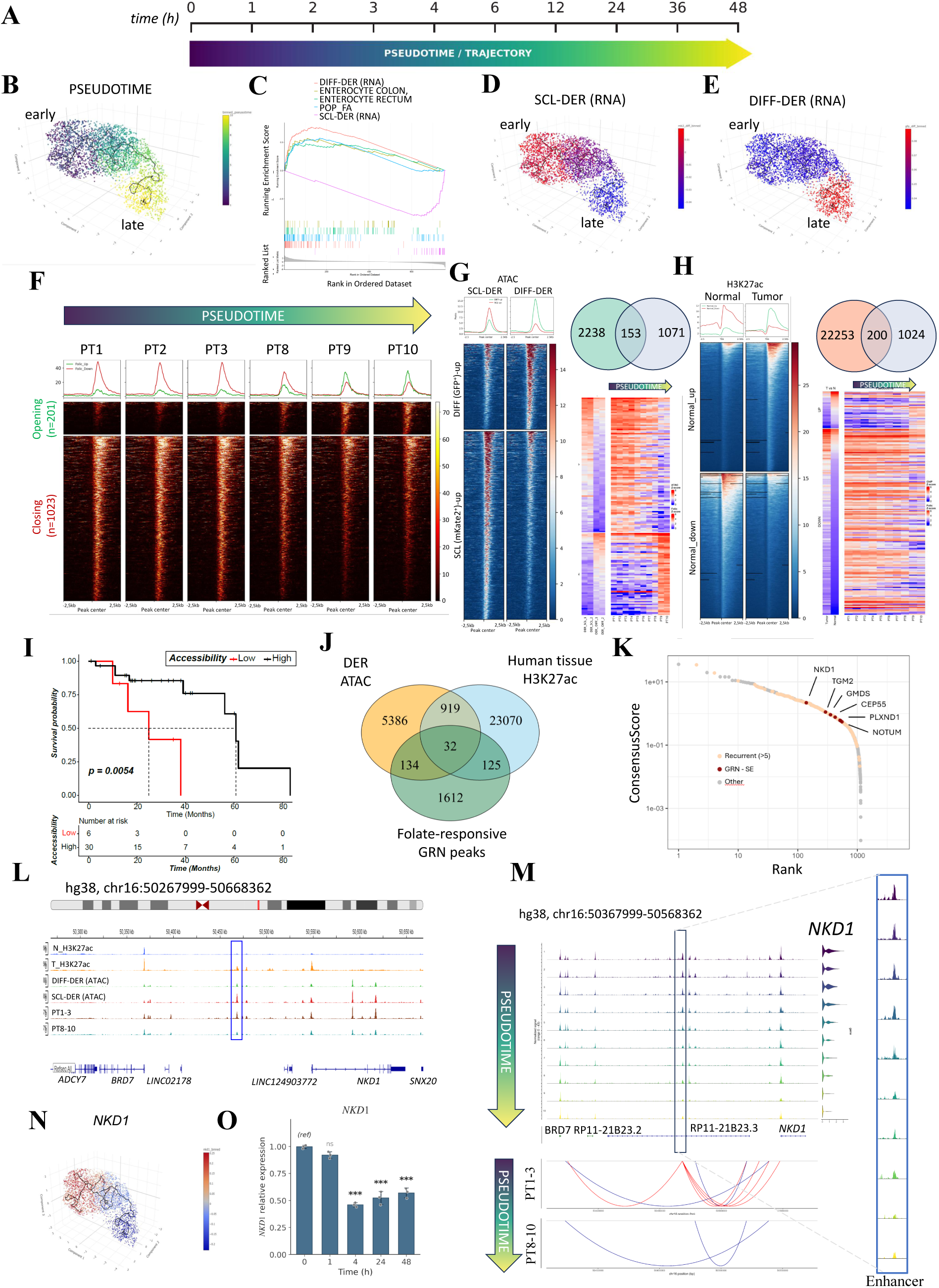
Folate drives chromatin remodeling along the epithelial state trajectory. **A**, Experimental design of time-resolved paired single-cell chromatin-accessibility and transcriptome profiling using the 10x Genomics Multiome ATAC + Gene Expression platform across ten time points following folate supplementation (0, 1, 2, 3, 4, 6, 12, 24, 36 and 48 h). **B,** Three-dimensional UMAP embedding of the pooled ten-time-point dataset, colored by inferred pseudotime. Cells form a continuous trajectory extending from early stem-like states toward differentiated endpoints. **C,** Gene set enrichment analysis of transcriptomes ranked along pseudotime. DIFF-DER (RNA) and SCL-DER (RNA) signatures are shown together with independent enterocyte colon, enterocyte rectum and folate-treated unsorted population (POP_FA) gene programs. Differentiation- and enterocyte-associated signatures are enriched at late pseudotime, whereas stem-like programs are enriched at early pseudotime. Gene sets and data sources are summarized in **Tables S2C, S7C.** **D**, Projection of the SCL-DER (RNA) signature onto the trajectory embedding. High signature scores localize predominantly to early pseudotime states. **E,** Projection of the DIFF-DER (RNA) signature onto the trajectory embedding. High signature scores localize predominantly to late pseudotime states. **F,** Heatmaps of chromatin accessibility across pseudotime bins (PT1–PT10) at 201 trajectory-opening and 1,023 trajectory-closing regulatory regions, centered on peak summits (±2.5 kb). Average accessibility profiles are shown above each heatmap. **G,** Comparison of trajectory-associated differential ATAC peaks with DER ATAC peaks from SCL and DIFF populations. The Venn diagram indicates shared regulatory regions, and heatmaps show chromatin accessibility across the overlapping peak sets. **H,** Comparison of trajectory-associated differential ATAC peaks with human colorectal tissue H3K27ac regions. The Venn diagram indicates shared regulatory regions, and heatmaps display H3K27ac signal across trajectory-opening and trajectory-closing regulatory elements in normal colon and colorectal tumor tissue. Human colorectal tissue H3K27ac data were obtained from GSE166254 (32); see **Table S7C**. **I,** Kaplan–Meier overall survival analysis of TCGA-COAD patients with available ATAC-seq profiles (*n* = 36). Samples were partitioned into low and high accessibility groups by k-means clustering of accessibility at COAD-specific trajectory-opening regulatory regions, yielding 6 and 30 samples, respectively (**Fig. S3B**). Statistical significance was determined by log-rank test. **J,** Overlap among regulatory regions identified by DER ATAC-seq, differential H3K27ac profiling of matched colorectal tumor and normal tissues, and the GRN-associated regulatory regions inferred from the folate time-course multiome data. The DER ATAC set comprises the union of the SCL-DER (ATAC) and DIFF-DER (ATAC) peaks. Thirty-two regulatory regions were shared across all three datasets. **K,** Ranking of candidate regulatory loci across the integrated regulatory analyses. Loci are ordered by the combined evidence from dataset recurrence, standardized signal across datasets, and super-enhancer annotation. *NKD1* and *TGM2* are indicated. **L,** Genome browser view of the *NKD1* locus showing H3K27ac profiles in normal colon and colorectal tumor tissue (as in H), DER ATAC profiles from DIFF and SCL populations, and multiome ATAC profiles representing early and late pseudotime states. *NKD1* enhancer position marked by blue rectangle. **M,** Pseudotime-resolved chromatin accessibility and inferred peak-to-gene co-accessibility at the *NKD1* locus, showing changes in putative enhancer–promoter connectivity across the epithelial state transition. **N,** Projection of *NKD1* expression onto the trajectory embedding. **O,** qRT-PCR analysis of *NKD1* expression following folate supplementation in HT29 cells at the indicated time points. Expression was normalized to *ACTB* and expressed as fold change relative to the mean of the 0 h control replicates. Data are presented as mean ± SD (n = 3 biological replicates). Statistical significance was determined by two-sided Student’s t-test versus 0 h. ns, not significant; \*\*\**p* < 0.001.

The reconstructed manifold revealed a continuous cellular trajectory extending from early stem-like states toward differentiated endpoints (**Fig. 3B**). Gene set enrichment analysis of single-cell transcriptomes ranked along pseudotime confirmed this organization, with differentiation- and enterocyte-associated programs enriched positively along the trajectory and the stem-like program enriched negatively (**Fig. 3C**). Stemness scores based on mKate2⁺-associated gene sets (30) localized to early pseudotime states, whereas differentiation-associated signatures derived from GFP⁺ gene sets localized to the differentiated endpoint, indicating progressive loss of stem-like identity under folate exposure (**Fig. 3D,E**).

### Trajectory-resolved chromatin remodeling accompanies the folate-driven state transition

To resolve the chromatin-level changes underlying this transition, we aligned ATAC accessibility profiles along pseudotime (**Fig. 3F**; **Table S3A**). Aggregate profiles revealed two opposing patterns, with regions accessible in early pseudotime progressively closing along the trajectory and initially inaccessible regions gradually opening toward differentiated states. In total, 1,224 regulatory regions displayed significant pseudotime-associated dynamics, 1,023 of which progressively closed and 201 of which gained accessibility. This reconfiguration occurred gradually across pseudotime rather than as a discrete switch, consistent with continuous regulatory remodeling accompanying folate-induced differentiation.

To determine whether these dynamics correspond to state-defining regulatory programs, we compared trajectory differential ATAC peaks with the SCL-DER (ATAC) and DIFF-DER (ATAC) signatures derived from bulk ATAC-seq of FACS-sorted stem-like (SCL, mKate2⁺) and differentiated (DIFF, GFP⁺) populations (**Fig. S3A**). Regions closing along pseudotime overlapped preferentially with mKate2⁺-enriched peaks, whereas regions gaining accessibility corresponded to sites preferentially accessible in GFP⁺ cells (**Fig. 3G**), demonstrating that pseudotime-resolved chromatin remodeling recapitulates the regulatory differences between DER-defined states.

To place these changes into a clinical epigenomic context, we intersected trajectory differential regions with publicly available colorectal cancer tissue H3K27ac ChIP–seq datasets from normal colon and tumor samples (31,32). A subset of dynamically regulated loci overlapped with enhancer regions displaying differential H3K27ac activity in colorectal cancer tissues (**Fig. 3H**), indicating that regions remodeled during folate-induced differentiation correspond to regulatory elements active in human colorectal cancer epigenomes.

Because the 201 progressively opening regions represented elements selectively activated during folate-induced differentiation, we asked whether accessibility at these loci was associated with clinical outcome. Using publicly available TCGA-COAD ATAC-seq profiles from 36 patients (33), reduced accessibility at trajectory-opening regions was associated with significantly worse overall survival (**Fig. 3I**). The same analysis using all overlapping trajectory-opening regions, without COAD-specific filtering, showed the same direction of association (**Fig. S3B,C**), showing that accessibility at folate-responsive regulatory regions is associated with clinical outcome.

### Recurrent folate-responsive regulatory loci emerge during stem-like state destabilization and differentiation-associated remodeling

To identify regulatory loci associated with folate-responsive remodeling, we intersected DER ATAC peaks, differential tissue H3K27ac regions and GRN-associated regulatory regions. This analysis identified 32 loci shared across all three datasets (**Fig. 3J**; **Table S3B,C**). Additional comparisons with public Cistrome datasets are shown in **Fig. S3D,E**.

Among the recurrent folate-responsive loci, NKD1 and TGM2 emerged as the strongest candidates across the integrated analyses (**Fig. 3K**; **Table S3D**). Subsequent analyses focused on NKD1, the highest-ranked locus, while TGM2 was retained as an additional example of the broader regulatory remodeling observed across multiple loci (**Fig. S3F**).

Integrated analysis of tissue H3K27ac, DER ATAC and trajectory-resolved accessibility profiles demonstrated coordinated chromatin remodeling at the NKD1 locus across stem-like and differentiated states (**Fig. 3L**), and peak-to-gene co-accessibility analysis revealed marked differences in putative enhancer–promoter interactions between early and late pseudotime (**Fig. 3M**). NKD1 expression localized predominantly to early pseudotime stem-like states and progressively declined along the trajectory (**Fig. 3N**), which qRT–PCR independently confirmed following folate exposure in HT29 cells (**Fig. 3O**). Together with the enhancer accessibility dynamics, these findings support a model in which high NKD1 expression is a feature of a stem-associated regulatory program that is progressively suppressed upon folate restoration.

Because *NKD1* emerged as the most consistently folate-responsive locus, we first asked how sensitively it responds to folate availability itself. Graded folate supplementation elicited coordinated changes in *NKD1* and *TGM2* expression across a broad concentration range, with marked responses already within the physiological range of human plasma folate concentrations (**Fig. S3G**). We next asked whether *NKD1* also responds to therapeutic perturbations that shape fluoropyrimidine response (**Fig. 1F–M**). 5-FU induced *NKD1* expression in a dose- and time-dependent manner, with the strongest induction after prolonged exposure at the highest concentration tested (**Fig. S3H**). Under folate-deprived conditions, the largest observed MTX-induced increase in *NKD1* expression occurred at 0.03 μM after 24 h. Under FAᴸ conditions, no induction was observed at 0.03 μM, although significant induction remained detectable at 0.1 μM (**Fig. S3I,J**). Folate deprivation and MTX exposure under FA⁰ conditions, both of which increased *NKD1* expression, were also associated with reduced 5-FU sensitivity and increased post-treatment persistence (**Fig. 1F–M**). Separately, the induction of *NKD1* by 5-FU is consistent with previous reports linking fluoropyrimidine exposure to WNT-dependent stem-like persistence programs in colorectal cancer (34). These findings identify *NKD1* as a regulatory locus responsive both to physiological variation in folate availability and to clinically used antimetabolites, placing it at the intersection of folate-responsive epithelial remodeling and treatment response.

### Folate rewires transcription factor-associated regulatory programs during epithelial state transition

Building on the trajectory-resolved chromatin remodeling identified in **Fig. 3**, we next asked which transcription factor-associated regulatory programs accompanied this state transition. We integrated three complementary regulatory readouts along pseudotime, chromVAR-based accessibility of TF-binding motifs, expression of the corresponding transcription factors, and expression of their associated target-gene programs. chromVAR analysis revealed a shift in TF-motif accessibility rather than uniform suppression of chromatin accessibility (**Fig. 4A; Fig. S4A**). AP-1-associated motifs progressively lost accessibility across pseudotime, whereas HNF1-, SP- and KLF-associated motifs became increasingly accessible.

**Figure 4.**
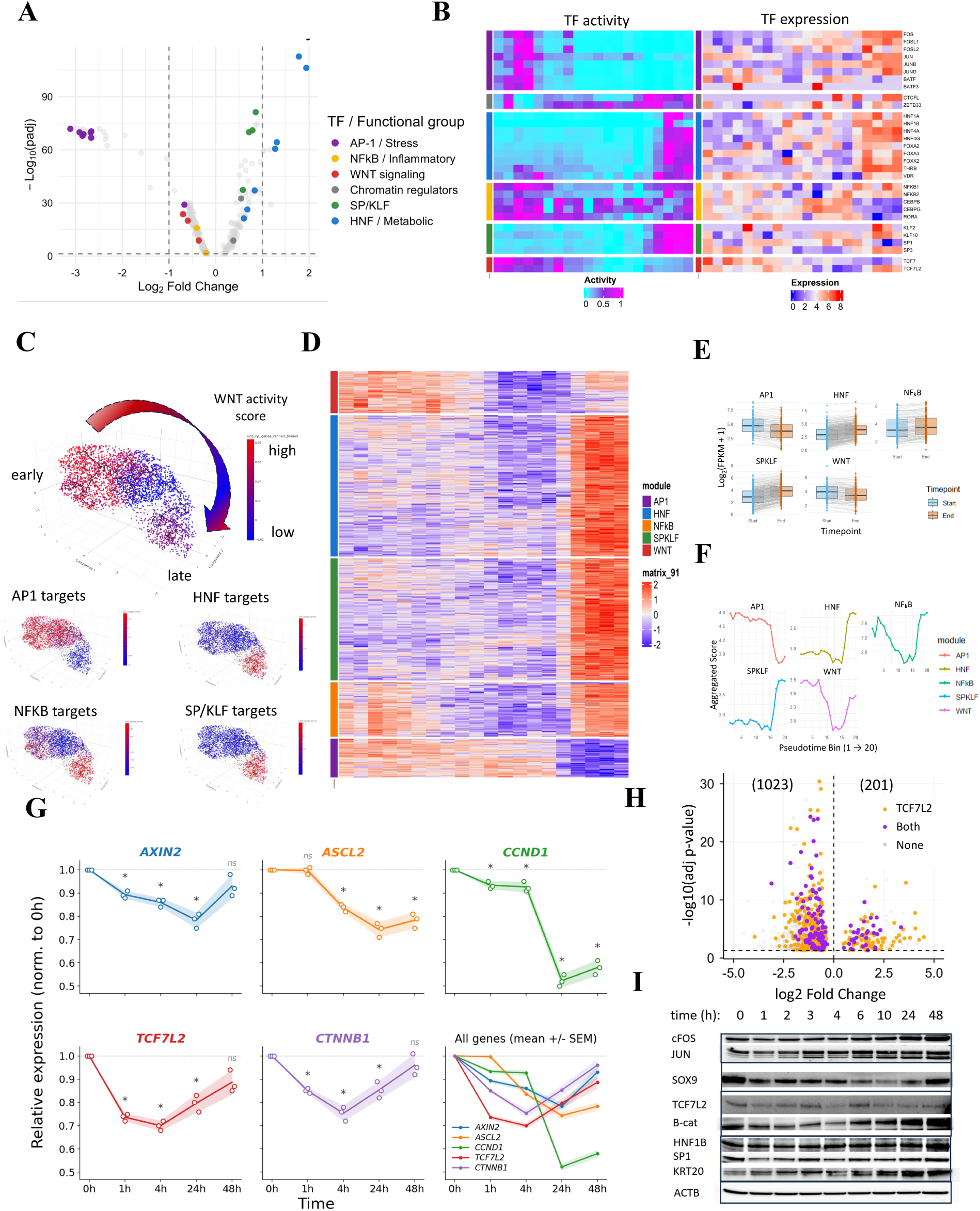
Folate rewires transcription factor activity and attenuates WNT signaling during epithelial state transition. **A**, Differential transcription factor motif activity inferred by chromVAR between late and early pseudotime states. Transcription factors are colored according to functional groups, including AP-1/stress, NF-κB/inflammatory, WNT signaling, chromatin regulators, SP/KLF, and HNF/metabolic regulators. The same color scheme is used in panels A and B. **B,** Heatmaps showing transcription factor motif activity (left) and corresponding transcription factor expression (right) across pseudotime. Transcription factors are grouped according to the functional classes shown in (A). **C,** Projection of WNT activity score (top) and AP-1, HNF, NF-κB, and SP/KLF target gene module scores (bottom) onto the pseudotime trajectory. WNT activity is highest at early pseudotime and progressively decreases along the trajectory. **D,** Heatmap of z-scored gene expression for genes assigned to the AP-1, HNF, NF-κB, SP/KLF, and WNT regulatory modules, ordered along pseudotime. Columns correspond to pseudotime bins (*n* = 20); module assignment is indicated by the color bar on the left. **E,** Comparison of regulatory module expression between early (pseudotime bins 1–4) and late (bins 17–20) trajectory states for the AP-1, HNF, NF-κB, SP/KLF, and WNT programs. Grey lines connect individual genes; boxplots show the distribution across each module. **F,** Aggregated regulatory module scores across the full pseudotime trajectory (bins 1–20) for the AP-1, HNF, NF-κB, SP/KLF, and WNT programs. **G,** qRT–PCR of representative WNT pathway genes in HT29 cells following folate supplementation. Expression was normalized to *ACTB* and is shown as fold change relative to the 0h baseline of each biological replicate. Circles, individual biological replicates (*n* = 3); lines, mean; shaded bands, SEM. Bottom-right panel, all genes overlaid (mean ± SEM). Statistical significance was determined using a two-sided one-sample *t*-test on log_2_-transformed fold changes, followed by Benjamini–Hochberg correction. \**q* < 0.05; ns, not significant. **H,** Overlap of folate-responsive trajectory-differential regions with TCF7L2 and β-catenin occupancy. The 1,224 trajectory-associated differentially accessible regions identified in Fig. 3F (1,023 trajectory-closing and 201 trajectory-opening) were intersected with our TCF7L2 and β-catenin ChIP-seq data and are plotted by log_2_ fold change between late and early pseudotime states against −log₁₀(adjusted p value). Orange, regions overlapping TCF7L2 only; purple, regions overlapping both TCF7L2 and β-catenin; grey, regions overlapping neither. The horizontal dashed line marks the significance threshold, and the vertical dashed line indicates zero log_2_ fold change. Related analyses using public and in-house ChIP-seq datasets are shown in **Fig. S4C,D**, respectively. **I,** Time-course immunoblot analysis of AP-1 components (cFOS, JUN), WNT-associated regulators (TCF7L2, β-catenin), stem- and differentiation-associated markers (SOX9, KRT20), and HNF1B and SP1 following folate supplementation in HT29 cells. ACTB served as a loading control.

Comparison of chromVAR-inferred motif activity with TF transcript abundance showed that changes in motif activity were not simply explained by corresponding changes in TF expression. AP-1-associated motif accessibility declined despite maintained or increased expression of several AP-1 family members, revealing a divergence between TF expression and the accessibility of their cognate motifs during folate-induced differentiation (35–37) (**Fig. 4B; Fig. S4B; Table S4A,B**).

Integration with target-gene expression further showed that these changes were accompanied by coordinated remodeling of downstream regulatory programs. The dominant directional change was attenuation of WNT signaling. Regulatory module activities projected onto the trajectory embedding showed progressive loss of WNT activity together with redistribution of AP-1-, NF-κB-, HNF- and SP/KLF-associated programs (**Fig. 4C**). Trajectory-resolved expression analysis showed coordinated suppression of WNT target genes and remodeling of the corresponding TF-associated target-gene programs (**Fig. 4D; Table S4C**). This pattern was evident both in direct early-versus-late comparisons and across the aggregated pathway dynamics along pseudotime (**Fig. 4E,F**). At the transcriptional output level, qRT–PCR independently confirmed progressive downregulation of canonical WNT target genes across the folate treatment time course (**Fig. 4G**).

### WNT effectors lose chromatin occupancy upon folate supplementation

The attenuation of WNT-associated regulatory activity along the trajectory prompted us to investigate WNT-effector occupancy at folate-responsive regulatory elements. CistromeDB analysis identified significant enrichment for public TCF7L2 and β-catenin ChIP-seq datasets at trajectory-associated differential peaks (38) (**Fig. S4C**). To determine their occupancy in HT29 cells under defined folate conditions, we performed TCF7L2 and β-catenin ChIP-seq following folate deprivation and supplementation. Integration with the trajectory-associated peaks revealed regions occupied by TCF7L2 alone or by both factors (**Fig. 4H**).

Within trajectory-opening peaks, folate supplementation significantly reduced both TCF7L2 and β-catenin occupancy despite concomitant increases in chromatin accessibility and H3K27ac enrichment, whereas trajectory-closing peaks showed less coordinated changes (**Fig. S4D**). Thus, increased chromatin accessibility was accompanied by reduced TCF7L2 and β-catenin occupancy at these regions.

Immunofluorescence and single-target proximity ligation assays showed that β-catenin redistributed from the nucleus toward the cell periphery following folate supplementation (**Fig. S4E,F**). Dual-target proximity ligation assays also showed reduced nuclear proximity between β-catenin and TCF7L2 (**Fig. S4G**). Although total β-catenin protein levels increased (**Fig. 4I**), its nuclear localization, proximity to TCF7L2 and chromatin occupancy decreased. These findings indicate that folate supplementation attenuates canonical WNT signaling through changes in β-catenin localization and regulatory engagement rather than reduced protein abundance. Immunoblot analysis further supported the broader regulatory reorganization observed along the trajectory (**Fig. 4I**). AP-1 components, particularly c-FOS and JUN, were maintained or increased despite reduced AP-1-associated motif accessibility, whereas TCF7L2 and SOX9 decreased and HNF1B and SP1 increased, consistent with the broader shift toward differentiation-associated transcription factor programs (39).

### Folate-pathway metabolites converge on fluoropyrimidine sensitization

To dissect how folate availability is coupled to chromatin state and chemotherapy response, we first mapped the interconnected folate and methionine-cycle pathways, in which folate-derived one-carbon units feed methionine and subsequently SAM biosynthesis, positioning folate availability upstream of methylation potential (**Fig. 5A**). Because folate-cycle intermediates contribute to both nucleotide biosynthesis and methyl-donor production, we asked whether methyl-donor availability contributes to the chemosensitizing effect of folate.

**Figure 5.**
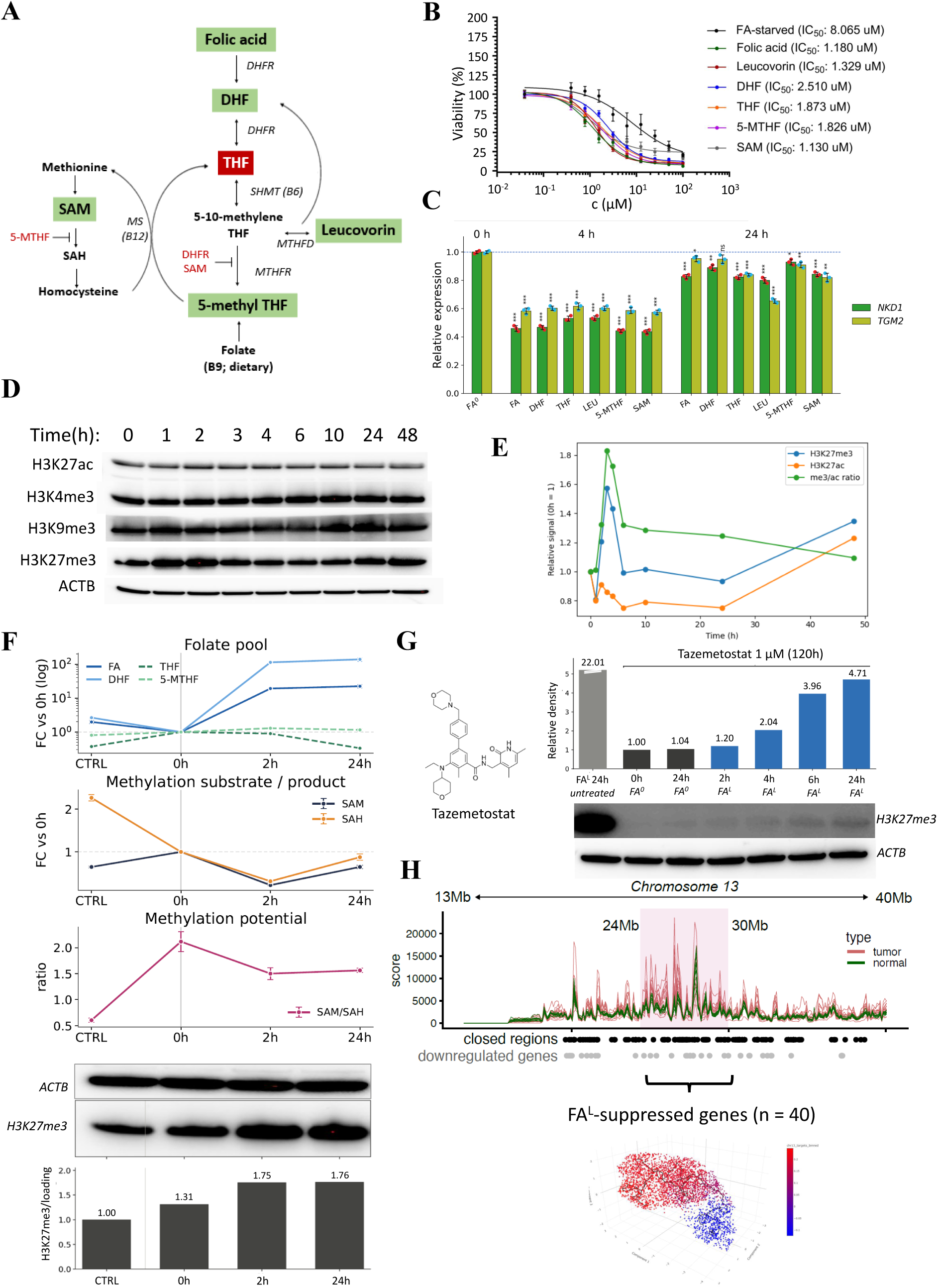
One-carbon metabolism supports fluoropyrimidine sensitization and EZH2-dependent H3K27me3 remodeling. **A**, Schematic of folate and methionine-cycle metabolism, highlighting folate intermediates (FA, DHF, THF, 5-MTHF), one-carbon transfer reactions, and SAM-dependent methylation. **B,** 5-FU dose–response in HT29 cells supplemented with folate-pathway metabolites (folic acid, leucovorin, DHF, THF, 5-MTHF) or SAM, versus folate-starved control (FA⁰). IC_50_ values were determined by nonlinear regression using a four-parameter logistic model from three biological replicates. No statistical comparison between conditions was performed. **C,** *NKD1* and *TGM2* expression at 0, 4 and 24 h following supplementation with folate-pathway metabolites under FA⁰ conditions. Data are presented as mean ± SD (*n* = 3 biological replicates). Statistical significance was determined by two-sided Student’s *t*-test against the 0 h FA⁰ control. \**p* < 0.05, \*\**p* < 0.01, \*\*\**p* < 0.001; ns, not significant. **D,** Time-course immunoblot of H3K27ac, H3K4me3, H3K9me3, and H3K27me3 in HT29 cells following folic acid supplementation (0–48 h). ACTB, loading control. **E,** Densitometric quantification of H3K27me3 and H3K27ac signals from Fig. 5D, and the H3K27me3/H3K27ac ratio over time, normalized to *t* = 0. A representative experiment of three independent repeats is shown. **F,** Targeted metabolomic profiling of folate intermediates and methylation-cycle metabolites in HT29 cells following folic acid treatment (CTRL, 0 h, 2 h, 24 h). Metabolite levels were normalized to cell number and are presented as mean ± SD (*n* = 3 biological replicates). Top, intracellular folate pools (FA, DHF, THF, 5-MTHF). Middle, SAM and SAH levels. Bottom, SAM/SAH ratio. Lower panels, parallel H3K27me3 immunoblot and densitometric quantification at matched time points, showing a representative experiment of three independent repeats. **G,** Folate-dependent recovery of H3K27me3 after EZH2 inhibition. HT29 cells were treated with 1 μM tazemetostat for 120 h, followed by inhibitor washout and recovery in folate-free (FA⁰) or folate-containing (FAᴸ) medium for the indicated times. Immunoblots and corresponding densitometry from one representative experiment of three independent biological experiments are shown. H3K27me3 signals were normalized to ACTB and expressed relative to the FA⁰ sample collected immediately after washout (0 h). The untreated FAᴸ reference showed a substantially higher, saturated H3K27me3 signal and is included for qualitative comparison only. No statistical comparisons were performed. **H,** Integration of folate-responsive chromatin remodeling with primary tumor H3K27me3 domains. Tumor-enriched H3K27me3 regions across chromosome 13 were aligned with folate-responsive closed-ATAC regions and downregulated genes (n=40) following folate restoration, and projected onto the epithelial trajectory shown in Fig. 3B.

We therefore supplemented folate-starved HT29 cells with individual folate-pathway intermediates (folic acid, leucovorin, DHF, THF, 5-MTHF) or with SAM directly, and measured 5-FU dose– response. All tested metabolites substantially reduced 5-FU IC_50_ values, from 8.1 μM under FA⁰ to 1.1–2.5 μM, corresponding to approximately 3- to 7-fold sensitization comparable in magnitude to that observed in parental HT29 cells and across reporter-defined epithelial fractions (**Fig. 5B**; **Fig. 1F,I**). Structurally and functionally distinct folate-pathway metabolites therefore produced convergent effects, while the ability of SAM itself to reproduce the sensitizing response specifically implicated methyl-donor availability as a shared component of folate-dependent fluoropyrimidine sensitization. The same metabolites also induced rapid and largely indistinguishable downregulation of *NKD1* and *TGM2* within 4 h, converging by 24 h (**Fig. 5C**), showing that the convergent metabolic response propagates to a coherent transcriptional output irrespective of which intermediate was supplied.

### Folate restoration increases methylation potential and supports EZH2-dependent H3K27me3 deposition

The ability of SAM to reproduce the sensitizing effect of folate-pathway metabolites prompted us to ask whether folate restoration is accompanied by changes in cellular methylation potential and chromatin methylation. Time-course immunoblotting following folic acid supplementation revealed dynamic changes across multiple histone marks (**Fig. 5D**), among which the Polycomb-associated repressive mark H3K27me3 showed the most pronounced and temporally structured response. Densitometry confirmed a rapid and transient increase in H3K27me3 within the first hours of folate restoration, accompanied by a modest concurrent decrease in H3K27ac and therefore an early elevation of the H3K27me3/H3K27ac ratio (**Fig. 5E**). Folate restoration thus does not simply increase repressive methylation, but transiently rebalances active and repressive chromatin states.

To link these chromatin dynamics to the underlying metabolic changes, we performed targeted metabolomic profiling of folate intermediates and methylation-cycle metabolites following folic acid treatment (**Fig. 5F; Fig. S5A–F**; **Table S5A**). Intracellular folic acid and DHF accumulated rapidly, consistent with active uptake and entry into the folate cycle. SAM and SAH levels transiently decreased at early time points, whereas the SAM/SAH ratio, an established indicator of cellular methylation potential, increased upon folate restoration and remained elevated through 24 h. This elevated methylation potential was paralleled by progressive H3K27me3 deposition at matched time points, showing that increased cellular methylation potential was temporally associated with H3K27me3 accumulation (**Fig. 5F**).

Correlation alone, however, does not establish whether folate availability is required for efficient H3K27me3 re-deposition. To test this directly, we depleted H3K27me3 with the EZH2 inhibitor tazemetostat (1 μM, 120 h), then washed out the inhibitor and allowed recovery in either folate-free or folate-containing medium. No detectable H3K27me3 recovery occurred within 24 h after inhibitor washout under FA⁰ conditions, whereas transfer to FAᴸ medium triggered progressive re-deposition that became apparent from 4 h onwards and increased at 6 and 24 h (**Fig. 5G**). Thus, folate availability supports efficient H3K27me3 re-deposition following release from EZH2 inhibition, consistent with a role for folate-dependent methyl-donor availability in this process. Together with the metabolomic and chromatin time-course data, these findings identify folate-dependent methyl-donor availability as a key metabolic constraint on H3K27me3 re-deposition under these conditions.

### Folate-induced loss of chromatin accessibility overlaps with H3K27me3-enriched domains in primary colorectal tumors

To assess whether this chromatin remodeling is relevant beyond cultured HT29 cells, we asked whether regions that close following folate restoration overlap chromatin domains marked by H3K27me3 in primary colorectal tumors. Integrating tumor H3K27me3-enriched regions with our single-cell multiome accessibility and expression data showed that folate-induced closed-ATAC regions and downregulated genes preferentially overlapped tumor-enriched H3K27me3 domains. A representative domain on chromosome 13 (24–30 Mb) showed concordant loss of chromatin accessibility, transcriptional downregulation and tumor-associated H3K27me3 enrichment, with differential expression of the 40 resident genes shown alongside (**Fig. 5H**; **Fig. S5G**; **Table S5B**). Projection of the closed-ATAC and downregulated-gene signatures onto the folate-responsive epithelial trajectory (**Fig. 3B**) confirmed concordant loss of accessibility and repression along the folate-induced state transition, and these findings extended to chromosome 8 and to independently defined compartment-, oncofetal- and enhancer-state programs (**Fig. S5H,I**). Folate-induced chromatin closure therefore intersects H3K27me3-enriched domains present in primary colorectal tumors, connecting the metabolic response to a tissue-relevant regulatory landscape without implying equivalent global state transitions.

### Folate restoration selectively remodels a polarized SCL–DIFF epithelial axis

To determine whether folate broadly affects colorectal cell-state programs or preferentially remodels a defined epithelial axis, we assembled 33 colorectal-relevant transcriptional programs from published intestinal, tumor-associated, regenerative and developmental datasets (**Table S6**) and quantified their changes across the ten-bin folate-restoration pseudotime trajectory. We used two complementary metrics, the Trajectory Remodeling Index (TRI), which measures the overall magnitude of program change, and the signed Trajectory Directionality Index (TDI), which measures whether that change occurs consistently in one direction. Although many programs changed along the trajectory, the strongest coherent and opposing changes were observed for SCL and DIFF, which shifted in opposite directions (TRI = 2.40 and 2.18; TDI = −0.74 and +0.70, respectively; **Fig. 6A**; **Fig. S6A,B**). Independent intestinal differentiation programs, including enterocyte rectum, absorptive intestine and secretory intestine, moved toward the DIFF pole, whereas endoderm development and injury-repair moved toward the SCL pole. Other programs fluctuated, remained largely stationary or fell below the predefined TRI threshold.

**Figure 6.**
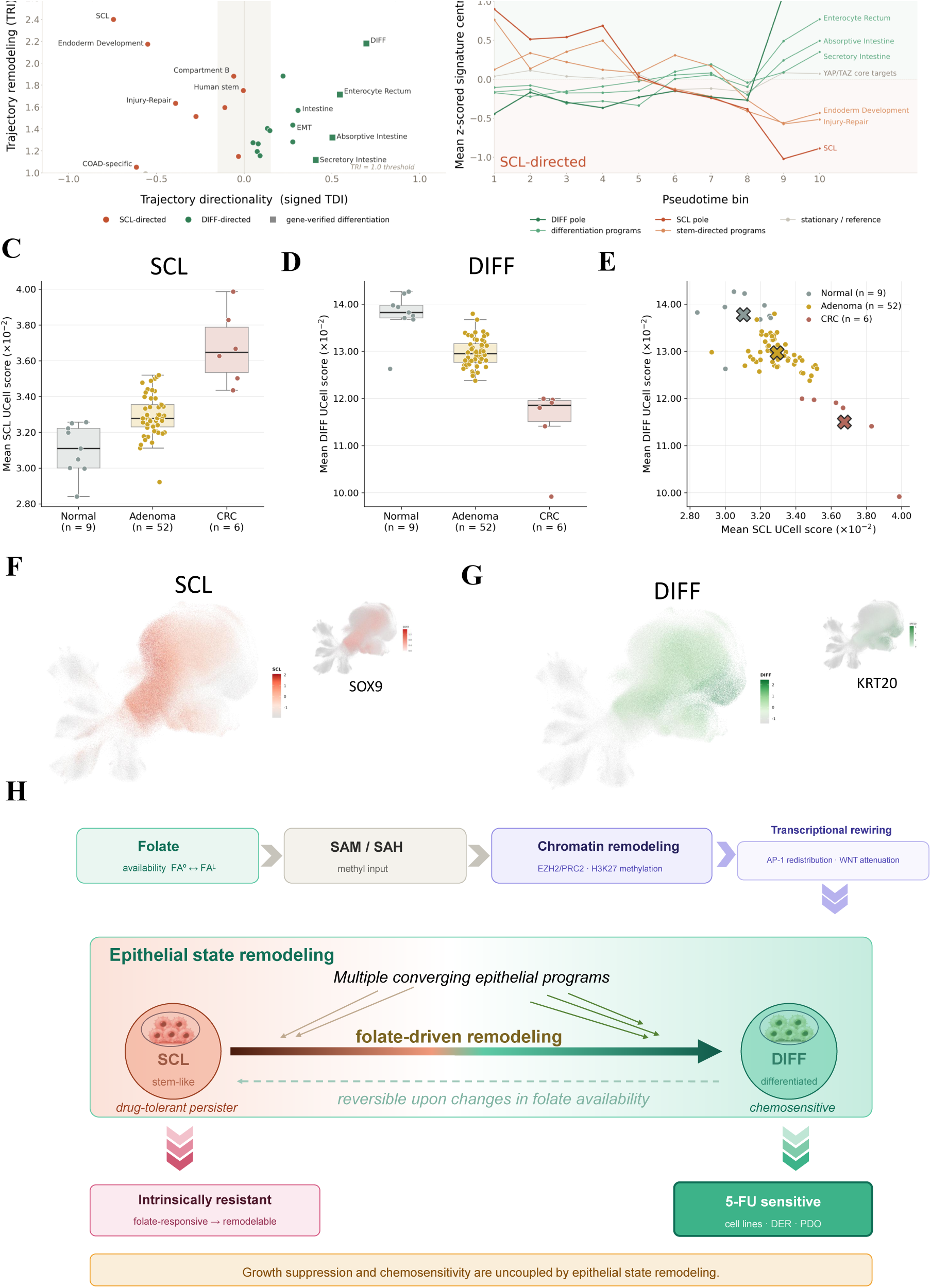
Folate-driven epithelial remodeling is directionally opposed to the SCL–DIFF program shift observed during colorectal tumor progression. **A**, Selective remodeling resolves a polarized SCL–DIFF axis. Each of 33 colorectal-relevant cell-state and tissue-associated programs is positioned according to its Trajectory Remodeling Index (TRI), which quantifies cumulative remodeling magnitude across ten pseudotime bins, and signed Trajectory Directionality Index (TDI), which measures directional coherence. Negative TDI values indicate SCL-directed remodeling (red) and positive values indicate DIFF-directed remodeling (green). The shaded central band marks the non-directional range around zero. The SCL and DIFF programs occupy opposing extremes, combining high TRI with strong directional coherence. The horizontal dashed line indicates the predefined TRI = 1 threshold below which directionality was not interpreted. Squares denote gene-verified differentiation programs, as defined in the Methods; circles denote the remaining programs. **B,** Representative program trajectories across folate-restoration pseudotime. Lines show mean standardized signature-score centroids across ten pseudotime bins. SCL and DIFF undergo reciprocal repositioning, with a pronounced transition between bins 8 and 9. Independent intestinal differentiation programs converge toward the DIFF pole, whereas Endoderm Development and Injury-Repair move toward the SCL pole and YAP/TAZ core targets remain comparatively stationary. **C, D,** SCL (C) and DIFF (D) UCell activity across normal colorectal mucosa, adenoma and colorectal carcinoma. Quantification was restricted to epithelial and malignant epithelial cells and excluded unaffected tissue. Each point represents the mean UCell score of one tissue sample; boxes show the median and interquartile range and whiskers extend to 1.5 times the interquartile range. The analysis included 9 normal, 52 adenoma and 6 carcinoma samples derived from 2, 9 and 6 donors, respectively. Standardized CRC-minus-normal differences are shown within each panel. **E,** Joint SCL–DIFF program space for the samples shown in Fig. 6C and Fig. 6D. Each point represents the paired mean SCL and DIFF UCell scores of one tissue sample and is colored by histological group. Crosses indicate group centroids. Normal mucosa, adenoma and CRC occupy progressively displaced positions toward higher SCL and lower DIFF activity. **F,** Projection of the SCL UCell score and *SOX9* expression onto the human colorectal single-cell atlas. All atlas cells are shown in light grey; program activity and expression are displayed within the main epithelial compartment. Color scales (red) indicate UCell score for SCL and normalized expression for *SOX9*. Unaffected tissue is retained in the embedding, which is shown in full, unlike the filtered quantification in Fig. 6C**–E**. **G,** Projection of the DIFF UCell score and *KRT20* expression onto the same atlas embedding, with color scales (green) indicating UCell score for DIFF and normalized expression for *KRT20*. The spatial correspondence between the program scores and their reporter-associated marker genes supports the stem-like and differentiated identities captured by SCL and DIFF while showing that each module extends beyond a single marker gene. **H,** Conceptual model. Folate availability increases intracellular methylation capacity, reflected by the SAM/SAH ratio, and supports EZH2/PRC2-dependent H3K27me3 deposition. The resulting chromatin remodeling and transcriptional rewiring, including AP-1 redistribution and attenuation of canonical WNT signaling, drive reversible remodeling from SCL epithelial state with lower fluoropyrimidine sensitivity toward DIFF state with increased sensitivity. Multiple independently derived epithelial programs converge on the differentiated pole. Across human normal mucosa, adenoma and carcinoma, the same SCL–DIFF programs show an opposing pattern (Fig. 6C–E), supporting the relevance of this folate-responsive axis to human disease without demonstrating an effect of folate on tumor progression in vivo. Folate availability therefore remodels the epithelial state in which fluoropyrimidines act.

The separation between SCL and DIFF increased across pseudotime, with the largest change occurring between bins 8 and 9 (**Fig. 6B**; **Fig. S6C,D**). Permutation analysis further showed that DIFF remodeling exceeded that observed after randomizing pseudotime order, with none of 1,000 permutations reproducing the observed DIFF TRI (empirical p < 0.001; FDR = 0.00174; **Fig. S6E**). Consistently, stepwise analysis localized the largest DIFF change to the same 8-to-9 transition (**Fig. S6F**).

Two additional analyses supported the reciprocal organization of the SCL–DIFF axis. SCL and DIFF changed in opposite directions across the trajectory, showing an inverse relationship between the two programs (Pearson r = −0.493; Spearman ρ = −0.483; **Fig. S6G–J**). Importantly, the DIFF direction was independently supported by epithelial programs derived outside the DER system, with enterocyte rectum, absorptive intestine and secretory intestine programs shifting toward the DIFF pole, whereas endoderm development and injury-repair programs shifted toward the SCL pole (**Fig. 6A,B**).

### The folate-responsive SCL–DIFF axis is directionally opposed to colorectal tumor progression

We next asked whether the folate-responsive SCL–DIFF axis was also evident in primary human tissue. We scored all 33 programs in the human colorectal single-nucleus atlas of Becker et al. (40), spanning normal mucosa, adenoma and carcinoma (**Fig. S6K**). Within epithelial and malignant epithelial cells, carcinoma showed higher SCL and lower DIFF activity than normal mucosa, with standardized differences of +3.11 and −3.46, respectively (**Fig. 6C,D**). Adenomas occupied intermediate but heterogeneous positions in the joint SCL–DIFF state space (**Fig. 6E**). These were the largest increase and decrease among all 33 programs (**Fig. S6L**). Given the two normal-tissue donors, we report effect sizes without statistical testing.

The spatial distribution of the scores across the atlas embedding supported their biological interpretation. SCL activity overlapped epithelial regions expressing *SOX9*, whereas DIFF activity overlapped regions expressing *KRT20* (**Fig. 6F,G**). Both programs extended beyond their respective marker distributions, indicating that they capture broader transcriptional states rather than individual marker expression.

Across the atlas, the 33 programs formed four correlated groups representing differentiated epithelial states, stem/progenitor and malignant states, non-epithelial populations, and fetal/regenerative/mesenchymal states (**Fig. S6M**). Gene overlap between the programs was generally limited, and the SCL and DIFF signatures shared no genes (**Fig. S6N–Q**). Their opposing activity therefore reflects distinct gene-expression programs rather than overlap between the signatures.

Folate restoration in HT29 cells shifted the SCL–DIFF balance toward the differentiated state, opposite to the pattern observed between normal human mucosa and carcinoma. The same SCL–DIFF axis was evident in human colorectal tissue, with folate restoration shifting these programs in the opposite direction to the normal–carcinoma difference. Together with the greater fluoropyrimidine sensitivity of prospectively isolated DIFF cells (**Fig. 1I,J**), these findings support a model in which folate availability remodels the epithelial-state context in which treatment acts (**Fig. 6H**).

## Discussion

Folate has been part of fluoropyrimidine-based chemotherapy for decades, yet why it improves outcome in some settings and not others has remained unresolved. The canonical explanation is biochemical, whereby reduced folate stabilizes the ternary complex between FdUMP, thymidylate synthase and 5,10-methylenetetrahydrofolate, thereby prolonging target inhibition (1). Our findings do not replace this model but reveal a temporally distinct layer that precedes drug exposure. Folate availability remodels the epithelial state of colorectal cancer cells before treatment. Because prospectively isolated states differ in fluoropyrimidine sensitivity, the metabolic environment preceding therapy establishes the cellular context in which the drug subsequently acts. Folate can therefore influence fluoropyrimidine response both acutely, through target potentiation, and prospectively, through chromatin-dependent epithelial-state remodeling. Critically, this remodeling was reversible within a single genetic background, showing that folate availability can redistribute epithelial-state occupancy without genetic change.

The remodeling resolved here occurs along an epithelial axis distinct from the canonical LGR5-positive stem-cell hierarchy of the normal crypt. Its SCL pole represents an aberrant, APC-dependent, SOX9-associated program rather than the homeostatic stem compartment (14,15,19,21), while neither SCL nor DIFF is reducible to SOX9 or KRT20 expression alone. Endogenous markers, independently derived DER transcriptional and accessibility signatures, curated intestinal programs and the trajectory behavior of 33 cell-state programs all converged on the same continuous organization. Together with their reciprocal movement and the complete absence of gene overlap between the two gene sets, this convergence identifies SCL and DIFF as opposing poles of a coordinated epithelial axis rather than independently varying marker-defined populations. The axis occupies a defined position within the broader regenerative, oncofetal and revival-like state space adopted by colorectal cancer cells under oncogenic and therapeutic stress, without being equivalent to any one of these configurations (16–19,23,41). Importantly, folate availability did not remodel all cells in parallel along this axis. Under folate deprivation, differentiated cells acquired progenitor-associated features, whereas stem-like cells redistributed into adaptive intermediate states. Within-state comparisons reinforced this asymmetry. SCL cells mounted the largest transcriptional response to folate restoration, while DIFF cells underwent the most extensive accessibility remodeling. Folate therefore acts on a pre-existing epithelial landscape, with the pre-existing epithelial state strongly shaping the regulatory response to the same metabolic perturbation.

These observations resolve an apparent temporal tension in the response. Changes in SOX9 and KRT20 transcription were detectable within four hours of folate restoration, whereas redistribution of the reporter-defined populations first became detectable at 24 hours and was pronounced by 48 hours. This lag may in part reflect the additional time required for synthesis, folding and maturation of the encoded fluorescent proteins, while also being compatible with a delayed population-level manifestation of the transcriptional response. Chromatin remodeling along the trajectory was gradual rather than switch-like, whereas the dominant polarization of the SCL–DIFF program pair emerged later, with its largest step occurring between the eighth and ninth pseudotime bins. Taken together, the data place early transcriptional and regulatory remodeling temporally upstream of the overt population-level polarization of the axis.

The trajectory-resolved chromatin response was strongly asymmetric, with 1,023 regions losing accessibility and only 201 gaining it. This predominance of closing regions suggested that repressive chromatin mechanisms contribute substantially to the folate-induced transition. Metabolic rescue experiments further linked this response to cellular methylation potential. Structurally distinct folate intermediates produced comparable chemosensitization and comparable NKD1 and TGM2 responses, while SAM itself independently reproduced the sensitizing effect, indicating that methyl-donor availability can contribute to the folate-responsive phenotype. Absolute SAM and SAH pools fell transiently while the SAM/SAH ratio increased, a divergence consistent with increased cellular methylation potential and rapid turnover within the methylation cycle. H3K27me3 showed the most pronounced and temporally structured response among the histone marks examined. After EZH2 inhibitor washout, H3K27me3 recovered within 24 hours only when folate was present, showing that folate availability supports efficient H3K27me3 re-deposition following release from EZH2 inhibition. Together, these findings link folate availability, cellular methylation potential and H3K27me3 remodeling without implying that the sensitizing effects of individual folate metabolites are mediated exclusively through SAM. This mark- and context-specific response is consistent with evidence that one-carbon perturbation does not produce uniform hypermethylation but affects particular modifications and loci according to metabolite availability, enzyme kinetics and chromatin context (3–5,42).

The relationship between EZH2 activity and colorectal epithelial differentiation is therefore unlikely to be monotonic. In KRAS-mutant CRC models, Loi et al. showed that combined EZH2 and RAS-pathway inhibition drives differentiation by derepressing PRC2 targets, including negative regulators of WNT signaling (43). This differs from the present setting, in which folate restoration transiently increased bulk H3K27me3 levels while producing selective chromatin closure at regions overlapping H3K27me3-enriched domains in primary colorectal tumors (44,45), together with attenuation of canonical WNT activity in HT29 cells. Together, these observations distinguish methyl-donor supply from the genomic targeting of PRC2-dependent repression. Folate availability regulates the capacity for H3K27me3 deposition, but the resulting epithelial-state output depends on where PRC2-dependent repression is established or relieved. Differentiation therefore cannot be inferred from bulk H3K27me3 abundance alone, but instead reflects a locus- and context-dependent regulatory response (22).

Downstream of this chromatin layer, folate uncoupled transcription-factor abundance from regulatory engagement. AP-1 motif accessibility declined even as c-FOS and JUN expression was maintained or increased (35–37). Within canonical WNT signaling, the same uncoupling involved a lineage-defining regulatory complex, as TCF7L2 and β-catenin occupancy decreased at regions that simultaneously gained accessibility and H3K27ac following folate restoration. Consistent with reduced canonical WNT engagement, β-catenin redistributed from the nucleus toward a membrane-associated pool and β-catenin–TCF7L2 proximity declined. Chromatin opening is therefore not equivalent to transcription-factor activation. Instead, folate-induced differentiation reflects redistribution of regulatory occupancy across a remodeled landscape in which output is determined jointly by element accessibility, factor abundance, subcellular localization and cooperative binding (39). NKD1 and TGM2 emerged as recurrent folate-responsive loci within this regulatory landscape, both overlapping super-enhancer annotations in the integrated regulatory ranking (46) and responding across the graded folate series. NKD1 additionally responded to methotrexate and 5-FU, making it a useful mechanistic readout at the intersection of folate-responsive epithelial remodeling and antimetabolite exposure.

Prospectively isolated SCL, DIFF, double-positive and double-negative fractions from the same HT29 model differed several-fold in 5-FU IC_50_ under matched folate conditions. Folate restoration further altered sensitivity within individual states rather than shifting the population uniformly. The double-negative fraction moved against the general trend, showing that the system is not a linear stemness-to-resistance scale. Two additional observations argue against a proliferation-only account. Stem-like cells were least fluoropyrimidine-sensitive precisely when their S-phase occupancy was highest, and folate restoration reduced S-phase occupancy in the sorted states while increasing population-level cell density and 5-FU sensitivity. Snapshot S-phase fraction is not equivalent to proliferative output, and high S-phase occupancy alone does not predict fluoropyrimidine sensitivity within this epithelial-state system.

Post-treatment endpoints were likewise non-parallel. Colony frequency and colony size did not change together, distinguishing the frequency of clonogenic survivors from the subsequent outgrowth of their progeny. Folate also reduced, whereas methotrexate increased, the accumulation of TIS-associated cells. These findings are consistent with pretreatment epithelial state predisposing cells to distinct post-treatment phenotypes and with the broader recognition that drug-tolerant colorectal cancer cells can occupy defined, reversible states rather than a fixed resistant identity (24,25). In breast cancer models, we previously showed that therapy-induced senescence can be transient rather than terminal (47), further supporting the view of TIS as a potentially reversible drug-tolerant state.

Together, these observations distinguish three dimensions of treatment response, the epithelial state present before treatment, the frequency of cells that retain clonogenic potential, and the proliferative or senescence-associated phenotype adopted after exposure. Their non-parallel behavior illustrates why neither S-phase occupancy nor a single viability endpoint is sufficient to capture treatment response. Together with the acute mitochondrial adaptation described by Moss et al. (26), these findings place fluoropyrimidine response across distinct temporal layers, from the epithelial state established before treatment, through the metabolic response to 5-FU, to later persistence phenotypes reflected by clonogenic recovery and TIS-associated accumulation.

The same axis is recognizable in primary human tissue. Across a published single-nucleus atlas of normal mucosa, adenoma and carcinoma (40), SCL activity was higher and DIFF activity lower in carcinoma than in normal mucosa, with adenomas intermediate and heterogeneous, and these two programs defined the extremes of the entire collection. Folate restoration moves the same programs in the opposite direction, and trajectory-responsive regions overlapped H3K27ac- and H3K27me3-defined domains in colorectal tumors (44,45), while accessibility at trajectory-opening regions was associated with overall survival in TCGA (33). This directional correspondence establishes tissue relevance but does not imply that folate exposure itself controls colorectal tumor progression in vivo, a caution reinforced by epidemiological and experimental studies showing that the association between folate and colorectal tumorigenesis depends on genotype, dose and timing rather than exposure alone (48–51).

Clinically, the implication is not that more or less folate should be given, but that its therapeutic consequence may depend on the epithelial and metabolic state present when fluoropyrimidine treatment begins. Folinic acid co-administration is currently applied without reference to either, while our data show that the effect of folate restoration on clonogenic survivor frequency changed with preconditioning duration even as its effect on colony-size composition persisted. Because acute target potentiation and prospective state remodeling operate on different timescales, their superposition may contribute to the long-recognized schedule dependence of methotrexate–fluoropyrimidine combinations (52–54), with the balance between them shaped by the interval between folate or antifolate exposure and treatment. The chemical form may matter as well, since folic acid and reduced folates enter the cycle at different points and human dihydrofolate reductase reduces folic acid slowly and variably (55). Prospective studies should therefore test whether the SCL–DIFF configuration present at treatment onset is associated with fluoropyrimidine response and clinical outcome, and whether it modifies the benefit of folinic acid co-administration.

### Study Limitations

The mechanistic dissection is anchored in the genetically and epigenetically well-characterized HT29 model, in which the reporter system permits prospective isolation of defined epithelial states (56,57). The therapeutic direction reproduced across seven additional colorectal cancer cell-line models, spheroids and four patient-derived organoids, and the axis itself was recognizable in primary human tissue. Extending the full chromatin mechanism across colorectal cancer molecular contexts, including distinct consensus molecular subtypes (58), MSI tumors and BRAF wild-type backgrounds, is the natural next step. Folate remodels epithelial state and alters drug response, and the sorted-fraction data establish that state stratifies sensitivity. Direct perturbation-based tests of mediation would determine how much of the pharmacological effect passes through state remodeling and how much arises from canonical thymidylate synthase potentiation acting in parallel. Thymidine rescue would further distinguish methyl-donor from nucleotide contributions. TIS burden is quantified as occupied culture area, providing a population-level rather than cell-number or lineage-resolved readout. The human atlas is cross-sectional and carries no folate-exposure information, and the survival association in TCGA is exploratory. Both point toward prospective cohorts with defined metabolic and epithelial states as the appropriate test.

### Conclusion

Together, these findings support a model in which one-carbon metabolism acts as an upstream metabolic input into epithelial plasticity, modulating cellular occupancy along a polarized SCL–DIFF axis. Folate availability is accompanied by changes in methylation potential, chromatin organization and transcription-factor engagement, thereby shaping the pretreatment epithelial configuration in which fluoropyrimidine response unfolds. This framework extends the canonical thymidylate synthase model by separating acute biochemical potentiation from slower, metabolically regulated state remodeling. More broadly, our findings position epithelial state as an interface between metabolic availability and therapeutic response, showing that metabolism not only influences the biochemical target of an antimetabolite but also helps define the cellular state on which treatment acts.

## Methods

### Cell culturing and folate manipulation

Colorectal cancer cell lines (HT29, HT29 DER, HT115, DLD1, SW48, LS180, KM12-SM and Colo 320) were maintained under standard culture conditions at 37 °C in 5% CO₂ in RPMI-1640 medium supplemented with 10% FBS, 1% penicillin/streptomycin, 1% sodium pyruvate, 1% non-essential amino acids (NEAA), and 1% HEPES (all from Gibco, Thermo Fisher Scientific). For folate-controlled experiments, folic acid–free RPMI-1640 (Gibco, Cat# 27016021) supplemented with 10% dialyzed FBS (Gibco, Thermo Fisher Scientific) was used as the base medium. Cells were subjected to folate starvation (FA⁰; folic acid–free medium) or folate treatment by supplementation with folic acid (Sigma-Aldrich, F8758) at defined concentrations (FAᴸ, 2.25 μM, corresponding to the folate concentration of standard RPMI-1640; FAᴴ, 22.5 μM, a tenfold-elevated supraphysiological dose) and time points. Conditioning intervals were fixed in advance on the basis of preliminary marker time courses: 72 h for folate deprivation and 24–48 h for folate supplementation. Where indicated, cells were preconditioned for 5 days prior to downstream analyses. Cell lines were routinely tested for mycoplasma once a month to ensure they were free of contamination using MycoAlert Mycoplasma Detection Kit (Lonza, Biocenter Ltd, Hungary) according to the manufacturer’s protocol.

### Dual endogenous reporter (DER) system

A dual endogenous reporter (DER) system in HT29 colorectal cancer cells (HT29 DER cells) was used to monitor cellular state transitions (30). In this system, mKate2 reports stem-like (SCL) states associated with *SOX9* expression, while GFP labels differentiated (DIFF) epithelial states associated with *KRT20* expression. Reporter activity was assessed by flow cytometry.

### Flow cytometry analysis

Reporter distributions were analyzed on an Attune NxT flow cytometer (Thermo Fisher Scientific, Waltham, MA, USA). Debris was excluded on forward and side scatter, and single cells were selected in three sequential gates, FSC-H versus FSC-W, then SSC-H versus SSC-W, then FSC-A versus SSC-A. No viability dye was used. Fluorescence thresholds were zeroed against parental HT29 cells lacking the reporter cassettes. Live single cells were then divided into four quadrants by mKate2 and GFP signal, defining the stem-like (SCL, mKate2⁺GFP⁻), differentiated (DIFF, GFP⁺mKate2⁻), double-positive (DP) and double-negative (DN) populations, and the frequency of each population was read from these quadrants.

### Fluorescence-activated cell sorting

Cells were sorted on a BD FACSAria III cell sorter (BD Biosciences) using the same debris, singlet and threshold settings. Four-way sorting was performed with mutually exclusive gates, and the stem-like (SCL) and differentiated (DIFF) fractions were isolated as the 2% of live single cells with the highest mKate2 and the highest GFP signal respectively, alongside the double-positive (DP) and double-negative (DN) fractions and the unsorted population (POP). Sorting used a 70 μm nozzle at 70 psi with a drop-formation frequency of approximately 89 kHz and standard sheath fluid, and sorted cells were collected directly into culture medium. Post-sort purity exceeded 96%.

### RNA extraction and qRT–PCR analysis

Total RNA was extracted using the RNeasy Mini Kit (Qiagen, Hilden, Germany) according to the manufacturer’s instructions. RNA quantity and quality were assessed using a NanoDrop One/OneC Microvolume UV-Vis Spectrophotometer (Thermo Fisher Scientific, Rockland, DE, USA); samples with a 260/280 ratio above 1.8 were used for further analysis. RNA was reverse transcribed using the MultiScribe Reverse Transcription Kit (Thermo Fisher Scientific, Waltham, MA, USA), and qPCR was performed on a QuantStudio 3 Real-Time PCR System using Power SYBR Green master mix (Thermo Fisher Scientific) with the following thermal profile: one cycle of 95 °C for 2 min, then 40 cycles of 95 °C for 5 s, 60 °C for 20 s, and 72 °C for 15 s, with fluorescence acquisition at each cycle. Gene-specific primers were designed using the Primer3Plus tool; primer sequences are summarized in **Table S7A**. Raw data were analyzed using the relative quantification (Livak) method with β-actin (ACTB) as the internal control gene.

### Immunofluorescence staining

HT29 cells were seeded in FNC-coated eight-well µ-Slides (Ibidi). At the indicated experimental endpoints, cells were washed with 1× PBS, fixed with 4% paraformaldehyde for 15 min, permeabilized with 0.1% Triton X-100 in PBS for 10 min and blocked with 5% bovine serum albumin in PBS for 15 min at room temperature. Cells were incubated with primary antibodies against SOX9, KRT20, β-catenin, H3K27me3 or p21 overnight at 4 °C, followed by the appropriate fluorescently labelled secondary antibodies for 2 h at room temperature. Nuclei were counterstained with DAPI. Images were acquired using a Nikon Eclipse Ti inverted fluorescence microscope (Nikon Instruments, Tokyo, Japan), and fluorescent slides were additionally digitized on a PANNORAMIC MIDI II whole-slide scanner (3DHISTECH Ltd., Budapest, Hungary). Antibodies and working dilutions are listed in **Table S7B.**

### Proximity ligation assay (PLA)

Protein–protein interactions and protein localization were assessed using the Duolink In Situ Red Starter Kit Mouse/Rabbit (Sigma-Aldrich, DUO92101) according to the manufacturer’s protocol. Two antibody pairings were used: β-catenin/TCF7L2 to detect the β-catenin–TCF7L2 transcriptional complex, and β-catenin/β-catenin to quantify β-catenin abundance and determine its subcellular localization. Primary antibodies are listed in **Table S7B**. Cell nuclei were counterstained with DAPI. Proximity ligation signals were quantified to evaluate interaction dynamics and β-catenin distribution across folate conditions.

### Metabolite treatments

Following 72 h of folate starvation (FA⁰), cells were treated with the following folate-pathway metabolites at defined concentrations and time points to assess transcriptional and functional effects: folic acid (FA), dihydrofolic acid (DHF), tetrahydrofolic acid (THF), folinic acid (leucovorin), 5-methyltetrahydrofolic acid (5-MTHF), and S-adenosylmethionine (SAM). FA, DHF, and THF were purchased from Sigma-Aldrich (Merck Life Science Kft, Budapest, Hungary); the remaining compounds were purchased from MedChemExpress (Monmouth Junction, NJ, USA).

### Mass spectrometry analysis

Folate and related one-carbon and methylation-cycle metabolites, including folate (FA), dihydrofolate (DHF), tetrahydrofolate (THF), 5-methyltetrahydrofolate (5-MTHF), S-adenosylmethionine (SAM), S-adenosylhomocysteine (SAH) and leucovorin, were quantified by tandem mass spectrometry using a Sciex 6500 QTRAP instrument equipped with a Turbo V™ ion source, operating in positive electrospray ionization and multiple reaction monitoring (MRM) mode. Source parameters were as follows: ion spray voltage, 5500 V; nebulizer, drying, and curtain gases, 45, 45, and 40 psi, respectively; drying gas temperature, 500 °C. The dwell time for each MRM transition was 80 ms, with a 5-ms pause between transitions, yielding an overall cycle time of 1.36 s. MRM transitions for all analytes are listed in **Table S5A**.

Chromatographic separation was performed on an Agilent Zorbax RX-SIL normal-phase column under hydrophilic interaction liquid chromatography (HILIC) conditions, based on previously established methods for the separation of nucleobases and other polar basic compounds (59,60). Acetonitrile and water, both containing 0.1% formic acid, served as mobile phases, with the aqueous phase acting as the stronger eluent. The flow rate was 1.0 mL min⁻¹ over a 10-min gradient program: the initial aqueous content of 20% was increased to 80% over 4.5 min and held for 1 min, after which the initial composition was re-established within 0.5 min and maintained for 4 min of column equilibration.

Absolute calibration was performed for folate over a concentration range of 0.1–1000 ng/mL using a five-point calibration curve, with lower-range points injected in replicate to assess reproducibility. Linear regression with 1/x weighting yielded excellent linearity (r = 0.99979), with accuracy meeting acceptance criteria of ±15% (±20% at the lowest concentration level). This approach confirmed the absence of detectable folate in the completed folic acid–free (FA⁰) medium and verified the folate content of FAᴸ (2.25 μM) medium (calibration curve and medium validation shown in **Fig. S5A–C**).

For intracellular metabolite quantification, including SAM and SAH, culture supernatant was diluted 1:3 with acetonitrile and centrifuged to remove cell debris. Cell pellets were washed with 1× PBS, and high-purity 80% methanol was added prior to centrifugation. SAM/SAH ratios were calculated to estimate cellular methylation capacity.

### In vitro cytotoxicity assay

A total of 5,000 (2D cultures) or 12,000 (3D cultures) cells per well were plated in 96-well plates and treated with 5-fluorouracil (5-FU) or methotrexate (MTX) across a serial dilution range (starting at 100 μM, two-fold dilutions) for 5 days. Viability was assessed using the PrestoBlue assay (Thermo Fisher Scientific, Waltham, MA, USA) according to the manufacturer’s instructions: cells were incubated for 1 h with 5% PrestoBlue Cell Viability Reagent diluted in 1× PBS at 37 °C and 5% CO₂. Fluorescence was measured using an EnSpire microplate reader (PerkinElmer, Waltham, MA, USA). Data were normalized to untreated control cells, and IC_50_ values were determined by nonlinear regression using a four-parameter logistic model in GraphPad Prism v8.0.1 (GraphPad Software, Boston, MA, USA).

### Tazemetostat treatment and H3K27me3 recovery after EZH2 inhibition

Tazemetostat concentration and treatment duration were established by titration, and 1 μM for 120 h was selected as the shortest exposure at the lowest concentration that produced near-complete H3K27me3 depletion. HT29 cells were seeded at 75,000 cells per well in 12-well plates in folic acid-free medium (FA⁰) and treated with 1 μM tazemetostat for 120 h. The medium and inhibitor were not replaced during the 120-h treatment period. Tazemetostat was then removed, cells were washed with pre-warmed PBS and recovery was initiated by adding either folate-free medium (FA⁰) or folate-containing medium (FAᴸ; 2.25 μM folic acid). Samples were collected immediately after inhibitor washout (0 h) in both arms, and subsequently after 24 h of recovery in FA⁰ medium or after 2, 4, 6 and 24 h of recovery in FAᴸ medium. Cells maintained in FAᴸ medium for 24 h without tazemetostat exposure served as an untreated reference control. H3K27me3 recovery was assessed by immunoblotting, and band intensities were quantified by densitometry using ImageJ and normalized to ACTB. The experiment was performed in three independent biological replicates.

### Colony formation assay

HT29 cells were plated at low density (100,000 cells per 90-mm dish) and cultured under defined conditions for at least 3 weeks. After the growth period, colonies were fixed with 100% methanol for 20 min at room temperature and stained with 0.5% crystal violet for 5 min at room temperature. Stained colonies were imaged and quantified using ImageJ v1.54s. See more details in **Fig. S1H–J** for quantification and statistical analysis.

### Therapy-induced senescence (TIS) induction and conditions

For TIS induction, 1 × 10^6^ HT29 cells were plated on 90-mm culture dishes 24 h prior to treatment with 20 µM 5-FU for 5 days. Cells were then washed with pre-warmed 1× PBS, the medium was renewed, and dishes were maintained for a further 7 days in drug-free medium. This 12-day endpoint was chosen because dying and transiently surviving cells are largely cleared during the post-treatment interval, whereas repopulation from treatment-surviving clones begins only after approximately 21 days, so the 7–14 day window after drug withdrawal is enriched for cells with a stable senescence-associated phenotype. Senescence was validated by p21 immunofluorescence and SA-β-Gal staining performed in parallel on the same conditions, and representative images are shown for both. All condition-level quantification was performed on brightfield images as occupied culture area.

### SA-β-Gal staining

Cells were plated into 96-well tissue culture plates. Senescence-associated β-galactosidase (SA-β-Gal) staining was performed using the Senescence β-Galactosidase Staining Kit (Cell Signaling Technology, Danvers, MA, USA) according to the manufacturer’s instructions.

### Folate/antifolate conditioning and cell-coverage quantification

For the folate arm, HT29 cells (1 × 10^6^ per dish) were seeded in folate-free (FA⁰) medium and maintained under folate deprivation for 72 h. Cells were then either kept in FA⁰ (reference) or repleted with low-folate (FAᴸ) medium and exposed to 20 μM 5-fluorouracil (5-FU) either immediately (FAᴸ 0 h) or after a 24 h folate pre-treatment (FAᴸ 24 h); the 0 h and 24 h designations refer to the duration of folate repletion preceding 5-FU addition, not to the length of starvation. For the antifolate arm, cells were seeded in normal (folate-replete) medium and pre-treated with methotrexate (MTX; 0.03 or 0.1 μM) for 24 h before the addition of 20 μM 5-FU. After 5 days of 5-FU treatment and a further 7 days in drug-free medium, cell coverage (occupied area, %) was quantified from brightfield images sampled across twelve predefined fields per dish using ImageJ (see **Fig. S1K,L** for the imaging strategy). FA⁰ served as the reference condition for the folate arm and normal folate-replete medium for the antifolate arm, and statistical significance was assessed by Welch’s two-sided t-test on biological replicate means (n = 6).

### 3D spheroid and patient-derived organoid (PDO) experiments

Experiments with human samples were approved by the Medical Research Council of Hungary (Wiener Z, ETT-TUKEB No. 51323-4/2015/EKU), and informed consent was obtained from all patients. Three-dimensional HT29 spheroids and four patient-derived colorectal cancer organoids, kindly provided by Z. Wiener (Semmelweis University), were cultured in RPMI-1640 (or folic acid– free RPMI-1640, Gibco Cat# 27016021, for folate-controlled experiments) supplemented with B27, nicotinamide, N-acetylcysteine, 10 µM SB202190, 500 nM A83-01, 50 ng/mL EGF, and 1% each of penicillin/streptomycin, HEPES, and glutamine (Thermo Fisher Scientific, Gibco). Organoids were mechanically removed from the 3D matrix every 5–7 days, centrifuged at 600 × g for 5 min, washed with 1× PBS, and digested with TrypLE (Thermo Fisher Scientific) until dissociated into clusters of 5– 10 cells, then washed and re-embedded.

Importantly, the folate content of all matrices was verified by mass spectrometry: Matrigel (Corning) and Geltrex (Thermo Fisher Scientific, A1048301) were found to contain detectable folate and were therefore used only for organoid maintenance, whereas VitroGel ORGANOID (TheWell Bioscience, WB-VHM04-3; xeno-free) and rat-tail collagen I (60% distilled water, 10% 10× MEM [Gibco], 30% collagen type I [Ibidi, Gräfelfing, Germany]) were confirmed to be folate-free and were used for all folate-controlled experiments. Folic acid treatments under defined folate conditions and 5-FU drug responses (PrestoBlue assay) were assessed in these systems. All components were tested for folic acid content by mass spectrometry.

### Bulk RNA-seq of HT29 cells under folate treatment

HT29 cells were cultured under three folate conditions, FA⁰ (folic acid–free control), FAᴸ (2.25 μM low-dose) and FAᴴ (22.5 μM high-dose), and harvested at two time points (2 days and 14 days), with two biological replicates per condition, yielding 12 paired-end RNA-seq libraries in total. Raw reads were quality-controlled using FastQC (v0.11.9) (61) and adapter-trimmed with Trimmomatic (v0.39) (62) in paired-end mode (LEADING:3; TRAILING:3; SLIDINGWINDOW:4:20; MINLEN:50); only properly paired surviving reads were retained, and post-trimming quality was re-assessed with FastQC.

Trimmed reads were aligned to the human reference genome (GRCh38) using STAR (v2.7.10a) (63) in two-pass mode (--twopassMode Basic; --outSAMunmapped Within), with alignments written as coordinate-sorted BAM files. BAM files were indexed using SAMtools (v1.16.1) (64), and normalized coverage tracks were generated with BAMscale (v1.0) (65) in RNA-seq scaling mode. Gene-level counts were obtained with featureCounts (Rsubread v2.18.0) (66) against the NCBI RefSeq GRCh38.p14 annotation, summarizing uniquely mapped, properly paired fragments overlapping a single exonic feature (allowMultiOverlap = FALSE).

Differential expression analysis was performed using DESeq2 (v1.38.3) (67). To disentangle dose effects from treatment duration, samples were partitioned by duration and analyzed independently within the 2-day and 14-day subsets, with folate dose (FA⁰, FAᴸ, FAᴴ) modeled as a single factor (∼dose) and FA⁰ set as the reference level. Genes with at least 10 reads in two or more samples within the relevant subset were retained for model fitting. Size factors were estimated by the median-of-ratios method, dispersions by the default mean–dispersion relationship, and a negative-binomial generalized linear model was fitted using the standard DESeq2 workflow. Principal component analysis on variance-stabilizing-transformed (vst) counts (blind = FALSE) was used for quality assessment and to verify expected clustering by dose. Within each duration subset, three pairwise contrasts were extracted (FAᴸ vs FA⁰, FAᴴ vs FA⁰, and FAᴴ vs FAᴸ), and log2-fold-changes were shrunk using the adaptive shrinkage estimator (lfcShrink, type = “ashr”) (68). p values were adjusted by the Benjamini–Hochberg procedure, and genes were considered differentially expressed at an adjusted *p* value < 0.05 and |log_2_ fold change| > 0.5. The six resulting differential expression tables (three contrasts × two durations; **Fig. S2A–C**) were used to characterize dose-dependent transcriptional responses to folate availability and were integrated with matched chromatin profiling data.

### Single-cell multiome library preparation and sequencing

The HT29 dual endogenous reporter (DER) cell line, in which GFP marks differentiated (*KRT20*-associated) cells and mKate2 marks stem-like (*SOX9*-associated) cells (30), was used for two separate multiome experiments. For the endpoint experiment, cells were cultured under folate starvation (FA⁰) or folate treatment (FAᴸ, 2.25 μM, 48 h), and the unsorted population (POP) together with the top-2% mKate2⁺ (SCL) and top-2% GFP⁺ (DIFF) fractions were isolated from each condition, yielding six populations. For the trajectory experiment, cells were treated with 2.25 μM folic acid and harvested at ten time points (0, 1, 2, 3, 4, 6, 12, 24, 36 and 48 h) following folate addition; equal numbers of cells from all ten time points were then combined into a single mixed-time-point sample to enable pseudotemporal reconstruction within one multiome reaction.

Following purification, all resulting samples were processed in parallel for nuclei isolation and used with the Chromium Next GEM Single Cell Multiome ATAC + Gene Expression workflow (10x Genomics) according to the manufacturer’s instructions. After nuclei quantification and QC, 7,500 nuclei were loaded per reaction for an expected recovery of ∼5,000 nuclei. Chromatin accessibility was captured by transposition prior to microfluidic partitioning; transposed nuclei were then encapsulated with barcoded gel beads in the Chromium controller, enabling simultaneous indexing of accessible chromatin fragments and nuclear mRNA. ATAC and gene expression libraries were constructed according to the standard Multiome protocol, evaluated for concentration and fragment-size distribution, and sequenced on an Illumina platform at the recommended depth for paired ATAC and gene expression profiling.

### Single-cell multiome data processing and integration

Libraries were processed using Cell Ranger ARC (v2.0.2, 10x Genomics) (69) with the GRCh38 reference (refdata-cellranger-arc-GRCh38-2020-A). The cellranger-arc count pipeline was run with default parameters using a sample sheet specifying matched RNA and ATAC libraries, performing demultiplexing, alignment, barcode processing, and generation of gene expression and chromatin accessibility count matrices.

Downstream analysis was performed in R (v4.4.0) using Seurat (v5.1.0) (70), Signac (v1.14.0) (71), and ArchR (v1.0.2) (72), across experimental conditions including GFP-reporter (DIFF), mKate2-reporter (SCL), population (POP) controls, and folate addition time-course samples. Gene annotations for hg38 were obtained from EnsDb.Hsapiens.v86 (v2.99.0). Chromatin accessibility assays were constructed from matched fragment files, with peaks identified using MACS2 through the Signac CallPeaks interface, restricted to standard chromosomes and excluding ENCODE blacklist regions; a consensus peak-by-cell matrix was generated using Signac.

Nuclei were retained with 1,000 < ATAC fragments < 100,000, 1,000 < RNA UMIs < 25,000, more than 1,500 detected genes, nucleosome signal < 2 and TSS enrichment > 1; no mitochondrial-content filter was applied. Nucleosome signal and TSS enrichment were computed with Signac (NucleosomeSignal, TSSEnrichment) prior to filtering, and identical thresholds were used for the endpoint and the trajectory datasets. Doublets were identified using scDblFinder (v1.18.0) (73) within a SingleCellExperiment (v1.26.0) framework and removed prior to downstream analysis. For RNA, data were log-normalized, 3,000 highly variable features identified, cell-cycle scores computed with Seurat CellCycleScoring using the canonical S and G2/M gene sets (cc.genes.updated.2019) and regressed during scaling, followed by PCA, k-nearest-neighbor graph construction, graph-based clustering, and UMAP visualization. ATAC data were normalized by TF-IDF transformation and reduced by latent semantic indexing (LSI); informative peaks were selected with FindTopFeatures and visualized by UMAP. For multi-condition datasets, batch effects were corrected using Harmony (v1.2.3) (74) with group.by.vars = “cond” (the six sample identities) on PCA (RNA) and LSI (ATAC) embeddings.

Multimodal integration used Seurat’s weighted nearest-neighbor (WNN) framework (FindMultiModalNeighbors), which constructs modality-specific neighbor graphs and computes cell-specific modality weights based on cross-modality prediction accuracy. For the six-sample endpoint dataset, Harmony-corrected RNA (PCA, dimensions 1–20) and ATAC (LSI, dimensions 2–20) embeddings were used as the modality inputs. The folate addition time-course sample (0, 1, 2, 3, 4, 6, 12, 24, 36 and 48 h following folate treatment) was generated as a single multiome reaction, and no between-sample batch correction was applied. For the time-course dataset, modality-specific UMAP embeddings were used as inputs to FindMultiModalNeighbors. These two-dimensional embeddings were generated from RNA PCA dimensions 1–15 and ATAC LSI dimensions 2–10 and resolved the transitional epithelial states along the folate response more clearly than the linear reductions. This differs from the standard WNN implementation, in which the PCA and LSI reductions are used directly. Because UMAP is a nonlinear, neighborhood-preserving embedding, modality weights are computed from the local neighborhood structure represented in UMAP space rather than directly from the structure captured by the linear reductions. We therefore evaluated whether the resulting trajectory was consistent with independent experimental measurements. The differentiated epithelial state was located at late pseudotime, where the independently derived DER_DIFF transcriptional and accessibility signatures were enriched (**Fig. 3C,E,G**). The trajectory direction was also consistent with the population-level response measured by flow cytometry, which showed an increase of 12.5 percentage points in the differentiated fraction and a decrease of 10.8 percentage points in the stem-like fraction following folate supplementation (**Fig. 1B**). In addition, the pseudotemporal changes in SOX9, KRT20 and NKD1 were consistent with the temporal patterns measured by qRT–PCR (**Fig. S1D**; **Fig. 3O**) and immunoblotting (**Fig. 1D,E**; **Fig. 4I**).

### Pseudotime and trajectory analysis

A three-dimensional UMAP embedding was computed from the WNN graph using Seurat (RunUMAP, n.components = 3). Trajectory inference was performed using Monocle3 (v1.3.7) (75): the Seurat object was converted into a cell_data_set using SeuratWrappers, retaining the WNN-derived UMAP coordinates to preserve multimodal structure, and existing Seurat cluster annotations were carried over. A principal graph was learned using learn_graph without partition constraints. Cells were ordered in pseudotime using order_cells, with root cells selected from a cluster enriched for the POP folate-response differential expression signature. Cells were assigned to ten equally populated bins on descending Monocle3 pseudotime, so that bin PT1 corresponds to the state most distant from the root and PT10 to the root population; this orients the bins along the folate response from SCL to DIFF. Ten bins were used throughout unless otherwise indicated, and a finer partition into twenty equally populated bins was applied where higher temporal resolution was required, including the regulatory module analyses shown in **Fig. 4B, D, F**. Trajectory-associated genes were identified using Monocle3 graph_test. Both continuous and discretized pseudotime representations were used to quantify transcriptional and chromatin accessibility changes across the response.

### Differential accessibility, gene expression, and functional enrichment

Differential analyses across trajectory-defined cell populations were performed using Seurat (v5.1.0). Late (clusters 8–10) and early (clusters 1–3) trajectory states were compared using FindMarkers (Wilcoxon rank-sum test). Differential chromatin accessibility was assessed using peak counts, and transcription factor activity using chromVAR deviation scores (average differences computed as row-wise means, reported as avg_diff); differential gene expression was computed on the RNA assay with default parameters. To complement single-cell testing, pseudobulk profiles were generated using AggregateExpression, grouping cells by discretized trajectory bins. Raw count matrices were aggregated for the RNA assay, whereas normalized values were aggregated for the ATAC peak assay and for chromVAR deviation scores, preserving the normalization applied to each.

Regulatory interpretation of differentially accessible regions was performed using the rGREAT Bioconductor package (v4.0.4) (76) with hg38 gene annotations (TxDb.Hsapiens.UCSC.hg38.knownGene) and MSigDB Hallmark gene sets. Up- and down-regulated peak sets were analyzed separately using the consensus accessible peak set as the background region set, and significant enrichments were defined by an adjusted hypergeometric P value (p_adjust_hyper) < 0.05. Gene-level functional enrichment of differentially expressed genes used clusterProfiler (4.2.2) (77) with GO (GO.db), KEGG (org.Hs.eg.db), and MSigDB gene sets (msigdbr); over-representation analysis was additionally performed using Enrichr (Reactome, WikiPathways, HDSigDB). Gene set enrichment analysis (GSEA) was performed on ranked gene lists derived from log2 fold-change statistics, using normalized enrichment scoring and FDR correction.

### Chromatin immunoprecipitation sequencing (ChIP-seq)

Folate-treated (FAᴸ) and folate-starved (FA⁰) HT29 cells were cross-linked either with 1% formaldehyde alone (histone ChIP) or with 2 mM disuccinimidyl glutarate (DSG; Thermo Fisher Scientific) followed by 1% formaldehyde (transcription factor ChIP), and quenched with 0.125 M glycine. Cells were washed twice in cold PBS, resuspended in lysis buffer (10 mM Tris-HCl pH 8.0, 1 mM EDTA, 1% SDS, protease inhibitor cocktail), and sonicated for 35 min using a PIXUL sonicator (Active Motif; Pulse 50, PRF 1 kHz, burst rate 20 Hz). SDS was quenched with 1% Triton X-100 and 150 mM NaCl, and 10% of the sheared chromatin was retained as input to estimate concentration and shearing efficiency.

DNA–protein complexes were immunoprecipitated according to the manufacturer’s instructions using antibodies against H3K27ac (Active Motif, Cat#91193), TCF4/TCF7L2 (C48H11; Cell Signaling Technology, Cat#2569), and β-catenin (D10A8 XP; Cell Signaling Technology, Cat#8480). Sheared chromatin (20–30 μg per reaction) was incubated overnight at 4 °C, and complexes were pulled down with Dynabeads Protein G (Thermo Fisher Scientific) and washed sequentially with low-salt, high-salt, LiCl, and TE buffers. ChIP-enriched chromatin was de-crosslinked overnight at 65 °C, and DNA was extracted using a DNA Clean & Concentrator kit (Zymo Research). ChIP-seq libraries were prepared using the NEBNext Ultra II DNA Library Prep Kit for Illumina (New England Biolabs) and paired-end sequenced.

### ATAC-seq

For ATAC-seq, 75,000 freshly collected folate-treated (FAᴸ) and folate-starved (FA⁰) HT29 cells were processed in duplicate using the Active Motif ATAC-seq Kit (Cat#53150). Tagmented DNA fragments were indexed using the Nextera adapters provided in the kit. Following PCR amplification, ATAC libraries were purified and paired-end sequenced.

### Transcription factor activity and gene regulatory network (GRN) analysis

Chromatin accessibility and transcription factor activity were quantified using chromVAR (v1.26.0) (78), motifmatchr (v1.26.0), TFBSTools (v1.42.0) (79), and JASPAR2020 (v0.99.10) (80) position weight matrices, with GC bias–corrected background models and regulatory elements annotated relative to transcription start sites using hg38 coordinates. Peak–gene associations were inferred using Signac (v1.14.0) based on co-accessibility and genomic proximity.

Gene regulatory networks were inferred using Pando (v1.1.1) (81). Candidate regulatory regions were defined as the phastCons 20-mammal conserved elements distributed with Pando, restricted to those overlapping the multiome peak set. Candidate regions were scanned for TF motifs using the Pando motif and motif-to-TF databases for hg38, peaks were assigned to genes using the Signac-based linking procedure, and TF–target relationships were modelled by generalized linear regression with a Gaussian family, scaled predictors and a TF–TF correlation cutoff of 0.05. Models were fitted across the cells of the time-course multiome library. Interactions with adjusted *p* < 0.05 were retained, and regulatory modules were extracted using a coefficient *p* < 0.05, at least two variables per module and R² ≥ 0.05, to derive TF- and regulon-associated activity scores.

scMEGA (v1.1.0) (82) was applied independently as a complementary analysis. A diffusion map was computed from the WNN UMAP embedding, and cells were ordered along the resulting trajectory using the first three diffusion components, with the ten pseudotime bins as the grouping variable. Candidate TFs were selected on the correlation between chromVAR motif activity and TF expression along this ordering (*p* < 0.05; correlation > 0.3), and TF–gene interactions with correlation > 0.2 were retained as descriptive selection criteria. The Pando and scMEGA networks were analysed separately and were not merged.

For the overlap analysis in **Fig. 3J**, the GRN-associated regulatory-region set comprised candidate regions carrying significant TF–target interactions in the Pando network (adjusted *p* < 0.05), and scMEGA-derived regions were not included. These regions therefore represent elements associated with the inferred time-course network rather than regions defined by differential accessibility. Overlapping intervals were merged to give a non-redundant set of 1,903 regulatory regions. Because the three compared sets derive from different region universes, intersection counts are reported descriptively rather than tested for enrichment. The inferred regulatory interactions also contributed to the integrated prioritization of candidate folate-responsive loci, including NKD1 and TGM2 (**Fig. 3K**; **Fig. S3F**).

### Gene set scoring and pathway activity along pseudotime

Gene set analysis used curated transcription factor modules representing key regulatory programs in the folate response, including signaling pathways and chromatin-associated regulators; module composition is listed in **Table S4C**. Per-cell gene set activity scores were computed using Seurat’s module scoring framework (AddModuleScore). To enable temporal interpretation, gene set scores were summarized across pseudotime bins, allowing comparison of pathway activity trends during progression of the folate response and linking transcriptional and regulatory pathway activity with continuous cell-state transitions.

### Publicly available ChIP-seq datasets (CistromeDB and GEO)

Publicly available transcription factor ChIP-seq datasets were obtained from CistromeDB (83,84) and GEO, including TCF7L2 (CistromeDB ID: 45714), CTNNB1 (CistromeDB ID: 53653; GEO: GSE53927; SRA: SAMN00009452), BCAT (CistromeDB IDs: 5454, 5455; GEO: SAMN00716123), and TCF4 (CistromeDB IDs: 5456, 54661) in colorectal cancer cell lines (all public datasets used in this study are summarized in **Table S7C**).

### Public H3K27ac dataset and super-enhancer analysis (GSE166254)

Publicly available H3K27ac ChIP–seq and matched RNA-seq data were obtained from GEO dataset GSE166254 (32), which profiles enhancer activity in primary colorectal carcinoma and matched adjacent normal colonic mucosa, enabling integrative analysis of enhancer landscapes, including super-enhancer identification, and their association with transcriptional outputs. Raw reads were quality-controlled with FastQC (v0.11.9) and adapter-trimmed with Trimmomatic (v0.39; SLIDINGWINDOW:4:20; MINLEN:25). Processed reads were aligned to hg38 using bwa-mem2 (v2.2.1) (85); SAM files were converted to sorted BAM with SAMtools (v1.16.1), and PCR duplicates were removed with Picard MarkDuplicates (v2.27.5 (86); REMOVE_DUPLICATES=true). H3K27ac-enriched regions were identified using MACS2 (v2.2.7.1) (87) in single-end mode with broad peak calling (--broad; q-value cutoff 0.01). Peak sets were merged across samples using BEDTools (88) to generate a unified set of regulatory regions, and read coverage was quantified from deduplicated BAM files using BAMscale (v1.0). Differential H3K27ac signal between tumor and normal samples was assessed using DESeq2 (v1.44.0); principal component analysis was used for quality assessment, and outlier samples (SRR13640019, SRR13640024) were excluded based on clustering. Differentially enriched regions defined tumor-associated enhancer activity changes, including candidate super-enhancers, and were integrated with matched RNA-seq to infer regulatory relationships. Super-enhancer (SE) analysis was performed using the ROSE algorithm (46) on tumor and matched normal H3K27ac data independently, stitching MACS2 peaks using a 12.5 kb stitching distance while excluding regions within 2.5 kb of transcription start sites, and ranking stitched regions by background-subtracted H3K27ac signal using the matched input as control to define SEs at the inflection point.

### Chromatin profiling (ATAC-seq and ChIP-seq alignment and peak calling)

ATAC–seq and ChIP–seq (transcription factor and H3K27ac) reads were quality-checked using FastQC (v0.11.9) and trimmed with Trimmomatic (v0.39) to remove adapter sequences (Nextera for ATAC–seq; TruSeq for ChIP–seq) and low-quality bases (SLIDINGWINDOW:4:20; MINLEN:25). Trimmed reads were aligned to hg38 using bwa-mem2 (v2.2.1), and SAM files were converted to sorted BAM with SAMtools (v1.16.1). PCR duplicates were removed using Picard MarkDuplicates (v2.27.5; REMOVE_DUPLICATES=true), and BAM files were indexed. Peaks were called from the deduplicated alignments using MACS2 (v2.2.7.1) in paired-end mode (-f BAMPE -g hs -q 0.01 --keep-dup all). Default narrow-peak calling was used for ATAC-seq and transcription factor ChIP-seq, whereas broad-peak calling (--broad) was used for H3K27ac ChIP-seq. The --keep-dup all option was used because PCR duplicates had already been removed with Picard. Peaks were pooled across samples and merged into a union peak set using BEDTools (sort followed by merge). Normalized coverage tracks and peak-level read counts were generated from the deduplicated BAM files using BAMscale (v1.0; scale and cov modes, respectively). Signal profiles and heatmaps were generated with deepTools using computeMatrix in reference-point mode (--referencePoint center, -b 2500 -a 2500), followed by plotHeatmap and plotProfile.

### Integration with external datasets and survival analysis

TCGA ATAC-seq normalized data (33) were obtained from the GDC repository. The log2-normalized peak matrix was imported into R (file code a544ed6f-4a27-4430-8f99-48b657da11fe) and filtered for COAD samples only (annotation file code 148b8d1e-a132-48f6-8e40-9ca96aa2f9b2). TCGA ATAC peaks were filtered to cases that overlapped with the open ATAC peaks identified in this study using the rtracklayer package (v1.68.0) (89). Sample clustering prior to survival analysis was performed using either COAD-specific peaks or any overlapping peak. Samples were partitioned into two groups by k-means clustering (seed = 1), and survival analysis was performed using the Surv and survfit functions of the survival package (v3.8-3) (90). Kaplan–Meier plots were generated using ggsurvplot from the survminer package (v0.5.2) (91). Signal heatmaps were generated using the ComplexHeatmap package (v2.24.1) (92).

### Colorectal cancer H3K27me3 ChIP-seq analysis

ChIP-seq data from colorectal cancer patients (44) were downloaded from SRA and GEO under accession GSE136888. H3K27me3 ChIP-seq samples were mapped to the hg38 genome assembly using the BWA mem aligner (v0.7.18) (93). Mapped reads were converted to BAM format and sorted using SAMtools (v1.21) (64). Duplicate reads were marked using Picard MarkDuplicates (v3.1.1) (86). Normalized coverage signals in bigWig format were generated using BAMscale (v0.0.9) (65) with a bin size of 50 bases. To identify differentially H3K27me3-enriched genomic windows between normal and tumor samples, the genome was binned into 100-kb windows, and the mean signal per bin per sample was quantified and stored in a matrix using rtracklayer (89). Signal-distribution boxplots for selected regions were generated with ggplot2, comparing conditions using the Wilcoxon rank-sum test. Chromosome- and locus-specific heatmaps were generated using the ComplexHeatmap package (v2.24.1) (92).

### Statistical analysis

Statistical analyses were performed using GraphPad Prism v8.0.1 (GraphPad Software) and R (v4.4.0). Unless otherwise stated in the figure legends, data are presented as mean ± SD from three independent biological replicates, defined as separately seeded and separately treated cultures. Tests were selected per assay and are named in the corresponding legend. Two-group comparisons of qRT–PCR and flow cytometry data used two-sided Student’s *t*-tests, whereas comparisons of IC_50_ values, colony counts and TIS-associated coverage used Welch’s two-sided *t*-test. WNT target-gene expression was assessed by one-sample *t*-test on log_2_ fold changes with Benjamini–Hochberg correction. Differential H3K27me3 and ChIP-seq signal comparisons used the Wilcoxon rank-sum test or, for paired regions, the paired Wilcoxon signed-rank test. Survival differences were assessed by log-rank test. Trajectory remodeling was evaluated by permutation testing with Benjamini–Hochberg correction and bootstrap confidence intervals. For bulk RNA-seq, differential expression was assessed using DESeq2 with Benjamini– Hochberg correction (adjusted *p* < 0.05). For single-cell multiome analyses, differential gene expression and chromatin accessibility were evaluated using the Wilcoxon rank-sum test with multiple-testing correction (FDR < 0.05). IC_50_ values were determined by nonlinear regression using a four-parameter logistic model.

### Quantification of trajectory-dependent epithelial state remodeling

Epithelial cell-state signatures were compiled from published single-cell and bulk transcriptomic studies of intestinal and colorectal epithelium; the candidate signatures and their source datasets are listed in **Table S6, S7C**. Signature gene symbols were matched case-insensitively to the features of the trajectory multiome object, and signatures with fewer than five matched genes were excluded, leaving 33 signatures for downstream analysis. Per-cell activity scores were computed for each retained signature using Seurat AddModuleScore on the RNA assay with default parameters, one call per signature. To quantify epithelial state remodeling during folate treatment, cells were ordered along the folate-responsive pseudotime trajectory and partitioned into ten equally populated pseudotime bins. Each epithelial cell-state signature score was standardized across all cells (z-score; zero mean and unit variance) to enable direct comparison across signatures with different dynamic ranges and gene-set sizes. For each signature, the mean standardized score was calculated within each pseudotime bin, yielding a trajectory of ten consecutive centroid values describing the temporal evolution of each epithelial program.

Trajectory-dependent remodeling was quantified using the Trajectory Remodeling Index (TRI), defined as the cumulative trajectory path length, where s*i* denotes the mean standardized signature score in pseudotime bin *i*. Unlike endpoint displacement, TRI quantifies the cumulative geometric distance travelled by each epithelial state program along pseudotime and therefore captures gradual, switch-like and other non-linear remodeling dynamics.

To distinguish sustained directional remodeling from stochastic trajectory fluctuations, we additionally calculated the Trajectory Directionality Index (TDI). First, the net trajectory displacement was calculated, where D represents the absolute difference between the first and last pseudotime centroids. TDI was then defined as the ratio of net displacement to total path length, where values approaching 1 indicate highly directed remodeling, whereas values approaching 0 indicate predominantly fluctuating trajectories.

Because TDI is defined as net displacement normalized by total path length (TRI), the two metrics are mathematically related rather than statistically independent: TRI quantifies the cumulative movement of a program, whereas TDI quantifies the fraction of that movement that is directionally coherent. The two indices were therefore interpreted as complementary descriptors of the same trajectory, cumulative versus net displacement, rather than as orthogonal variables.

For directional analyses, a signed TDI was used, in which the sign of the net displacement (s₁₀ − s₁) indicates whether remodeling proceeds toward the differentiated (positive) or stem-like (negative) pole of the epithelial state axis, while its magnitude retains the directionality value defined above. In this representation, each epithelial program is described by a single directional remodeling vector whose magnitude corresponds to TRI and whose orientation is given by the signed TDI.

Because the TDI ratio becomes unstable for near-stationary trajectories with negligible path length, TDI was interpreted only for signatures exceeding a minimum remodeling threshold (TRI ≥ 1.0); programs below this threshold were considered stationary and excluded from directional interpretation. Among the remaining programs, those with |signed TDI| ≥ 0.15 were classified as directionally remodeling, whereas programs with |signed TDI| < 0.15 were classified as non-directional (fluctuating). Of the 33 epithelial programs, 13 met the criteria for directional remodeling, whereas 20 were stationary or non-directional.

For two-dimensional epithelial state spaces (for example, DIFF versus SCL), trajectory remodeling was quantified analogously as the cumulative Euclidean distance travelled by successive pseudotime centroids, where x*i* and y*i* denote the standardized centroid coordinates in pseudotime bin *i*. This geometric metric directly quantifies movement through epithelial state space rather than pairwise similarity between epithelial programs. Trajectory remodeling metrics (TRI and TDI) were calculated for all epithelial cell-state signatures.

### Statistical analysis of trajectory remodeling

To determine whether the observed trajectory remodeling exceeded that expected by chance, pseudotime values were randomly permuted across cells (1,000 permutations) while signature scores were held fixed. Cells were subsequently reassigned to pseudotime bins using the original bin boundaries, preserving the number of cells per bin. TRI values were then recalculated for every epithelial program to generate a null distribution representing the absence of an ordered relationship between pseudotime and epithelial state. Empirical p values were calculated from 1,000 permutations and corrected for multiple testing across epithelial state signatures using the Benjamini–Hochberg procedure. Uncertainty of the Trajectory Directionality Index was estimated by resampling cells with replacement within each pseudotime bin (1,000 bootstrap replicates) and recomputing bin centroids and TDI for each replicate; 95% confidence intervals were defined as the 2.5th and 97.5th percentiles of the bootstrap distribution (percentile method), constrained to the [0,1] range of the index.

## Acknowledgements

We thank Nilay Sethi, MD, PhD (Dana-Farber Cancer Institute, Harvard Medical School) for valuable discussions and advice regarding the DER system. We are grateful to Dr. Zoltán Wiener (Semmelweis University, Budapest, Hungary) for providing patient-derived colon organoids and for his guidance and support related to organoid experiments. We also thank the staff of the HUN-REN TTK Flow Cytometry Core Facility, in particular Dr. György Várady and Dr. Edit Szabó, for their expert assistance with FACS analysis and cell sorting experiments. We are also grateful to Dr. Lóránt Székvölgyi (University of Debrecen, Faculty of Pharmacy, Department of Molecular and Nanopharmaceutics), Dr. Zsolt Czimmerer (HUN-REN Biological Research Centre) for critical reading of the manuscript and insightful comments.

## Author contributions

L.V. performed multiome and the majority of experimental work, and wrote and revised the manuscript. C.K. conducted computational analyses and contributed to manuscript revision. V.T. performed multiome and epigenetic experiments and contributed to writing and revision of the manuscript. A.Z. performed experimental work. A.M. carried out 3D culture and organoid experiments. A.K. contributed to manuscript revision. P.S. performed mass spectrometry analyses. I.C. contributed to manuscript revision and computational supervision. L.S.P. conducted epigenetic and TCGA data analysis and interpretation and contributed to manuscript revision. S.S. conceived the study, supervised the project, wrote and revised the manuscript. B.M. conceived the study, supervised the project, and contributed to manuscript revision.

## Data and code availability

All sequencing data generated in this study have been deposited in the Gene Expression Omnibus (GEO) under the following accession numbers, ATAC-seq, GSE336123; ChIP-seq, GSE336272; RNA-seq, GSE336267; and single-cell multiome, GSE336270. Previously published datasets used in this study are referenced in the Methods and summarized in **Table S7C**. All code used for data processing and analysis is available at https://github.com/kmcs111/folic-acid-analysis and will be made publicly accessible upon publication. All other materials are available from the corresponding authors upon reasonable request.

## Declaration of generative AI and AI-assisted technologies in the writing process

During the preparation of this work, the authors used Anthropic Claude and OpenAI ChatGPT (4o) to proofread the text, improve its readability and language, and assist with structural and editorial refinement. After using these tools, the authors reviewed and edited the content as needed and take full responsibility for the content of the publication.

## References

1. Longley DB, Harkin DP, Johnston PG. 5-fluorouracil: mechanisms of action and clinical strategies. Nat Rev Cancer 2003;3(5):330–8.

2. de Gramont A, Bosset JF, Milan C, Rougier P, Bouche O, Etienne PL, et al. Randomized trial comparing monthly low-dose leucovorin and fluorouracil bolus with bimonthly high-dose leucovorin and fluorouracil bolus plus continuous infusion for advanced colorectal cancer: a French intergroup study. J Clin Oncol 1997;15(2):808–15.

3. Locasale JW. Serine, glycine and one-carbon units: cancer metabolism in full circle. Nat Rev Cancer 2013;13(8):572–83.

4. Petrova B, Maynard AG, Wang P, Kanarek N. Regulatory mechanisms of one-carbon metabolism enzymes. J Biol Chem 2023;299(12):105457.

5. Reid MA, Dai Z, Locasale JW. The impact of cellular metabolism on chromatin dynamics and epigenetics. Nat Cell Biol 2017;19(11):1298–306.

6. Yang C, Zhang J, Liao M, Yang Y, Wang Y, Yuan Y, et al. Folate-mediated one-carbon metabolism: a targeting strategy in cancer therapy. Drug Discov Today 2021;26(3):817–25.

7. Lee Y, Vousden KH, Hennequart M. Cycling back to folate metabolism in cancer. Nat Cancer 2024;5(5):701–15.

8. Liu X, Ren B, Ren J, Gu M, You L, Zhao Y. The significant role of amino acid metabolic reprogramming in cancer. Cell Commun Signal 2024;22(1):380.

9. Thirion P, Michiels S, Pignon JP, Buyse M, Braud AC, Carlson RW, et al. Modulation of fluorouracil by leucovorin in patients with advanced colorectal cancer: an updated meta-analysis. J Clin Oncol 2004;22(18):3766–75.

10. QUASAR Collaborative Group. Comparison of fluorouracil with additional levamisole, higher-dose folinic acid, or both, as adjuvant chemotherapy for colorectal cancer: a randomised trial. Lancet 2000;355(9215):1588–96.

11. Schippinger W, Samonigg H, Schaberl-Moser R, Greil R, Thodtmann R, Tschmelitsch J, et al. A prospective randomised phase III trial of adjuvant chemotherapy with 5-fluorouracil and leucovorin in patients with stage II colon cancer. Br J Cancer 2007;97(8):1021–7.

12. Clevers H. The intestinal crypt, a prototype stem cell compartment. Cell 2013;154(2):274–84.

13. Fearon ER, Vogelstein B. A genetic model for colorectal tumorigenesis. Cell 1990;61(5):759–67.

14. Bala P, Rennhack JP, Aitymbayev D, Morris C, Moyer SM, Duronio GN, et al. Aberrant cell state plasticity mediated by developmental reprogramming precedes colorectal cancer initiation. Sci Adv 2023;9(13):eadf0927.

15. Barker N, van Es JH, Kuipers J, Kujala P, van den Born M, Cozijnsen M, et al. Identification of stem cells in small intestine and colon by marker gene Lgr5. Nature 2007;449(7165):1003–7.

16. Canellas-Socias A, Cortina C, Hernando-Momblona X, Palomo-Ponce S, Mulholland EJ, Turon G, et al. Metastatic recurrence in colorectal cancer arises from residual EMP1(+) cells. Nature 2022;611(7936):603–13.

17. Jackstadt R, van Hooff SR, Leach JD, Cortes-Lavaud X, Lohuis JO, Ridgway RA, et al. Epithelial NOTCH signaling rewires the tumor microenvironment of colorectal cancer to drive poor-prognosis subtypes and metastasis. Cancer Cell 2019;36(3):319–36 e7.

18. Moorman A, Benitez EK, Cambulli F, Jiang Q, Mahmoud A, Lumish M, et al. Progressive plasticity during colorectal cancer metastasis. Nature 2025;637(8047):947–54.

19. Mzoughi S, Schwarz M, Wang X, Demircioglu D, Ulukaya G, Mohammed K, et al. Oncofetal reprogramming drives phenotypic plasticity in WNT-dependent colorectal cancer. Nat Genet 2025;57(2):402–12.

20. Sato T, Vries RG, Snippert HJ, van de Wetering M, Barker N, Stange DE, et al. Single Lgr5 stem cells build crypt-villus structures in vitro without a mesenchymal niche. Nature 2009;459(7244):262–5.

21. Centonze A, Roura AJ, Novillo-Font M, Giordano C, Hernando-Momblona X, Llanses M, et al. A plastic EMP1+ to LGR5+ cell state conversion as a bypass to KRASG12D pharmacologic inhibition in metastatic colorectal cancer. Cancer Discov 2026;16(2):320–44 doi 10.1158/2159-8290.CD-25-0679.

22. Larue AEM, Atlasi Y. The epigenetic landscape in intestinal stem cells and its deregulation in colorectal cancer. Stem Cells 2024;42(6):509–25.

23. Perez-Gonzalez A, Bevant K, Blanpain C. Cancer cell plasticity during tumor progression, metastasis and response to therapy. Nat Cancer 2023;4(8):1063–82.

24. Alvarez-Varela A, Novellasdemunt L, Barriga FM, Hernando-Momblona X, Canellas-Socias A, Cano-Crespo S, et al. Mex3a marks drug-tolerant persister colorectal cancer cells that mediate relapse after chemotherapy. Nat Cancer 2022;3(9):1052–70.

25. Rehman SK, Haynes J, Collignon E, Brown KR, Wang Y, Nixon AML, et al. Colorectal cancer cells enter a diapause-like DTP state to survive chemotherapy. Cell 2021;184(1):226–42 e21.

26. Moss DY, Brown CN, Shaw AM, McCann C, Lewis N, Downs M, et al. Mitochondrial metabolism determines chemotherapy sensitivity in colorectal cancer. Nat Metab 2026;8(8):1751–71 doi 10.1038/s42255-026-01578-w.

27. Bartak BK, Nagy ZB, Szakallas N, Kalmar A, Farkas E, Banyai F, et al. Examination of genetic and epigenetic characteristics of patients with hyperhomocysteinemia following high-dose folic acid consumption. Nutrients 2025;17(13).

28. Zsigrai S, Kalmar A, Bartak BK, Nagy ZB, Szigeti KA, Valcz G, et al. Folic acid treatment directly influences the genetic and epigenetic regulation along with the associated cellular maintenance processes of HT-29 and SW480 colorectal cancer cell lines. Cancers (Basel) 2022;14(7).

29. Zsigrai S, Kalmar A, Nagy ZB, Bartak BK, Valcz G, Szigeti KA, et al. S-adenosylmethionine treatment of colorectal cancer cell lines alters DNA methylation, DNA repair and tumor progression-related gene expression. Cells 2020;9(8).

30. Spisak S, Chen D, Likasitwatanakul P, Doan P, Li Z, Bala P, et al. Identifying regulators of aberrant stem cell and differentiation activity in colorectal cancer using a dual endogenous reporter system. Nat Commun 2024;15(1):2230.

31. Akhtar-Zaidi B, Cowper-Sal-lari R, Corradin O, Saiakhova A, Bartels CF, Balasubramanian D, et al. Epigenomic enhancer profiling defines a signature of colon cancer. Science 2012;336(6082):736–9.

32. Zhou RW, Xu J, Martin TC, Zachem AL, He J, Ozturk S, et al. A local tumor microenvironment acquired super-enhancer induces an oncogenic driver in colorectal carcinoma. Nat Commun 2022;13(1):6041 doi 10.1038/s41467-022-33377-8.

33. Corces MR, Granja JM, Shams S, Louie BH, Seoane JA, Zhou W, et al. The chromatin accessibility landscape of primary human cancers. Science 2018;362(6413).

34. Cho YH, Ro EJ, Yoon JS, Mizutani T, Kang DW, Park JC, et al. 5-FU promotes stemness of colorectal cancer via p53-mediated WNT/beta-catenin pathway activation. Nat Commun 2020;11(1):5321.

35. Balsalobre A, Drouin J. Pioneer factors as master regulators of the epigenome and cell fate. Nat Rev Mol Cell Biol 2022;23(7):449–64.

36. Iwafuchi-Doi M, Zaret KS. Pioneer transcription factors in cell reprogramming. Genes Dev 2014;28(24):2679–92.

37. Vierbuchen T, Ling E, Cowley CJ, Couch CH, Wang X, Harmin DA, et al. AP-1 transcription factors and the BAF complex mediate signal-dependent enhancer selection. Mol Cell 2017;68(6):1067– 82 e12.

38. Taing L, Dandawate A, L’Yi S, Gehlenborg N, Brown M, Meyer CA. Cistrome Data Browser: integrated search, analysis and visualization of chromatin data. Nucleic Acids Res 2024;52(D1):D61– D6.

39. Chen L, Toke NH, Luo S, Vasoya RP, Fullem RL, Parthasarathy A, et al. A reinforcing HNF4-SMAD4 feed-forward module stabilizes enterocyte identity. Nat Genet 2019;51(5):777–85.

40. Becker WR, Nevins SA, Chen DC, Chiu R, Horning AM, Guha TK, et al. Single-cell analyses define a continuum of cell state and composition changes in the malignant transformation of polyps to colorectal cancer. Nat Genet 2022;54(7):985–95 doi 10.1038/s41588-022-01088-x.

41. Gilroy K, Najumudeen AK, Ridgway RA, Campbell AD, Sansom OJ, et al. MAPK-driven epithelial cell plasticity drives colorectal cancer therapeutic resistance. Nature 2026 doi 10.1038/s41586-025-09763-9.

42. Mentch SJ, Locasale JW. One-carbon metabolism and epigenetics: understanding the specificity. Ann N Y Acad Sci 2016;1363(1):91–8 doi 10.1111/nyas.12956.

43. Loi P, Schade AE, Rodriguez CL, Krishnan A, Perurena N, Nguyen VTM, et al. Epigenetic and oncogenic inhibitors cooperatively drive differentiation and kill KRAS-mutant colorectal cancers. Cancer Discov 2024;14(12):2430–49 doi 10.1158/2159-8290.CD-23-0866.

44. Orouji E, Raman AT, Singh AK, Sorokin A, Arslan E, Ghosh AK, et al. Chromatin state dynamics confers specific therapeutic strategies in enhancer subtypes of colorectal cancer. Gut 2022;71(5):938– 49 doi 10.1136/gutjnl-2020-322835.

45. Johnstone SE, Reyes A, Qi Y, Adriaens C, Hegazi E, Pelka K, et al. Large-scale topological changes restrain malignant progression in colorectal cancer. Cell 2020;182(6):1474–89 e23 doi 10.1016/j.cell.2020.07.030.

46. Whyte WA, Orlando DA, Hnisz D, Abraham BJ, Lin CY, Kagey MH, et al. Master transcription factors and mediator establish super-enhancers at key cell identity genes. Cell 2013;153(2):307–19 doi 10.1016/j.cell.2013.03.035.

47. Bajtai E, Kiss C, Bakos E, Lango T, Lovrics A, Schad E, et al. Therapy-induced senescence is a transient drug resistance mechanism in breast cancer. Mol Cancer 2025;24(1):128 doi 10.1186/s12943-025-02310-0.

48. Bailey SW, Ayling JE. The extremely slow and variable activity of dihydrofolate reductase in human liver and its implications for high folic acid intake. Proc Natl Acad Sci U S A 2009;106(36):15424–9 doi 10.1073/pnas.0902072106.

49. Guerreiro CS, Carmona B, Goncalves S, Carolino E, Fidalgo P, Brito M, et al. Risk of colorectal cancer associated with the C677T polymorphism in 5,10-methylenetetrahydrofolate reductase in Portuguese patients depends on the intake of methyl-donor nutrients. Am J Clin Nutr 2008;88(5):1413– 8 doi 10.3945/ajcn.2008.25877.

50. Sharp L, Little J, Brockton NT, Cotton SC, Masson LF, Haites NE, et al. Polymorphisms in the methylenetetrahydrofolate reductase (MTHFR) gene, intakes of folate and related B vitamins and colorectal cancer: a case-control study in a population with relatively low folate intake. Br J Nutr 2008;99(2):379–89 doi 10.1017/S0007114507801073.

51. Fardous AM, Beydoun S, James AA, Ma H, Cabelof DC, Unnikrishnan A, et al. The timing and duration of folate restriction differentially impacts colon carcinogenesis. Nutrients 2021;14(1):16 doi 10.3390/nu14010016.

52. Advanced Colorectal Cancer Meta-Analysis Project. Meta-analysis of randomized trials testing the biochemical modulation of fluorouracil by methotrexate in metastatic colorectal cancer. J Clin Oncol 1994;12(5):960–9 doi 10.1200/JCO.1994.12.5.960.

53. Bertino JR, Mini E, Fernandes DJ. Sequential methotrexate and 5-fluorouracil: mechanisms of synergy. Semin Oncol 1983;10(2 Suppl 2):2–5.

54. Cadman E, Heimer R, Davis L. Enhanced 5-fluorouracil nucleotide formation after methotrexate administration: explanation for drug synergism. Science 1979;205(4411):1135–7 doi 10.1126/science.472732.

55. Chavez-Hidalgo LP, Martin-Fernandez-de-Labastida S, Marques-Perez MdP, Arroyo-Izaga M. Influence of methyl donor nutrients as epigenetic regulators in colorectal cancer: a systematic review of observational studies. World J Gastroenterol 2023;29(7):1219–34 doi 10.3748/wjg.v29.i7.1219.

56. Ahmed D, Eide PW, Eilertsen IA, Danielsen SA, Eknaes M, Hektoen M, et al. Epigenetic and genetic features of 24 colon cancer cell lines. Oncogenesis 2013;2(9):e71 doi 10.1038/oncsis.2013.35.

57. Kawai K, Viars C, Arden K, Tarin D, Urquidi V, Goodison S. Comprehensive karyotyping of the HT-29 colon adenocarcinoma cell line. Genes Chromosomes Cancer 2002;34(1):1–8 doi 10.1002/gcc.10003.

58. Guinney J, Dienstmann R, Wang X, de Reynies A, Schlicker A, Soneson C, et al. The consensus molecular subtypes of colorectal cancer. Nat Med 2015;21(11):1350–6 doi 10.1038/nm.3967.

59. Nemeth K, Meszaros K, Szabo B, Butz H, Aranyi T, Szabo PT. A relative quantitation method for measuring DNA methylation and hydroxymethylation using guanine as an internal standard. Anal Methods 2021;13(39):4614–22 doi 10.1039/d1ay00897h.

60. Simon J, Kovacs M, Szabo PT. Finding the best column for polar basic analytes across reversed-phase and hydrophilic interaction liquid chromatography. Heliyon 2025;11(4):e42461 doi 10.1016/j.heliyon.2025.e42461.

61. Andrews S. FastQC: a quality control tool for high throughput sequence data. Babraham Bioinformatics; 2010.

62. Bolger AM, Lohse M, Usadel B. Trimmomatic: a flexible trimmer for Illumina sequence data. Bioinformatics 2014;30(15):2114–20 doi 10.1093/bioinformatics/btu170.

63. Dobin A, Davis CA, Schlesinger F, Drenkow J, Zaleski C, Jha S, et al. STAR: ultrafast universal RNA-seq aligner. Bioinformatics 2013;29(1):15–21 doi 10.1093/bioinformatics/bts635.

64. Danecek P, Bonfield JK, Liddle J, Marshall J, Ohan V, Pollard MO, et al. Twelve years of SAMtools and BCFtools. Gigascience 2021;10(2) doi 10.1093/gigascience/giab008.

65. Pongor LS, Gross JM, Vera Alvarez R, Murai J, Jang SM, Zhang H, et al. BAMscale: quantification of next-generation sequencing peaks and generation of scaled coverage tracks. Epigenetics Chromatin 2020;13(1):21 doi 10.1186/s13072-020-00343-x.

66. Liao Y, Smyth GK, Shi W. featureCounts: an efficient general purpose program for assigning sequence reads to genomic features. Bioinformatics 2014;30(7):923–30 doi 10.1093/bioinformatics/btt656.

67. Love MI, Huber W, Anders S. Moderated estimation of fold change and dispersion for RNA-seq data with DESeq2. Genome Biol 2014;15(12):550 doi 10.1186/s13059-014-0550-8.

68. Stephens M. False discovery rates: a new deal. Biostatistics 2017;18(2):275–94 doi 10.1093/biostatistics/kxw041.

69. 10x Genomics. Cell Ranger ARC. Pleasanton, CA; 2021.

70. Hao Y, Stuart T, Kowalski MH, Choudhary S, Hoffman P, Hartman A, et al. Dictionary learning for integrative, multimodal and scalable single-cell analysis. Nat Biotechnol 2024;42(2):293–304 doi 10.1038/s41587-023-01767-y.

71. Stuart T, Srivastava A, Madad S, Lareau CA, Satija R. Single-cell chromatin state analysis with Signac. Nat Methods 2021;18(11):1333–41 doi 10.1038/s41592-021-01282-5.

72. Granja JM, Corces MR, Pierce SE, Bagdatli ST, Choudhry H, Chang HY, et al. ArchR is a scalable software package for integrative single-cell chromatin accessibility analysis. Nat Genet 2021;53(3):403–11 doi 10.1038/s41588-021-00790-6.

73. Germain PL, Lun A, Garcia Meixide C, Macnair W, Robinson MD. Doublet identification in single-cell sequencing data using scDblFinder. F1000Res 2021;10:979 doi 10.12688/f1000research.73600.2.

74. Korsunsky I, Millard N, Fan J, Slowikowski K, Zhang F, Wei K, et al. Fast, sensitive and accurate integration of single-cell data with Harmony. Nat Methods 2019;16(12):1289–96 doi 10.1038/s41592-019-0619-0.

75. Cao J, Spielmann M, Qiu X, Huang X, Ibrahim DM, Hill AJ, et al. The single-cell transcriptional landscape of mammalian organogenesis. Nature 2019;566(7745):496–502 doi 10.1038/s41586-019-0969-x.

76. McLean CY, Bristor D, Hiller M, Clarke SL, Schaar BT, Lowe CB, et al. GREAT improves functional interpretation of cis-regulatory regions. Nat Biotechnol 2010;28(5):495–501 doi 10.1038/nbt.1630.

77. Wu T, Hu E, Xu S, Chen M, Guo P, Dai Z, et al. clusterProfiler 4.0: a universal enrichment tool for interpreting omics data. Innovation (Camb) 2021;2(3):100141 doi 10.1016/j.xinn.2021.100141.

78. Schep AN, Wu B, Buenrostro JD, Greenleaf WJ. chromVAR: inferring transcription-factor-associated accessibility from single-cell epigenomic data. Nat Methods 2017;14(10):975–8 doi 10.1038/nmeth.4401.

79. Tan G, Lenhard B. TFBSTools: an R/Bioconductor package for transcription factor binding site analysis. Bioinformatics 2016;32(10):1555–6 doi 10.1093/bioinformatics/btw024.

80. Fornes O, Castro-Mondragon JA, Khan A, van der Lee R, Zhang X, Richmond PA, et al. JASPAR 2020: update of the open-access database of transcription factor binding profiles. Nucleic Acids Res 2020;48(D1):D87–D92 doi 10.1093/nar/gkz1001.

81. Fleck JS, Jansen SMJ, Wollny D, Zenk F, Seimiya M, Jain A, et al. Inferring and perturbing cell fate regulomes in human brain organoids. Nature 2023;621(7978):365–72 doi 10.1038/s41586-022-05279-8.

82. Li Z, Nagai JS, Kuppe C, Kramann R, Costa IG. scMEGA: single-cell multi-omic enhancer-based gene regulatory network inference. Bioinform Adv 2023;3(1):vbad003 doi 10.1093/bioadv/vbad003.

83. Mei S, Qin Q, Wu Q, Sun H, Zheng R, Zang C, et al. Cistrome Data Browser: a data portal for ChIP-seq and chromatin accessibility data in human and mouse. Nucleic Acids Res 2017;45(D1):D658–D62 doi 10.1093/nar/gkw983.

84. 84. Zheng R, Wan C, Mei S, Qin Q, Wu Q, Sun H, et al. Cistrome Data Browser: expanded datasets and new tools for gene regulatory analysis. Nucleic Acids Res 2019;47(D1):D729–D35 doi 10.1093/nar/gky1094.

85. Vasimuddin M, Misra S, Li H, Aluru S. Efficient architecture-aware acceleration of BWA-MEM for multicore systems. IEEE International Parallel and Distributed Processing Symposium (IPDPS) 2019:314–24.

86. Broad Institute. Picard Toolkit. GitHub repository; 2019.

87. Zhang Y, Liu T, Meyer CA, Eeckhoute J, Johnson DS, Bernstein BE, et al. Model-based analysis of ChIP-seq (MACS). Genome Biol 2008;9(9):R137 doi 10.1186/gb-2008-9-9-r137.

88. Quinlan AR, Hall IM. BEDTools: a flexible suite of utilities for comparing genomic features. Bioinformatics 2010;26(6):841–2 doi 10.1093/bioinformatics/btq033.

89. Lawrence M, Gentleman R, Carey V. rtracklayer: an R package for interfacing with genome browsers. Bioinformatics 2009;25(14):1841–2 doi 10.1093/bioinformatics/btp328.

90. Therneau TM, Grambsch PM. Modeling survival data: extending the Cox model. Springer New York; 2000.

91. Kassambara A, Kosinski M, Biecek P, Fabian S. survminer: drawing survival curves using ggplot2. R package v0.5.2; 2026.

92. Gu Z, Eils R, Schlesner M. Complex heatmaps reveal patterns and correlations in multidimensional genomic data. Bioinformatics 2016;32(18):2847–9 doi 10.1093/bioinformatics/btw313.

93. Li H, Durbin R. Fast and accurate short read alignment with Burrows-Wheeler transform. Bioinformatics 2009;25(14):1754–60 doi 10.1093/bioinformatics/btp324.

