## Supplementary Figures for "One-carbon metabolism controls colorectal cancer cell state and chemosensitivity through chromatin remodeling"

### Supplementary Figure S1.

**Reporter-based epithelial fractionation and folate-dependent modulation of 5-fluorouracil sensitivity, clonogenic capacity, and therapy-induced senescence.**

*Related to Figure 1.*

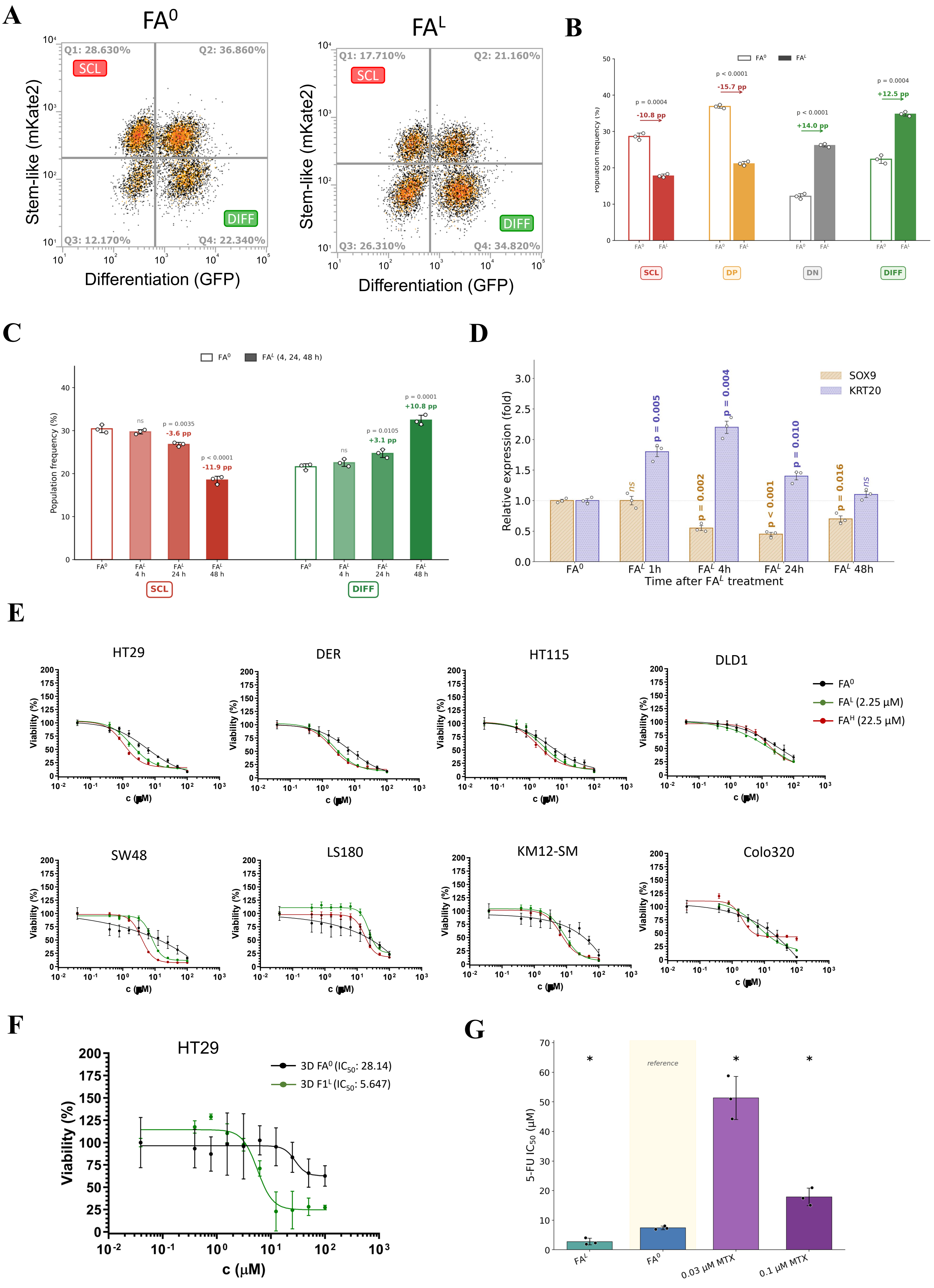
A
FA0
FAL
Q1: 28.630%
Q2: 36.860%
Q3: 12.170%
Q4: 22.340%
Q1: 17.710%
Q2: 21.160%
Q3: 26.310%
Q4: 34.820%
SCL
DIFF
Stem-like (mKate2)
Differentiation (GFP)
B
FA0
FAL
p = 0.0004
-10.8 pp
p < 0.0001
-15.7 pp
p < 0.0001
+14.0 pp
p = 0.0004
+12.5 pp
Population frequency (%)
SCL
DP
DN
DIFF
C
FA0
FAL (4, 24, 48 h)
ns
p = 0.0035
-3.6 pp
p < 0.0001
-11.9 pp
p = 0.0105
+3.1 pp
p = 0.0001
+10.8 pp
Population frequency (%)
FAL 4 h
FAL 24 h
FAL 48 h
SCL
DIFF
D
SOX9
KRT20
Relative expression (fold)
ns
p = 0.005
p = 0.002
p = 0.004
p < 0.001
p = 0.010
p = 0.016
ns
FA0
FAL 1h
FAL 4h
FAL 24h
FAL 48h
Time after FAL treatment
E
HT29
DER
HT115
DLD1
SW48
LS180
KM12-SM
Colo320
FA0
FAL (2.25 μM)
FAH (22.5 μM)
Viability (%)
c (μM)
F
HT29
3D FA0 (IC50: 28.14)
3D F1L (IC50: 5.647)
Viability (%)
c (μM)
G
reference
5-FU IC50 (μM)
FAL
FA0
0.03 μM MTX
0.1 μM MTX

**H**

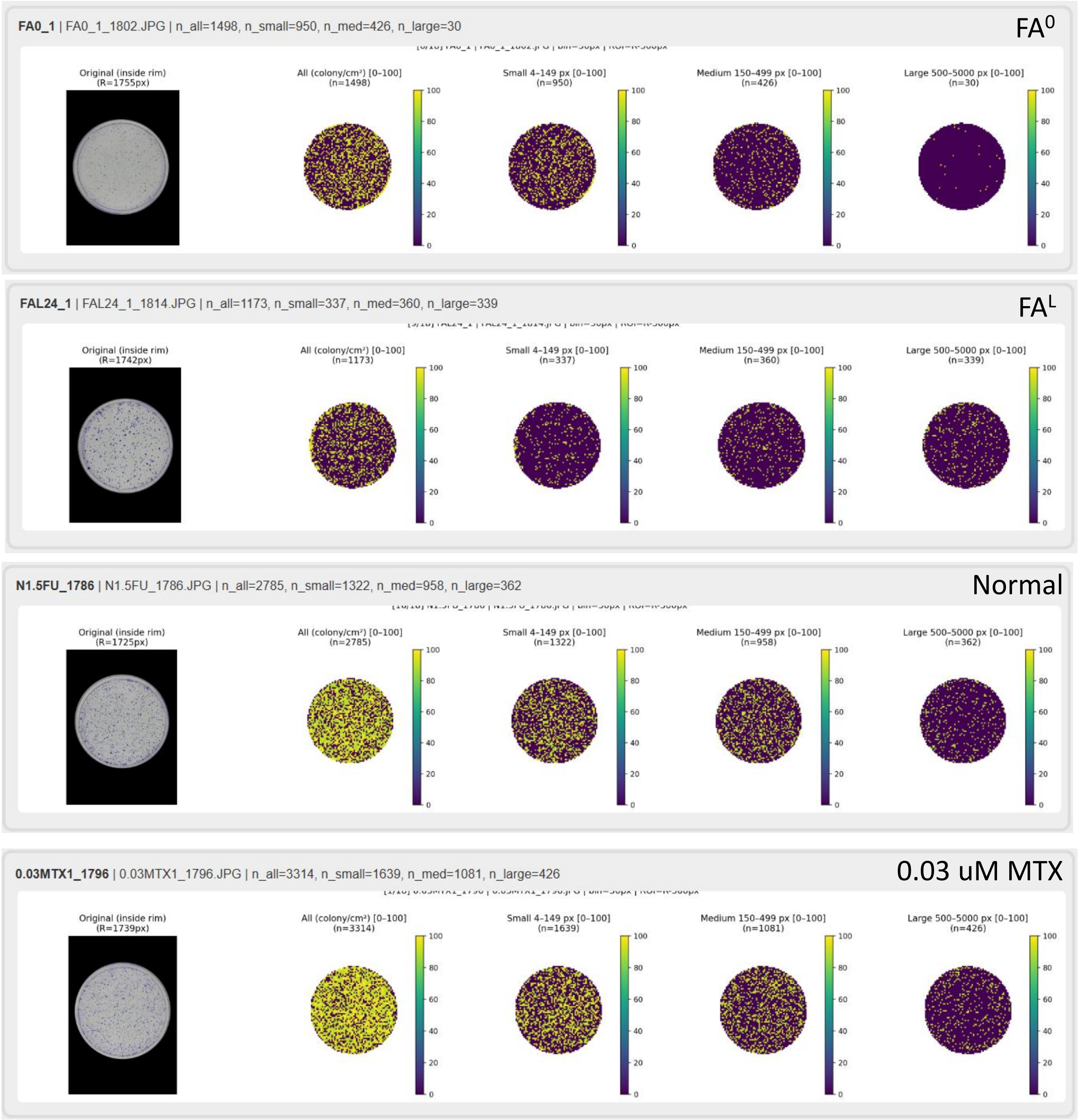
FA0_1 | FA0_1_1802.JPG | n_all=1498, n_small=950, n_med=426, n_large=30
FA0
Original (inside rim) (R=1755px)
All (colony/cm²) [0-100] (n=1498)
Small 4-149 px [0-100] (n=950)
Medium 150-499 px [0-100] (n=426)
Large 500-5000 px [0-100] (n=30)
FAL24_1 | FAL24_1_1814.JPG | n_all=1173, n_small=337, n_med=360, n_large=339
FAL
Original (inside rim) (R=1742px)
All (colony/cm²) [0-100] (n=1173)
Small 4-149 px [0-100] (n=337)
Medium 150-499 px [0-100] (n=360)
Large 500-5000 px [0-100] (n=339)
N1.5FU_1786 | N1.5FU_1786.JPG | n_all=2785, n_small=1322, n_med=958, n_large=362
Normal
Original (inside rim) (R=1725px)
All (colony/cm²) [0-100] (n=2785)
Small 4-149 px [0-100] (n=1322)
Medium 150-499 px [0-100] (n=958)
Large 500-5000 px [0-100] (n=362)
0.03MTX1_1796 | 0.03MTX1_1796.JPG | n_all=3314, n_small=1639, n_med=1081, n_large=426
0.03 uM MTX
Original (inside rim) (R=1739px)
All (colony/cm²) [0-100] (n=3314)
Small 4-149 px [0-100] (n=1639)
Medium 150-499 px [0-100] (n=1081)
Large 500-5000 px [0-100] (n=426)
100
80
60
40
20
0

**I**

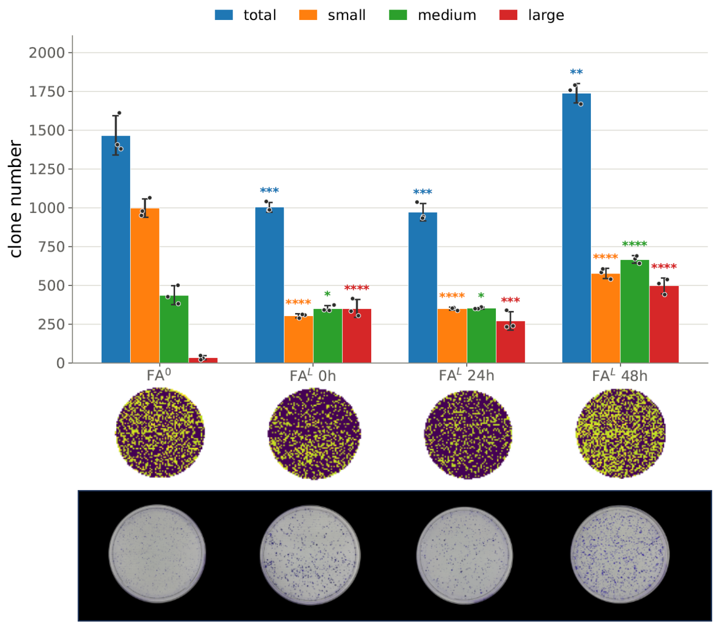
total
small
medium
large
clone number
2000
1750
1500
1250
1000
750
500
250
0
FA0
FAL 0h
FAL 24h
FAL 48h

**J**

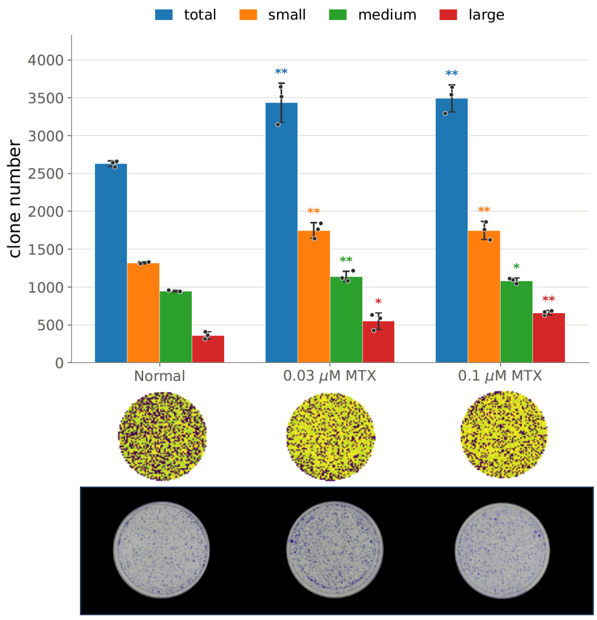
total
small
medium
large
clone number
4000
3500
3000
2500
2000
1500
1000
500
0
Normal
0.03 μM MTX
0.1 μM MTX

K

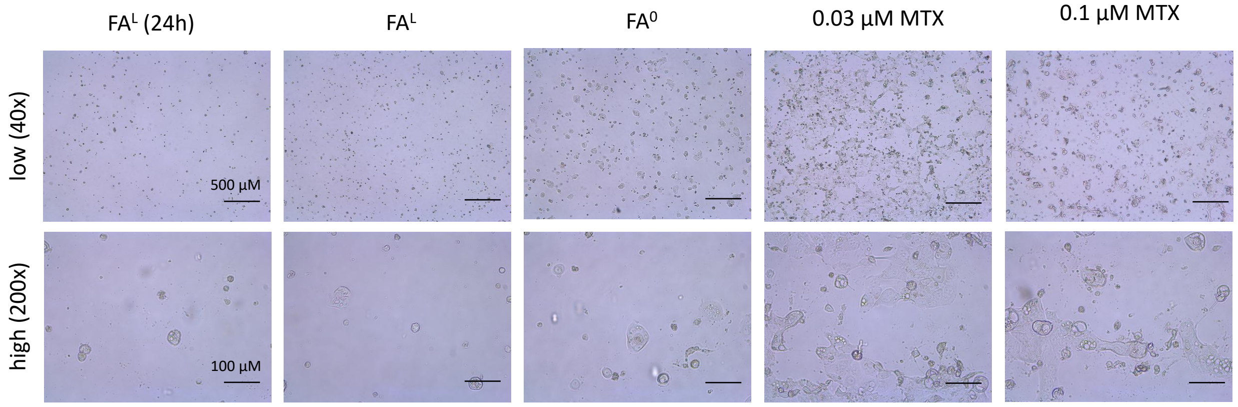
FAL (24h)
FAL
FA0
0.03 µM MTX
0.1 µM MTX
low (40x)
high (200x)
500 µM
100 µM

L

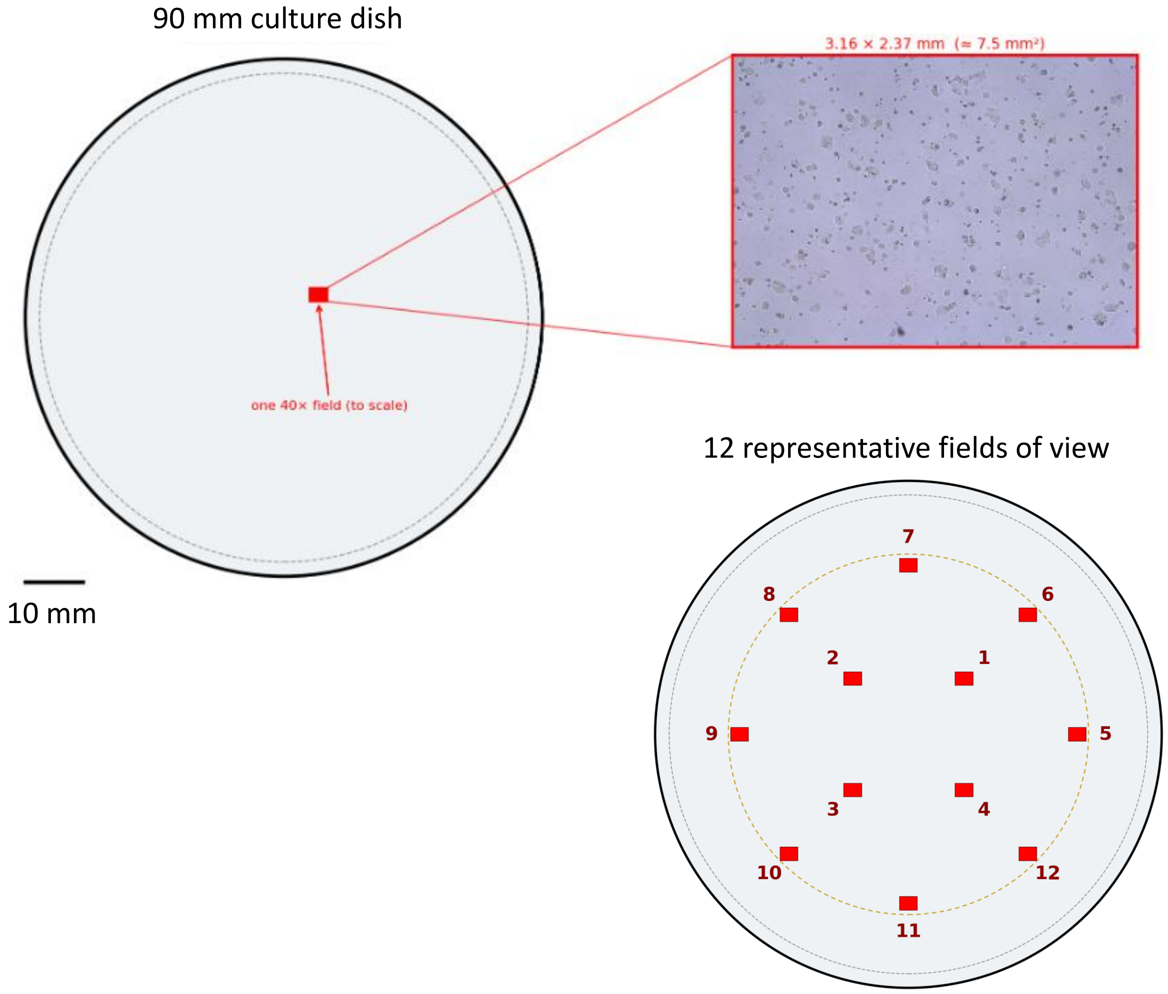
90 mm culture dish
3.16 × 2.37 mm (≈ 7.5 mm²)
one 40× field (to scale)
10 mm
12 representative fields of view
1
2
3
4
5
6
7
8
9
10
11
12

M

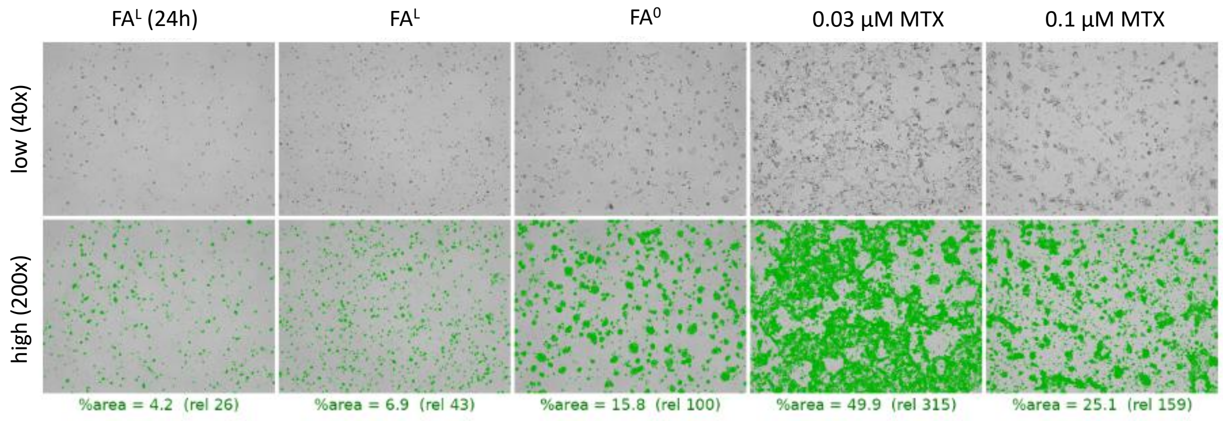
FAL (24h)
FAL
FA0
0.03 µM MTX
0.1 µM MTX
low (40x)
high (200x)
%area = 4.2 (rel 26)
%area = 6.9 (rel 43)
%area = 15.8 (rel 100)
%area = 49.9 (rel 315)
%area = 25.1 (rel 159)

**N**

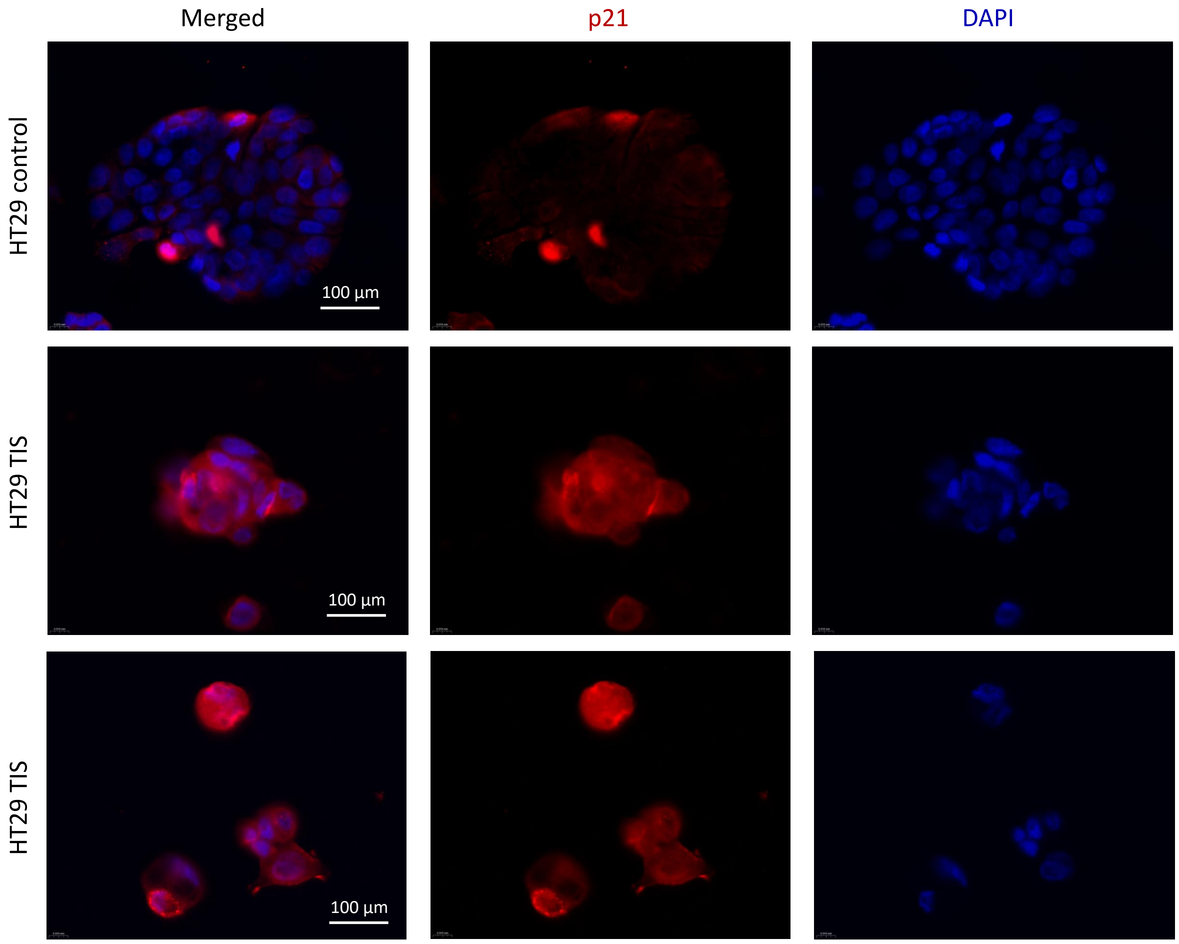

Merged
p21
DAPI
HT29 control
HT29 TIS
HT29 TIS
100 µm
100 µm
100 µm

**A,** Representative flow cytometry plots of HT29 DER cells cultured under $FA^0$ and $FA^L$ conditions. Quadrants indicate SCL ($mKate2^+/GFP^-$), double-positive, double-negative, and DIFF ($mKate2^-/GFP^+$) populations. Folate supplementation shifts the distribution of epithelial cell states toward the differentiated state.

**B,** Extended FACS quantification of the HT29 DER reporter response to folate, encompassing all four quadrant populations, stem-like (SCL; $mKate2^+/GFP^-$), double-positive (DP; $mKate2^+/GFP^+$), double-negative (DN; $mKate2^-/GFP^-$), and differentiated (DIFF; $mKate2^-/GFP^+$). Bar plot shows population frequencies under $FA^0$ (open bars) and $FA^L$ (filled bars). Data represent mean ± SD from three independent biological replicates; two-sided Student's t-test (SCL, −10.8 pp, $p = 0.0004$; DP, −15.7 pp, $p < 0.0001$; DN, +14.0 pp, $p < 0.0001$; DIFF, +12.5 pp, $p = 0.0004$). This analysis complements the two-population summary shown in **Fig. 1B**.

**C,** Time-resolved reporter response following folate restoration. After folate starvation, HT29 DER cells either remained under $FA^0$ conditions or received $FA^L$ supplementation (2.25 μM) for 4, 24 or 48 h, and SCL and DIFF population frequencies were determined by flow cytometry. Open bars indicate the $FA^0$ reference; filled bars indicate $FA^L$, with progressively darker shading denoting increasing exposure duration. Data are presented as mean ± SD ($n = 3$ biological replicates), with dots representing individual values. Comparisons with the corresponding $FA^0$ reference were performed using two-sided Student's *t*-tests. Percentage-point changes and $p$ values are indicated for significant comparisons; ns, not significant. No significant difference was detected at 4 h. Population frequencies shifted significantly from 24 h onwards, reaching −11.9 percentage points for SCL and +10.8 percentage points for DIFF at 48 h.

**D,** Time-course qRT-PCR analysis of *SOX9* and *KRT20* expression following folate supplementation. Transcript abundance was normalized to *ACTB* and expressed as fold change relative to the mean of $FA^0$ controls. Data are presented as mean ± SD ($n = 3$ biological replicates). Statistical significance was determined by two-sided Student's *t*-test versus $FA^0$.

**E,** 5-Fluorouracil (5-FU) dose–response curves across eight colorectal cancer cell lines (HT29, DER, HT115, DLD1, SW48, LS180, KM12-SM, and Colo320) cultured under folate-free ($FA^0$), low-folate ($FA^L$, 2.25 μM), and high-folate ($FA^H$, 22.5 μM) conditions. Viability is shown as a function of 5-FU concentration.

**F,** 5-FU dose–response of HT29 three-dimensional spheroids under $FA^0$ and $FA^L$ conditions, with the corresponding $IC_{50}$ values indicated.

**G,** 5-FU sensitivity of HT29 cells following folate manipulation or methotrexate (MTX; 0.03 or 0.1 μM) pretreatment. $FA^0$ served as the reference condition and is shaded. $IC_{50}$ values are presented as mean ± SD (n = 3 biological replicates), with dots showing individual values. Statistical significance was assessed against $FA^0$ by Welch's two-sided t-test; $^*p < 0.05$.

**H,** Representative automated colony segmentation workflow illustrating automated classification of total, small, medium, and large colonies under representative treatment conditions.

**I,** Quantification of colony number (total, small, medium, and large) across folate treatment conditions ($FA^0$ and $FA^L$ applied 0, 24, or 48 h before treatment).

**J,** Quantification of colony number (total, small, medium, and large) following methotrexate treatment (normal folate medium, 0.03 μM MTX, and 0.1 μM MTX).

**K,** Representative brightfield images of therapy-induced senescent (TIS) formation under the indicated treatment conditions at low (40x) and high (200x) magnification. Scale bars as indicated.

**L,** Schematic illustrating imaging coverage of a 90-mm culture dish and the sampling strategy (12 representative fields) used for TIS quantification.

**M,** Representative segmentation masks corresponding to the brightfield images shown in panel K, with quantified occupied area (%) indicated.

**N,** Representative immunofluorescence images of p21 (red) and DAPI (blue) in untreated control and TIS HT29 cells.

### Supplementary Figure S2.

**Bulk transcriptomic response to folate and single-cell multiome analysis of folate-induced state-specific remodeling.**
*Related to Figure 2.*

### BULK RNA HT29 48h

**A**

Short – FA$^L$ vs. FA$^0$ – 286 up / 256 down

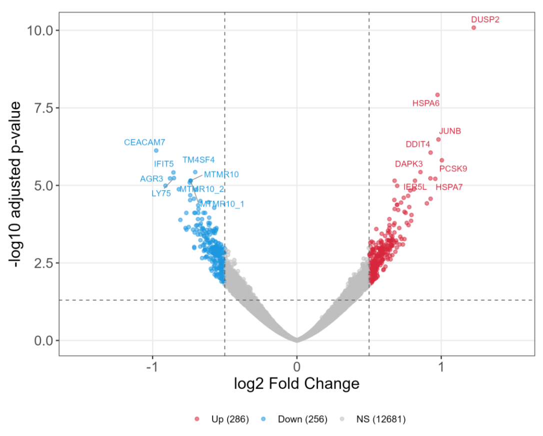


MSigDB Hallmark GSEA – Short FA$^L$ vs. FA$^0$

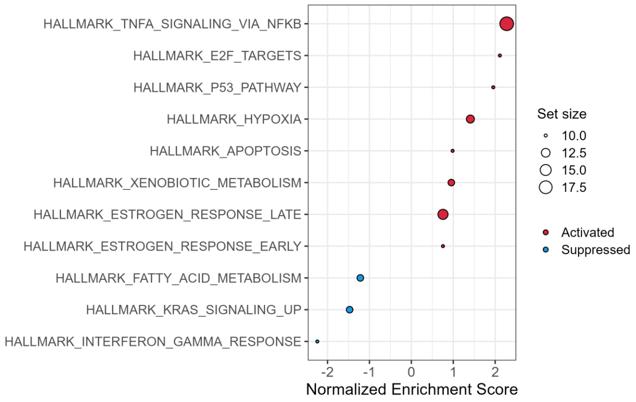


Short – FA$^H$ vs. FA$^0$ – 1478 up / 1827 down

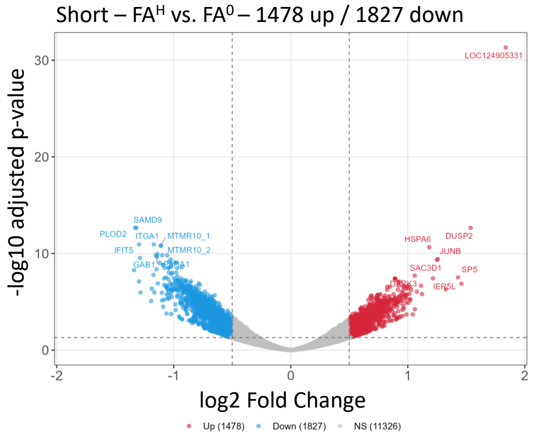


MSigDB Hallmark GSEA – Short FA$^H$ vs. FA$^0$

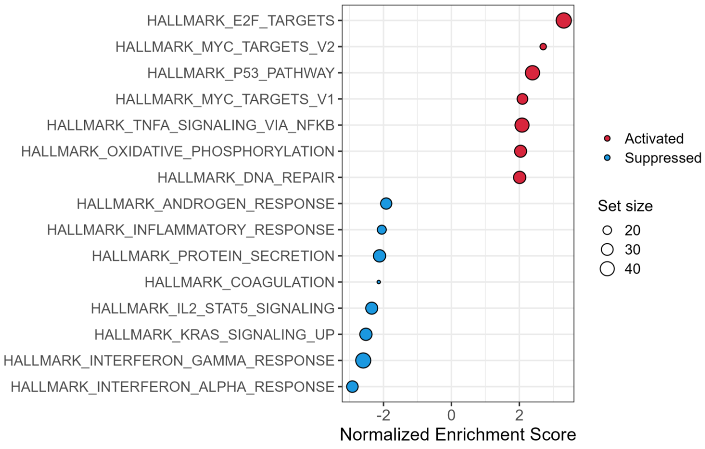


Short – FA$^H$ vs. FA$^L$ – 4 up / 2 down

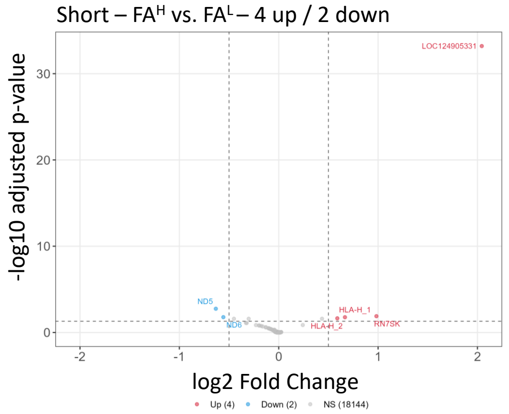


MSigDB Hallmark GSEA – Short FA$^H$ vs. FA$^L$

No significant gene sets

Short treatment: FA$^H$ vs. FA$^0$ ∩ FA$^L$ vs. FA$^0$

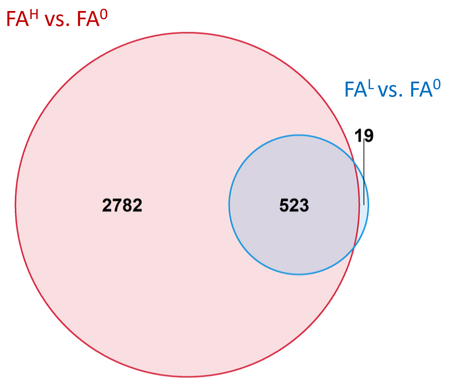


Short: genes shared between FA$^H$ vs. FA$^0$ & FA$^L$ vs. FA$^0$ (n=523) – sorted by FA$^H$ vs. FA$^0$ log2Fc

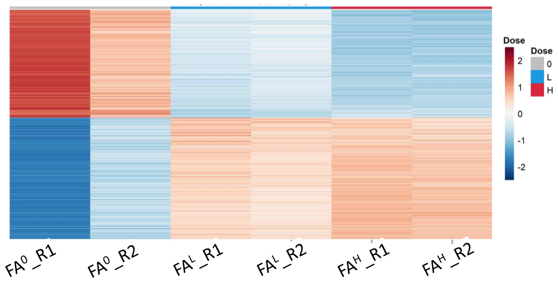

### BULK RNA HT29 14d

**B**

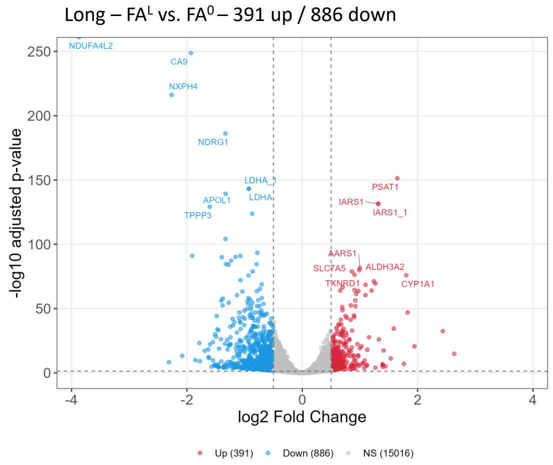


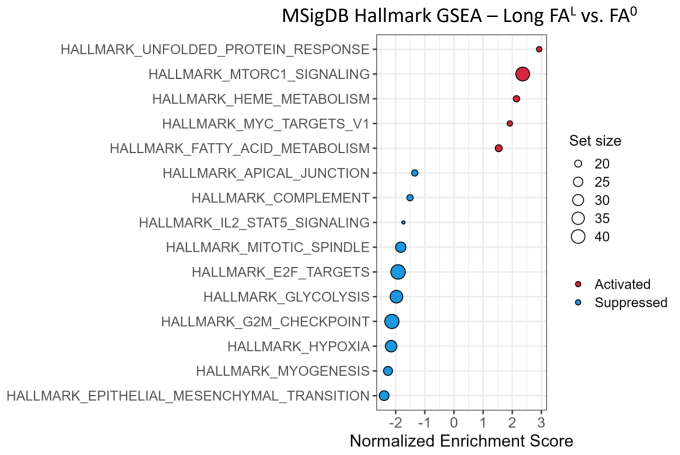


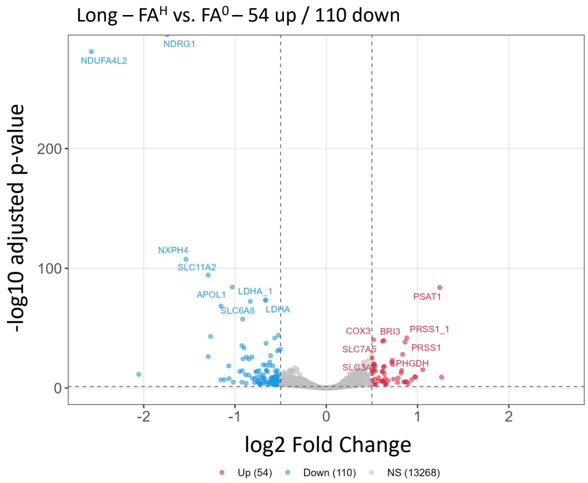


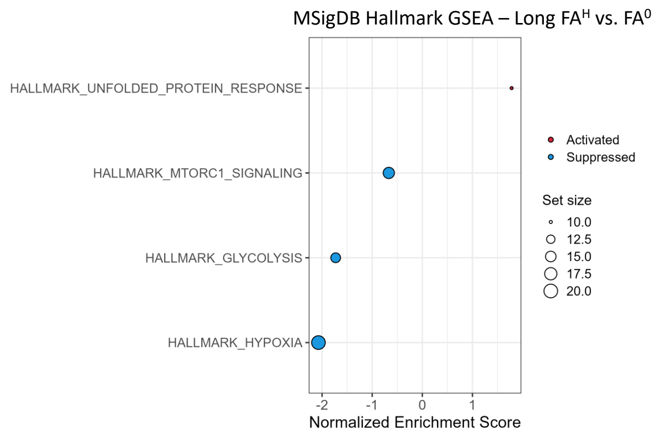


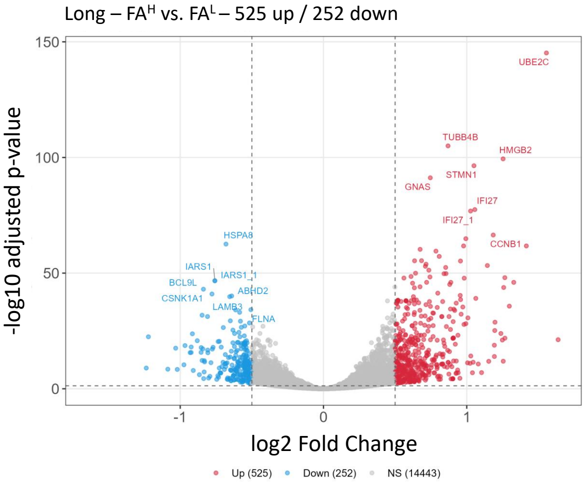


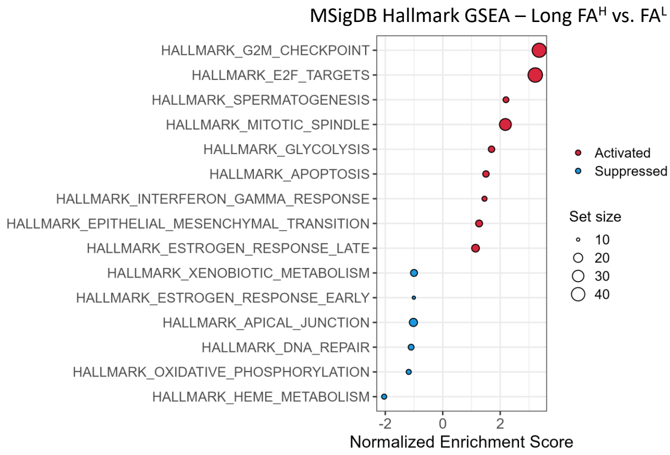


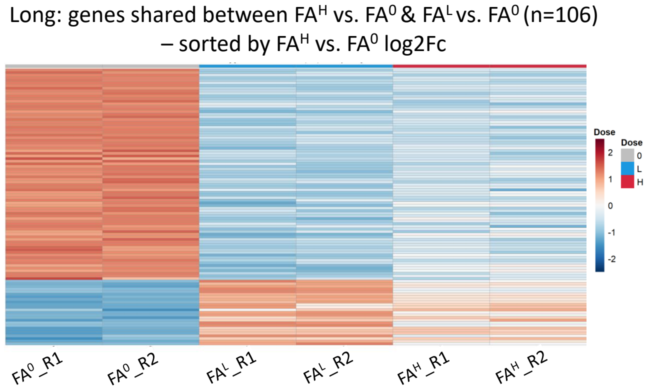


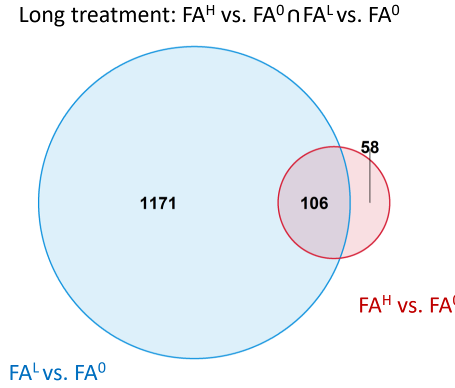

Supplementary analysis of bulk RNA-seq (HT29, folate dose x duration) – derived from Table S1D

**C**

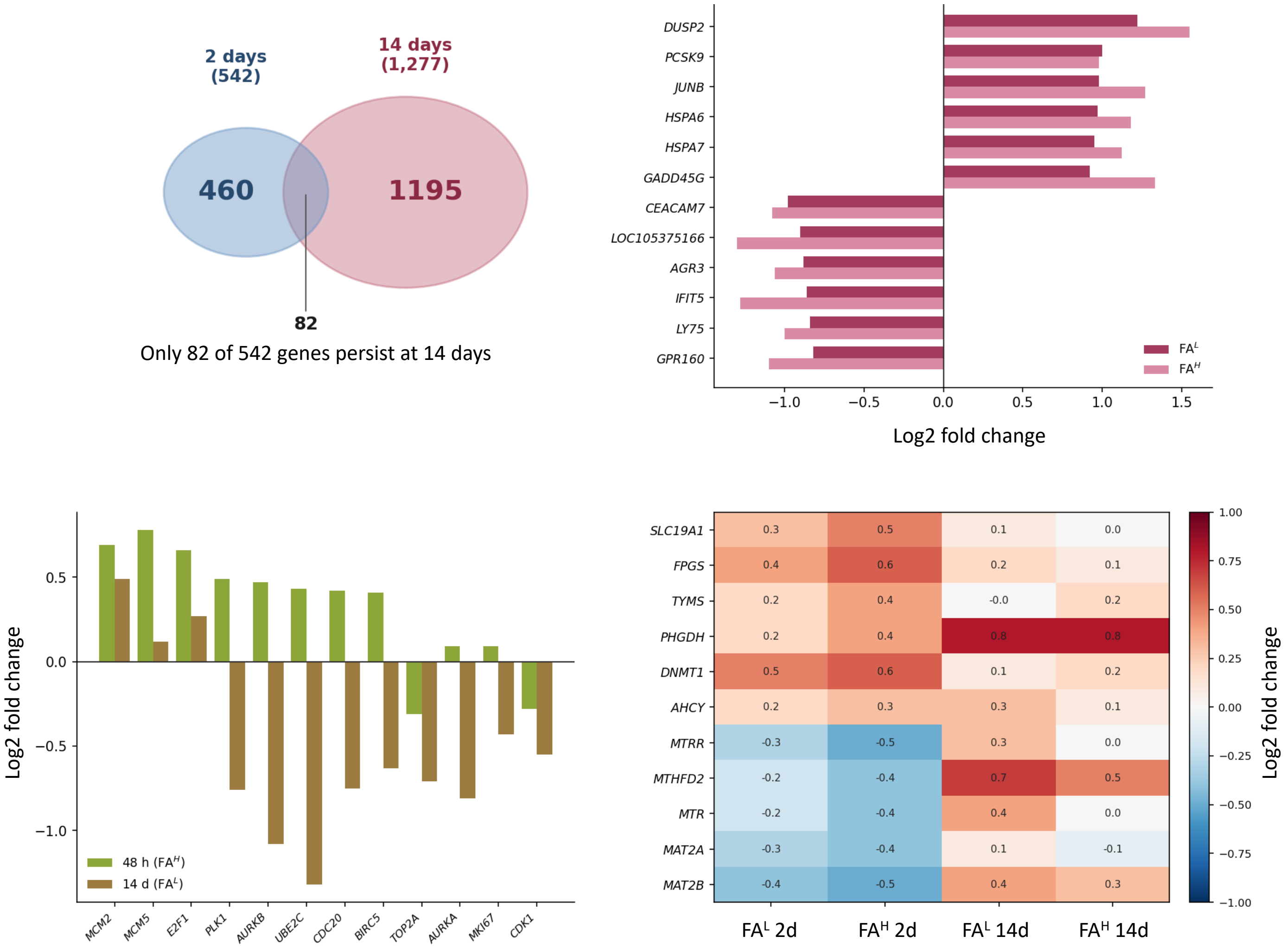

**D**

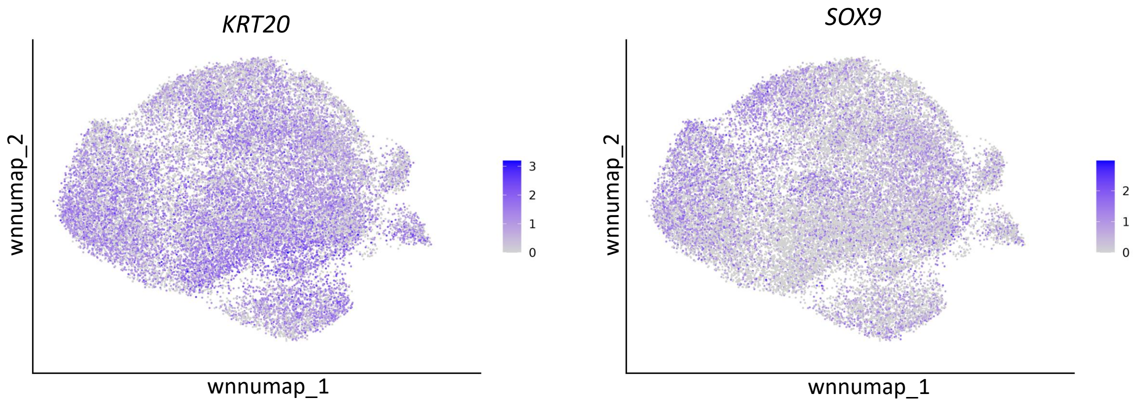


**E**

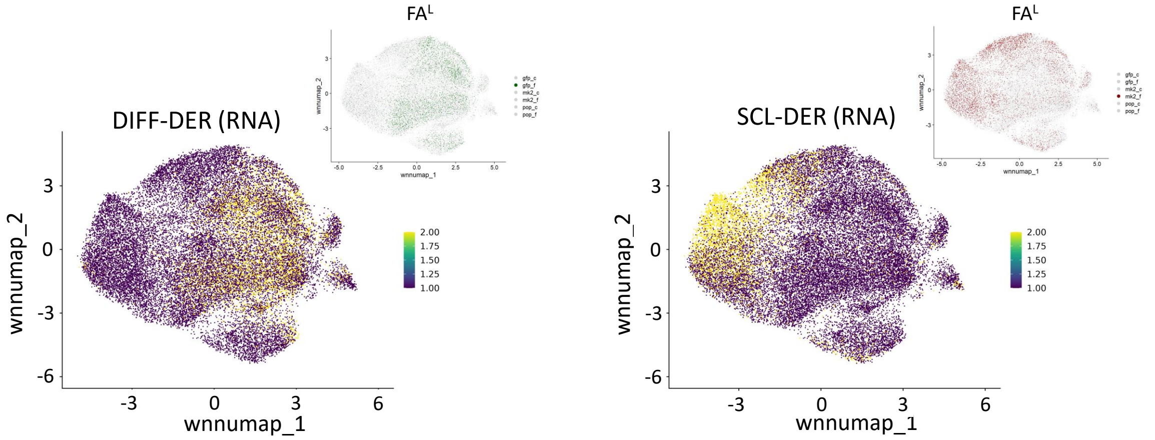


**F**

Sorted Fractions
(Fig.2. A,B)

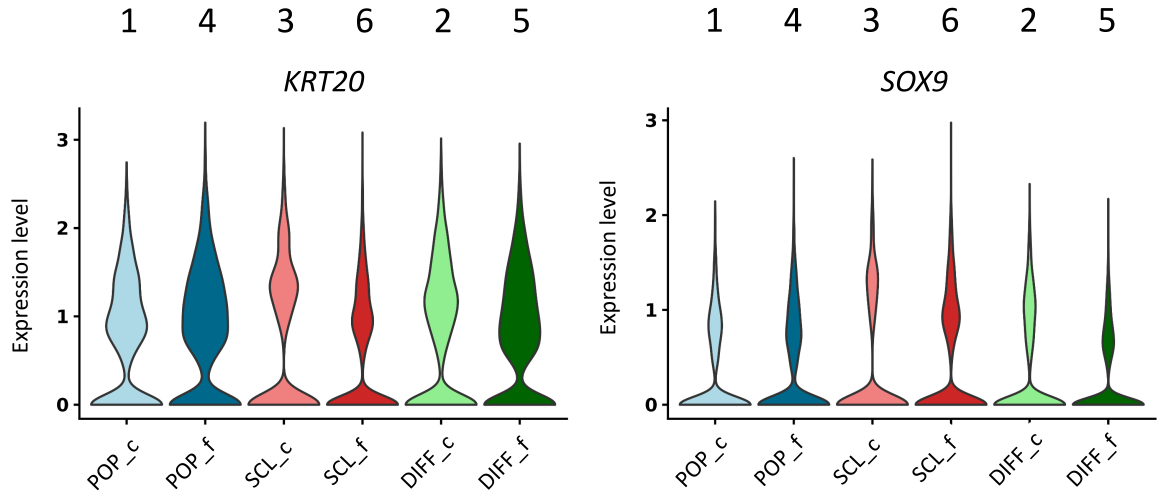


**G**

Sorted Fractions
(Fig.2. A, B)

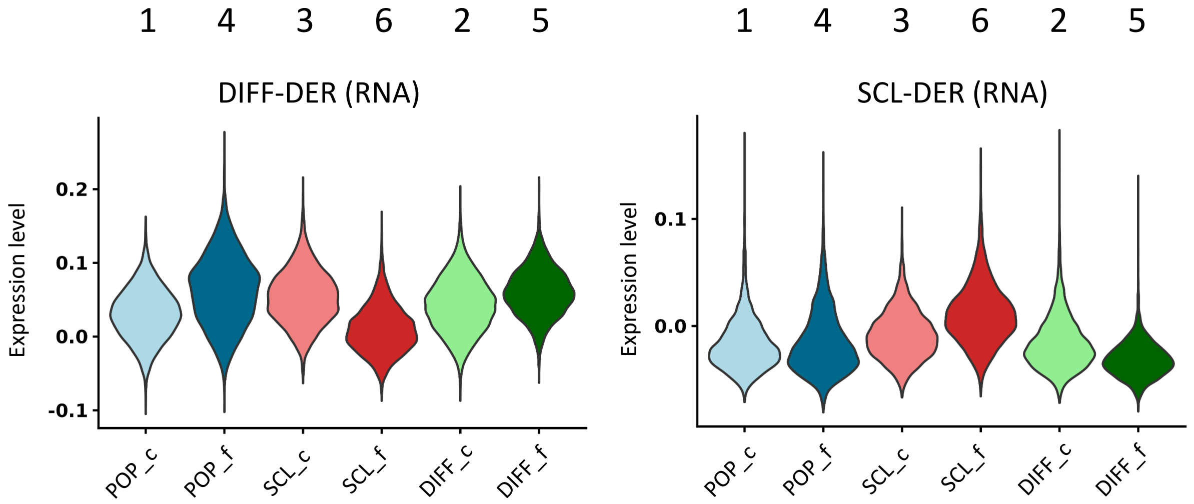

H

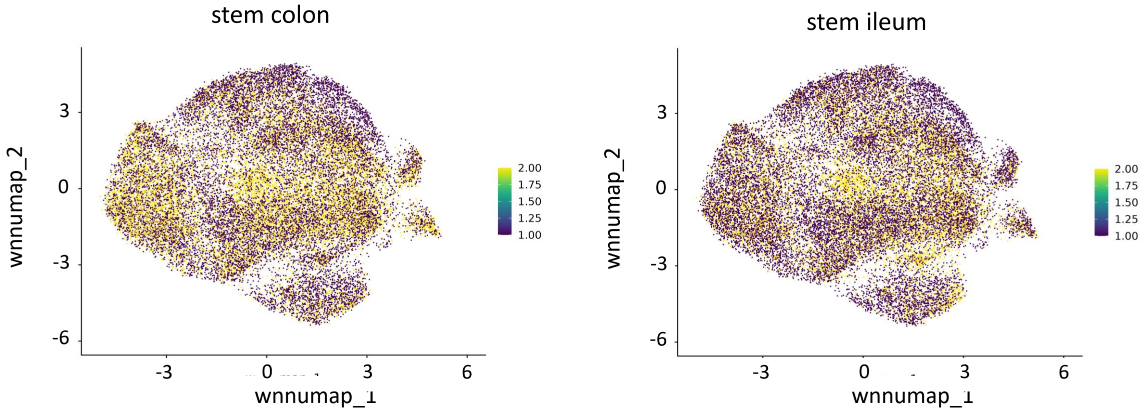

stem colon
stem ileum
wnnumap_2
wnnumap_1
3
0
-3
-6
6
2.00
1.75
1.50
1.25
1.00


I

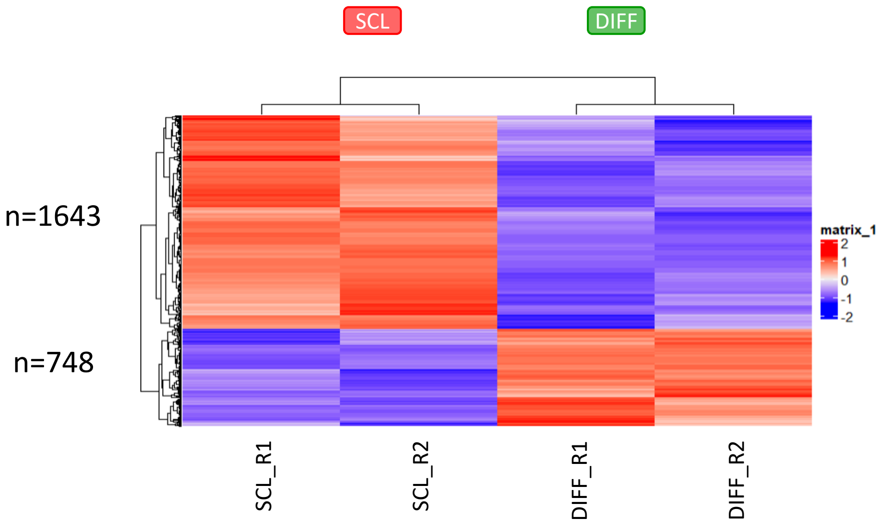

SCL
DIFF
n=1643
n=748
matrix_1
2
1
0
-1
-2
SCL_R1
SCL_R2
DIFF_R1
DIFF_R2


J

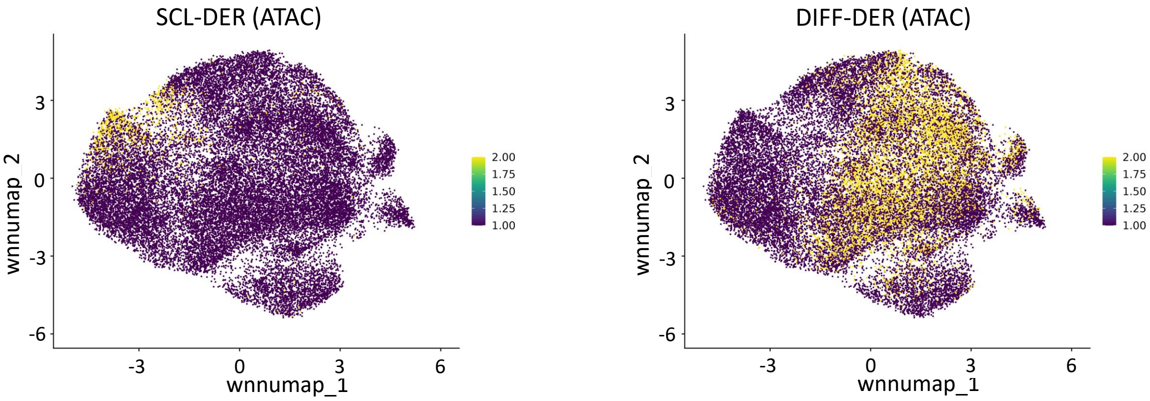

SCL-DER (ATAC)
DIFF-DER (ATAC)
wnnumap_2
wnnumap_1
3
0
-3
-6
6
2.00
1.75
1.50
1.25
1.00

K

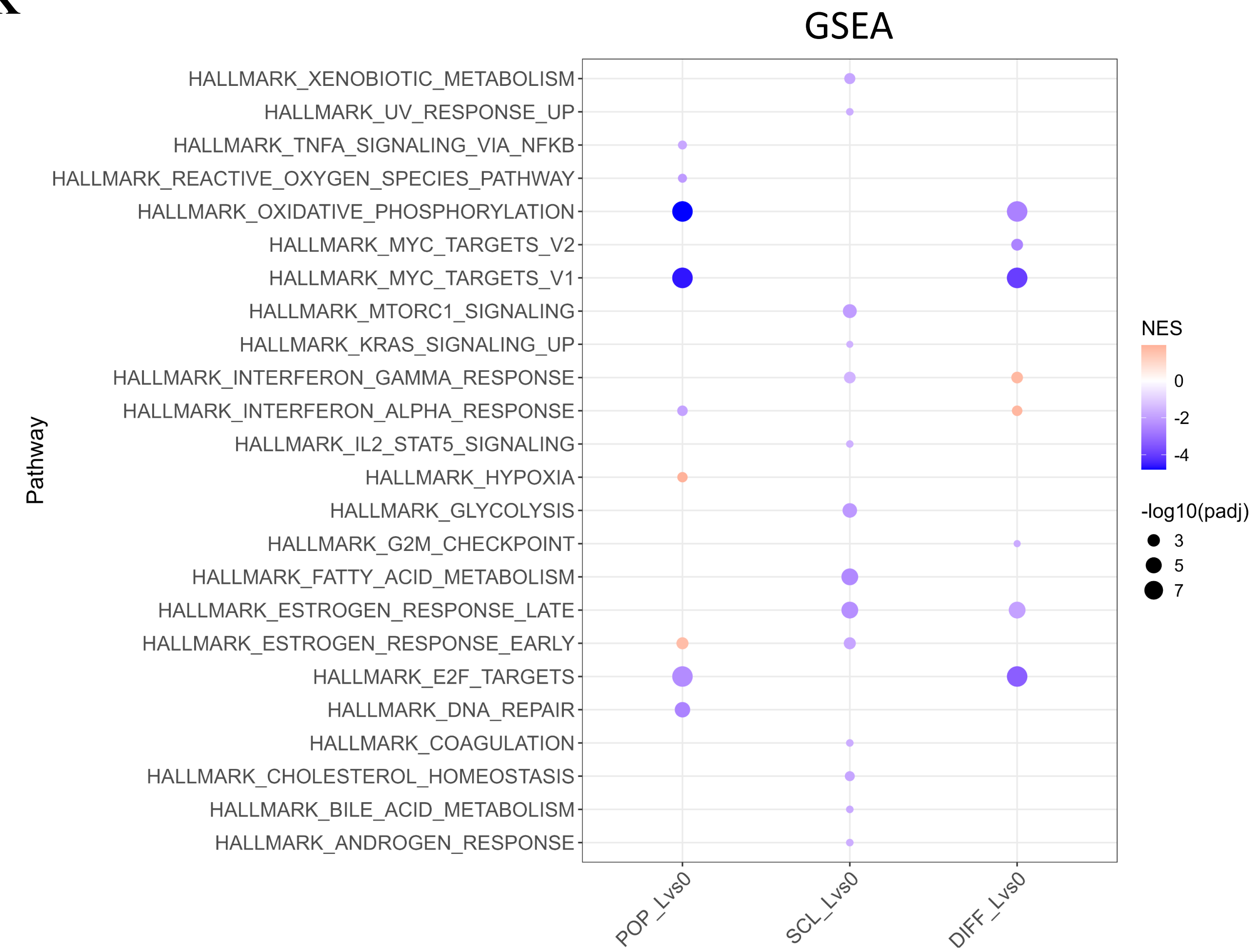
GSEA
HALLMARK_XENOBIOTIC_METABOLISM
HALLMARK_UV_RESPONSE_UP
HALLMARK_TNFA_SIGNALING_VIA_NFKB
HALLMARK_REACTIVE_OXYGEN_SPECIES_PATHWAY
HALLMARK_OXIDATIVE_PHOSPHORYLATION
HALLMARK_MYC_TARGETS_V2
HALLMARK_MYC_TARGETS_V1
HALLMARK_MTORC1_SIGNALING
HALLMARK_KRAS_SIGNALING_UP
HALLMARK_INTERFERON_GAMMA_RESPONSE
HALLMARK_INTERFERON_ALPHA_RESPONSE
HALLMARK_IL2_STAT5_SIGNALING
HALLMARK_HYPOXIA
HALLMARK_GLYCOLYSIS
HALLMARK_G2M_CHECKPOINT
HALLMARK_FATTY_ACID_METABOLISM
HALLMARK_ESTROGEN_RESPONSE_LATE
HALLMARK_ESTROGEN_RESPONSE_EARLY
HALLMARK_E2F_TARGETS
HALLMARK_DNA_REPAIR
HALLMARK_COAGULATION
HALLMARK_CHOLESTEROL_HOMEOSTASIS
HALLMARK_BILE_ACID_METABOLISM
HALLMARK_ANDROGEN_RESPONSE
Pathway
POP_Lvs0
SCL_Lvs0
DIFF_Lvs0
NES
0
-2
-4
-log10(padj)
3
5
7

L

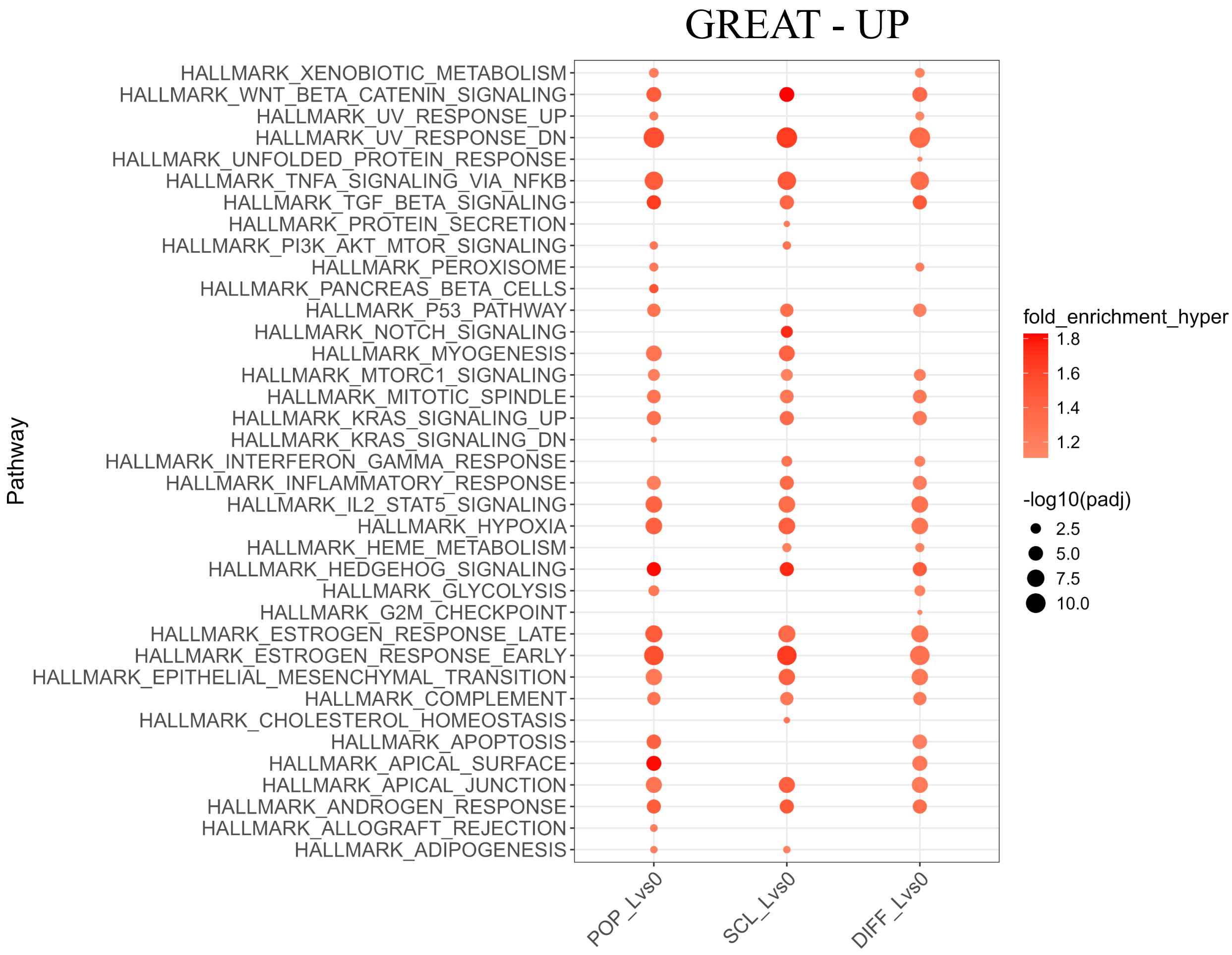
GREAT - UP
HALLMARK_XENOBIOTIC_METABOLISM
HALLMARK_WNT_BETA_CATENIN_SIGNALING
HALLMARK_UV_RESPONSE_UP
HALLMARK_UV_RESPONSE_DN
HALLMARK_UNFOLDED_PROTEIN_RESPONSE
HALLMARK_TNFA_SIGNALING_VIA_NFKB
HALLMARK_TGF_BETA_SIGNALING
HALLMARK_PROTEIN_SECRETION
HALLMARK_PI3K_AKT_MTOR_SIGNALING
HALLMARK_PEROXISOME
HALLMARK_PANCREAS_BETA_CELLS
HALLMARK_P53_PATHWAY
HALLMARK_NOTCH_SIGNALING
HALLMARK_MYOGENESIS
HALLMARK_MTORC1_SIGNALING
HALLMARK_MITOTIC_SPINDLE
HALLMARK_KRAS_SIGNALING_UP
HALLMARK_KRAS_SIGNALING_DN
HALLMARK_INTERFERON_GAMMA_RESPONSE
HALLMARK_INFLAMMATORY_RESPONSE
HALLMARK_IL2_STAT5_SIGNALING
HALLMARK_HYPOXIA
HALLMARK_HEME_METABOLISM
HALLMARK_HEDGEHOG_SIGNALING
HALLMARK_GLYCOLYSIS
HALLMARK_G2M_CHECKPOINT
HALLMARK_ESTROGEN_RESPONSE_LATE
HALLMARK_ESTROGEN_RESPONSE_EARLY
HALLMARK_EPITHELIAL_MESENCHYMAL_TRANSITION
HALLMARK_COMPLEMENT
HALLMARK_CHOLESTEROL_HOMEOSTASIS
HALLMARK_APOPTOSIS
HALLMARK_APICAL_SURFACE
HALLMARK_APICAL_JUNCTION
HALLMARK_ANDROGEN_RESPONSE
HALLMARK_ALLOGRAFT_REJECTION
HALLMARK_ADIPOGENESIS
Pathway
POP_Lvs0
SCL_Lvs0
DIFF_Lvs0
fold_enrichment_hyper
1.8
1.6
1.4
1.2
-log10(padj)
2.5
5.0
7.5
10.0

**M**

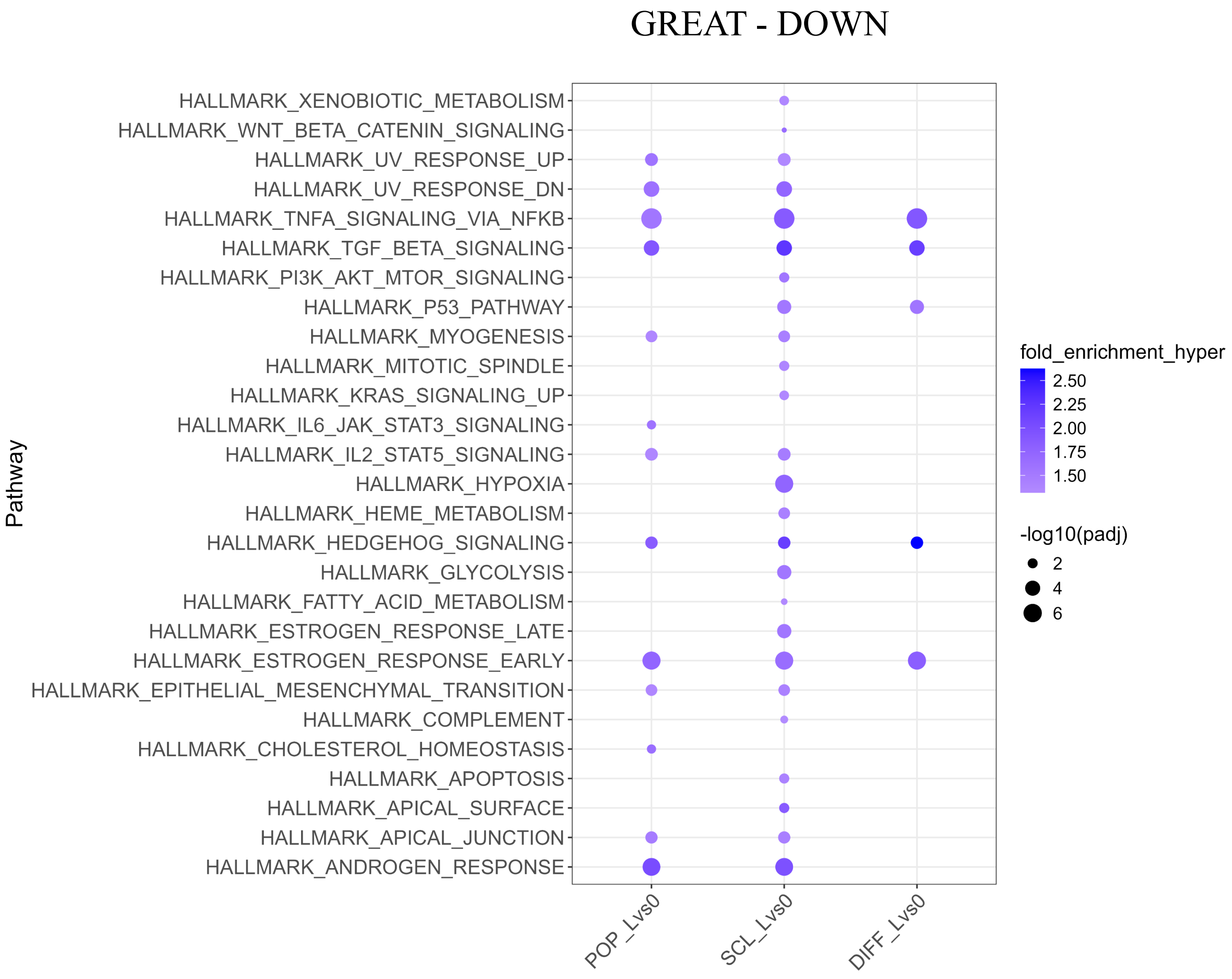
GREAT - DOWN
Pathway
HALLMARK_XENOBIOTIC_METABOLISM
HALLMARK_WNT_BETA_CATENIN_SIGNALING
HALLMARK_UV_RESPONSE_UP
HALLMARK_UV_RESPONSE_DN
HALLMARK_TNFA_SIGNALING_VIA_NFKB
HALLMARK_TGF_BETA_SIGNALING
HALLMARK_PI3K_AKT_MTOR_SIGNALING
HALLMARK_P53_PATHWAY
HALLMARK_MYOGENESIS
HALLMARK_MITOTIC_SPINDLE
HALLMARK_KRAS_SIGNALING_UP
HALLMARK_IL6_JAK_STAT3_SIGNALING
HALLMARK_IL2_STAT5_SIGNALING
HALLMARK_HYPOXIA
HALLMARK_HEME_METABOLISM
HALLMARK_HEDGEHOG_SIGNALING
HALLMARK_GLYCOLYSIS
HALLMARK_FATTY_ACID_METABOLISM
HALLMARK_ESTROGEN_RESPONSE_LATE
HALLMARK_ESTROGEN_RESPONSE_EARLY
HALLMARK_EPITHELIAL_MESENCHYMAL_TRANSITION
HALLMARK_COMPLEMENT
HALLMARK_CHOLESTEROL_HOMEOSTASIS
HALLMARK_APOPTOSIS
HALLMARK_APICAL_SURFACE
HALLMARK_APICAL_JUNCTION
HALLMARK_ANDROGEN_RESPONSE
POP_Lvs0
SCL_Lvs0
DIFF_Lvs0
fold_enrichment_hyper
2.50
2.25
2.00
1.75
1.50
-log10(padj)
2
4
6

**N**

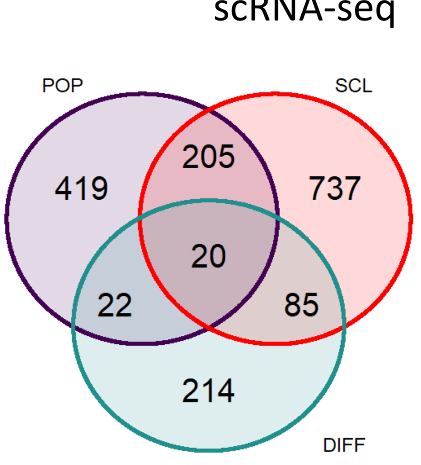
scRNA-seq
POP
SCL
DIFF
419
205
737
20
22
85
214

**O**

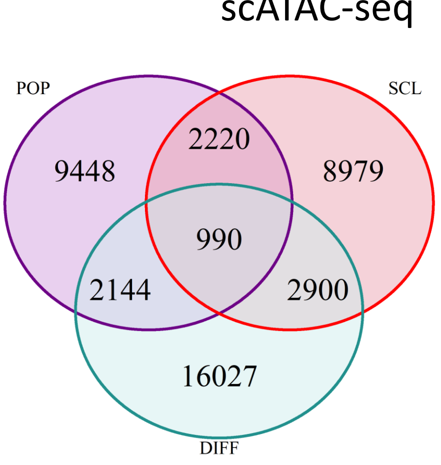
scATAC-seq
POP
SCL
DIFF
9448
2220
8979
990
2144
2900
16027

**P**

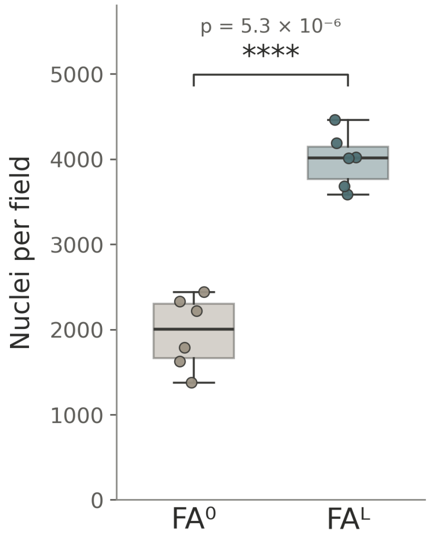
p = 5.3 × 10⁻⁶
****
Nuclei per field
5000
4000
3000
2000
1000
0
FA⁰
FAᴸ

Differential expression in panels A–C was defined using DESeq2 (adjusted $p < 0.05$, $|\log2FC| > 0.5$; red, upregulated; blue, downregulated). For all gene set enrichment analysis (GSEA) plots, dot size indicates gene-set size and color indicates positive (red) or negative (blue) normalized enrichment score against the MSigDB Hallmark collection.

**A,** Short-term (48 h) bulk transcriptomic response of HT29 cells to folate. Rows one to three show the $FA^L$ vs $FA^0$, $FA^H$ vs $FA^0$, and $FA^H$ vs $FA^L$ comparisons, each with the volcano plot on the left and the corresponding Hallmark GSEA on the right (286 up / 256 down, 1,478 up / 1,827 down, and 4 up / 2 down, respectively). In the third row, only six genes met the differential expression threshold and no Hallmark gene sets were significantly enriched, showing highly similar transcriptional responses to $FA^H$ and $FA^L$ at 48 h. Row four summarizes the short-term response, with the heatmap of top differentially expressed genes on the left (row-normalized z-scores; red, high; blue, low) and, on the right, the Venn diagram of significant DEGs shared between $FA^H$ vs $FA^0$ and $FA^L$ vs $FA^0$; 523 of 542 genes responsive to $FA^L$ were also differentially expressed in $FA^H$ (96.5% overlap).

**B,** Long-term (14-day) bulk transcriptomic response of HT29 cells to folate, shown in the same layout as panel A. Rows one to three show the $FA^L$ vs $FA^0$, $FA^H$ vs $FA^0$, and $FA^H$ vs $FA^L$ comparisons (391 up / 886 down, 54 up / 110 down, and 525 up / 252 down, respectively), each with the volcano plot on the left and the corresponding Hallmark GSEA on the right. Compared with the 48 h time point, substantially fewer genes were differentially expressed following prolonged $FA^H$ treatment, whereas a substantial transcriptional difference emerged between $FA^H$ and $FA^L$. Row four summarizes the long-term response, with the heatmap of top differentially expressed genes on the left and, on the right, the Venn diagram of significant DEGs shared between $FA^H$ vs $FA^0$ and $FA^L$ vs $FA^0$ (106 shared), with reduced overlap relative to the 48 h comparison (panel A).

**C, Supplementary analyses of the bulk RNA-seq dataset across folate dose and treatment duration, derived from Table S2A.** Upper left, temporal overlap of differentially expressed genes between short-term (48h) and long-term (14d) low-dose ($FA^L$ vs $FA^0$) treatment; of the 542 early-responsive genes, only 82 remained differentially expressed at 14 days (460 early-specific; 1,195 late-specific). Upper right, representative consistently folate-responsive genes at 48 h shown for both $FA^L$ and $FA^H$ (log2 fold change); upregulated genes include DUSP2, PCSK9, JUNB, HSPA6, HSPA7, and GADD45G, and downregulated genes include CEACAM7 and AGR3. Bottom left, cell-cycle and proliferation-associated genes across conditions; replication-associated genes (MCM2, MCM5, E2F1) were induced at 48 h, whereas several mitotic regulators (TOP2A, AURKA, CDC20) showed reduced expression after prolonged treatment. Bottom right, one-carbon metabolism and methylation-associated genes across folate dose and duration (log2 fold change); at the early time point folate exposure was accompanied by coordinated, modest upregulation of SLC19A1, FPGS, DNMT1, and AHCY with downregulation of MAT2A, MAT2B, and MTHFD2, whereas at 14 days several of these genes showed altered patterns relative to the early response, including reversal of MTHFD2 expression and stronger induction of PHGDH.

**D,** *KRT20* (left) and *SOX9* (right) expression projected onto the integrated single-cell multiome manifold. *KRT20* localizes predominantly to differentiated epithelial regions, whereas *SOX9* marks stem-like domains, highlighting the epithelial state organization recapitulated by the reporter system.

**E,** DIFF-DER (RNA) and SCL-DER (RNA) transcriptional programs, derived from bulk reporter-sorted HT29 populations (30), projected onto the integrated epithelial manifold (insets show the corresponding $FA^L$-treated reporter-positive populations). DIFF-DER (RNA) localizes to differentiated domains and SCL-DER (RNA) to stem-like domains, consistent with the epithelial organization defined by endogenous *KRT20* and *SOX9* expression in panel D.

**F,** *KRT20* and *SOX9* expression across the six FACS-sorted epithelial fractions (**Fig. 2A,B**). *KRT20* and *SOX9* show reciprocal expression across differentiated and stem-like fractions, with the clearest separation observed in folate-treated DIFF (fraction 5) and SCL (fraction 6), whereas POP fractions display more heterogeneous expression.

**G,** DIFF-DER (RNA) and SCL-DER (RNA) program enrichment across the six FACS-sorted epithelial fractions. DIFF-DER (RNA) scores are highest in differentiated fractions, whereas SCL-DER (RNA) scores are highest in stem-like fractions. Consistent with the bulk DER RNA-seq reference datasets, enrichment is strongest in folate-treated DIFF (fraction 5) and SCL (fraction 6).

**H,** Colon and ileum stem cell-associated transcriptional programs mapped onto the integrated epithelial manifold.

**I,** Genome-wide chromatin-accessibility heatmap of the SCL-DER (ATAC) and DIFF-DER (ATAC) signatures, derived from bulk ATAC-seq profiling of top-2% $mKate2^+$ (SCL) and $GFP^+$ (DIFF) sorted DER populations. Differential analysis identified 1,643 peaks preferentially accessible in the SCL state SCL-DER (ATAC) and 748 in the DIFF state DIFF-DER (ATAC) (row-normalized z-scores; red, high; blue, low), shown across two independent SCL and DIFF replicates.

**J,** DIFF-DER (ATAC) and SCL-DER (ATAC) accessibility signatures projected onto the epithelial manifold. DIFF-DER (ATAC)-associated regions localize within differentiated domains and SCL-DER (ATAC)-associated regions within stem-like domains.

**K,** GSEA of folate-responsive transcriptional changes between $FA^L$ and $FA^0$ conditions across the POP, SCL, and DIFF states, summarized as normalized enrichment scores (NES) for MSigDB Hallmark pathways.

**L,** GREAT enrichment analysis of folate-responsive gained chromatin-accessibility regions across POP, SCL, and DIFF states, summarized as hypergeometric fold enrichment for MSigDB Hallmark pathways.

**M,** GREAT enrichment analysis of folate-responsive lost chromatin-accessibility regions across POP, SCL, and DIFF states, summarized as hypergeometric fold enrichment for MSigDB Hallmark pathways.

**N,** Three-way Venn diagram of state-specific differentially expressed genes across POP, SCL, and DIFF fractions (scRNA-seq).

**O,** Three-way Venn diagram of state-specific differential chromatin-accessibility regions across POP, SCL, and DIFF fractions (scATAC-seq).

**P,** Population-level cell density following folate restoration. HT29 cells were seeded at 1,000 cells per well and cultured for five days under $FA^0$ or $FA^L$ conditions. Nuclei were counted by high-content imaging across four fixed fields per well, the sampling format used for this assay, and values are expressed as the mean number of nuclei per field. Because the four fields sample only part of the circular well, values reflect relative cell density rather than absolute cell number. Folate restoration increased cell density approximately twofold ($n$ = 6 wells per condition). Boxes show the interquartile range and median, whiskers the range, and dots individual wells. Statistical significance was assessed by Welch's two-sided $t$-test; ****$p < 0.0001$.

### Supplementary Figure S3.

**Trajectory-resolved chromatin remodeling, recurrent folate-responsive regulatory loci, and their response to folate and therapeutic perturbations.**
*Related to Figure 3.*

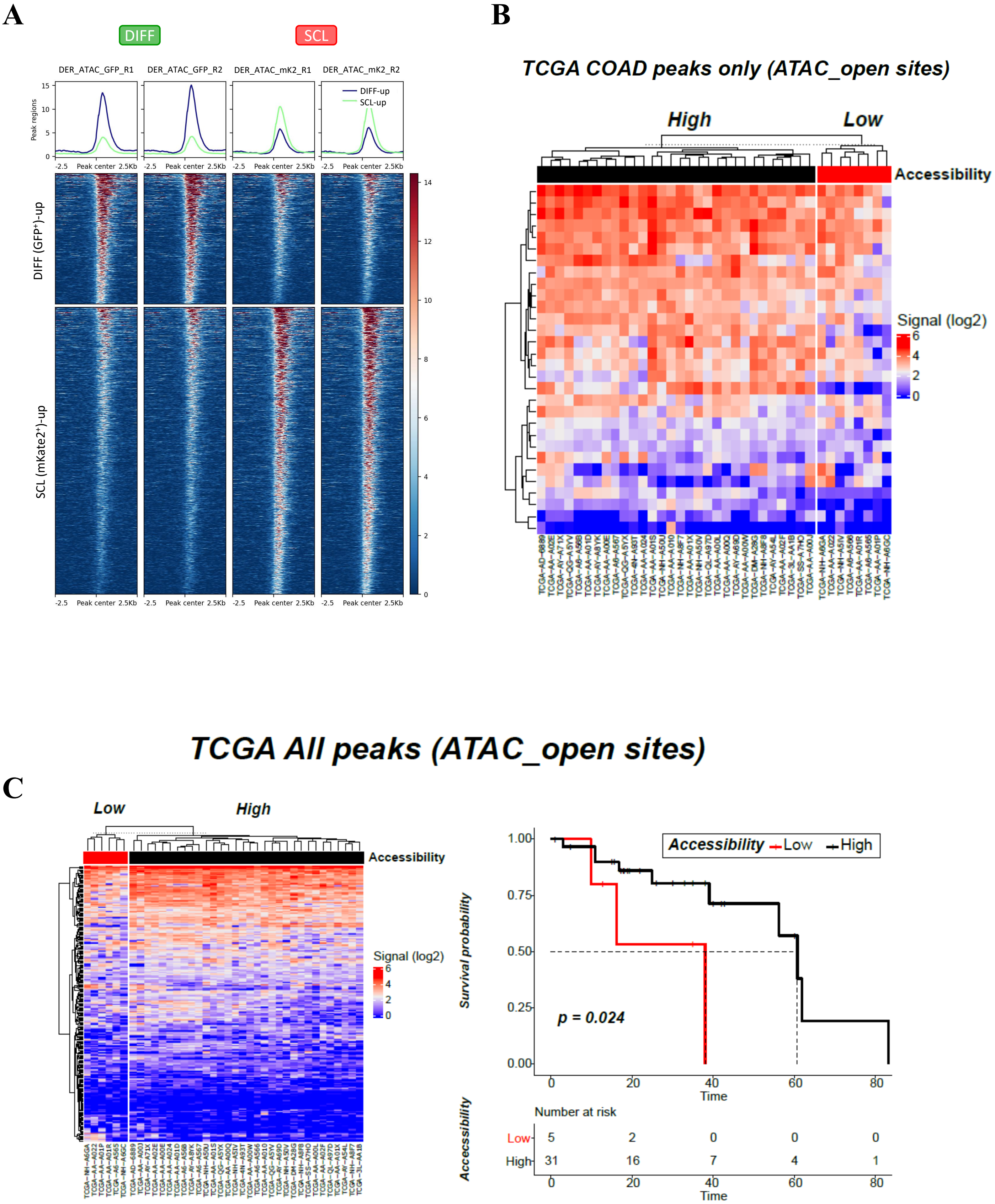
A
DIFF
SCL
DER_ATAC_GFP_R1
DER_ATAC_GFP_R2
DER_ATAC_mK2_R1
DER_ATAC_mK2_R2
Peak regions
DIFF-up
SCL-up
-2.5 Peak center 2.5Kb
DIFF (GFP+)-up
SCL (mKate2+)-up
B
TCGA COAD peaks only (ATAC_open sites)
High
Low
Accessibility
Signal (log2)
C
TCGA All peaks (ATAC_open sites)
Low
High
Accessibility
Signal (log2)
Accessibility — Low — High
Survival probability
p = 0.024
Time
Number at risk
Low 5 2 0 0 0
High 31 16 7 4 1
Accecssibility

**D**

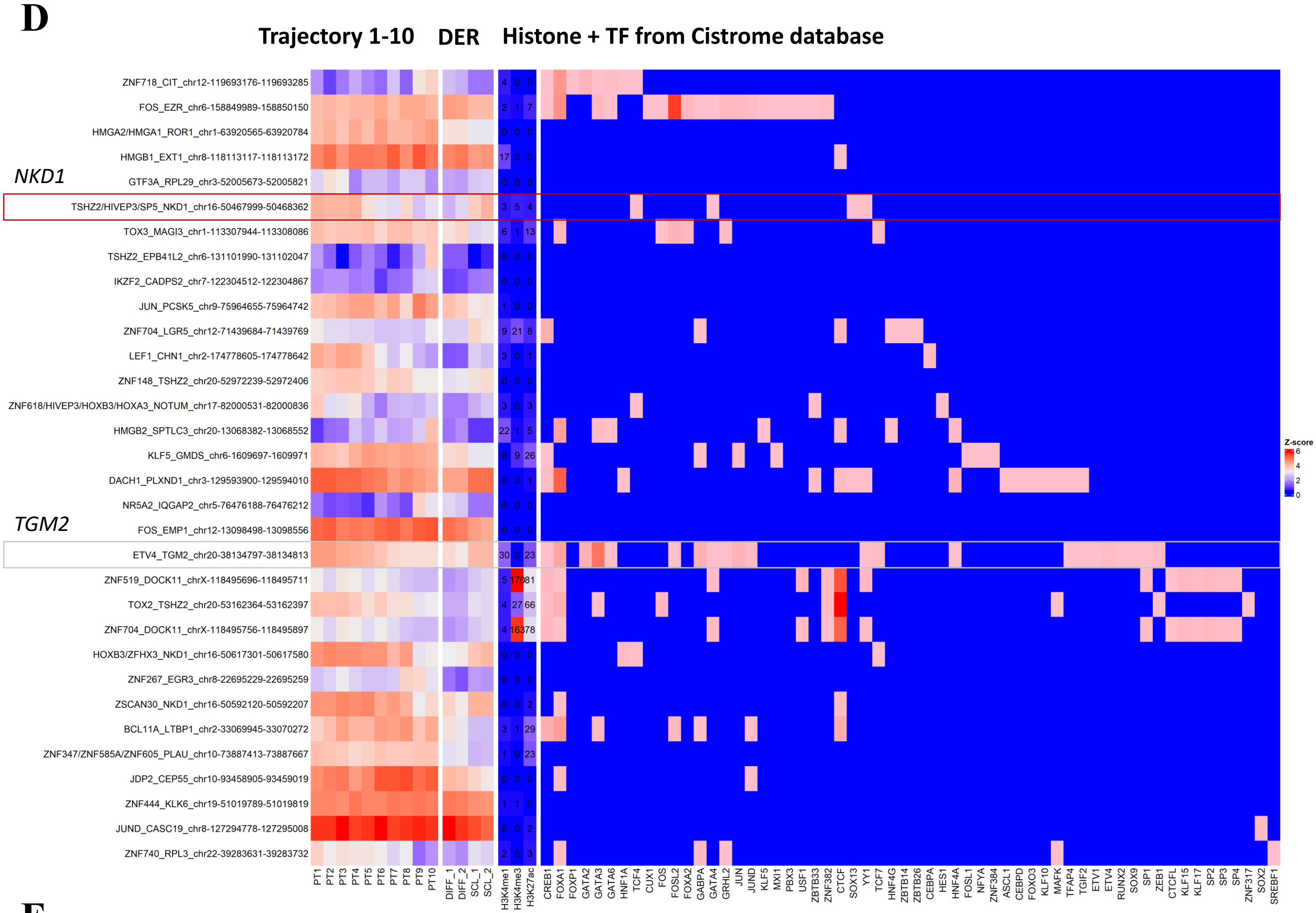

Trajectory 1-10
DER
Histone + TF from Cistrome database
ZNF718_CIT_chr12-119693176-119693285
FOS_EZR_chr6-158849989-158850150
HMGA2/HMGA1_ROR1_chr1-63920565-63920784
HMGB1_EXT1_chr8-118113117-118113172
NKD1
GTF3A_RPL29_chr3-52005673-52005821
TSHZ2/HIVEP3/SP5_NKD1_chr16-50467999-50468362
TOX3_MAGI3_chr1-113307944-113308086
TSHZ2_EPB41L2_chr6-131101990-131102047
IKZF2_CADPS2_chr7-122304512-122304867
JUN_PCSK5_chr9-75964655-75964742
ZNF704_LGR5_chr12-71439684-71439769
LEF1_CHN1_chr2-174778605-174778642
ZNF148_TSHZ2_chr20-52972239-52972406
ZNF618/HIVEP3/HOXB3/HOXA3_NOTUM_chr17-82000531-82000836
HMGB2_SPTLC3_chr20-13068382-13068552
KLF5_GMDS_chr6-1609697-1609971
DACH1_PLXND1_chr3-129593900-129594010
NR5A2_IQGAP2_chr5-76476188-76476212
TGM2
FOS_EMP1_chr12-13098498-13098556
ETV4_TGM2_chr20-38134797-38134813
ZNF519_DOCK11_chrX-118495696-118495711
TOX2_TSHZ2_chr20-53162364-53162397
ZNF704_DOCK11_chrX-118495756-118495897
HOXB3/ZFHX3_NKD1_chr16-50617301-50617580
ZNF267_EGR3_chr8-22695229-22695259
ZSCAN30_NKD1_chr16-50592120-50592207
BCL11A_LTBP1_chr2-33069945-33070272
ZNF347/ZNF585A/ZNF605_PLAU_chr10-73887413-73887667
JDP2_CEP55_chr10-93458905-93459019
ZNF444_KLK6_chr19-51019789-51019819
JUND_CASC19_chr8-127294778-127295008
ZNF740_RPL3_chr22-39283631-39283732
PT1 PT2 PT3 PT4 PT5 PT6 PT7 PT8 PT9 PT10
DIFF_1 DIFF_2 SCL_1 SCL_2
H3K4me1 H3K4me3 H3K27ac
CREB1 FOXA1 FOXP1 GATA2 GATA3 GATA6 HNF1A TCF4 CUX1 FOS FOSL2 FOXA2 GABPA GATA4 GRHL2 JUN JUND KLF5 MXI1 PBX3 USF1 ZBTB33 ZNF382 CTCF SOX13 YY1 TCF7 HNF4G ZBTB14 ZBTB26 CEBPA HES1 HNF4A FOSL1 NFYA ZNF384 ASCL1 CEBPD FOXO3 KLF10 MAFK TFAP4 TGIF2 ETV1 ETV4 RUNX2 SOX9 SP1 ZEB1 CTCFL KLF15 KLF17 SP2 SP3 SP4 ZNF317 SOX2 SREBF1
Z-score
6
4
2
0


**E**

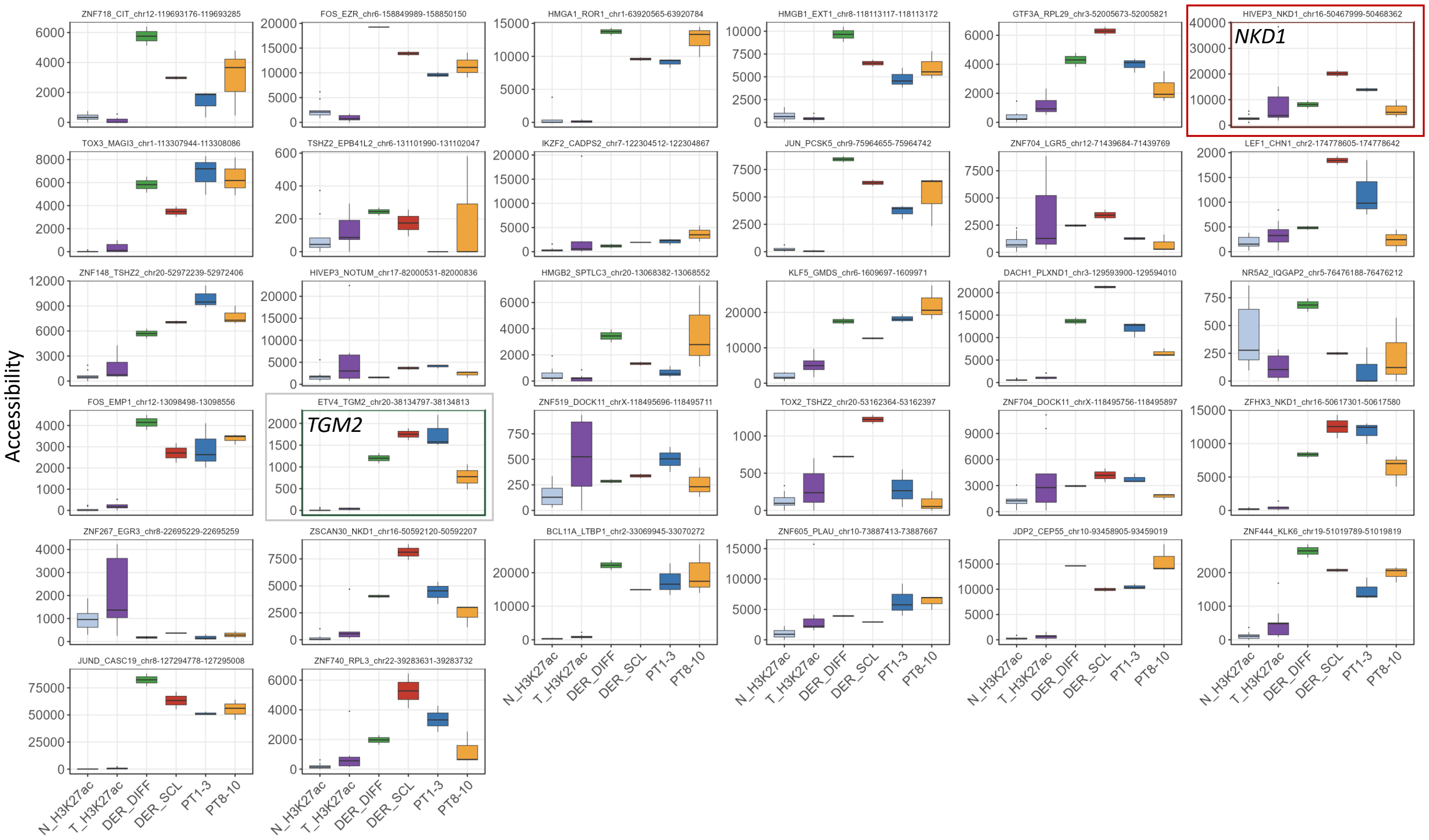

Accessibility
ZNF718_CIT_chr12-119693176-119693285
FOS_EZR_chr6-158849989-158850150
HMGA1_ROR1_chr1-63920565-63920784
HMGB1_EXT1_chr8-118113117-118113172
GTF3A_RPL29_chr3-52005673-52005821
HIVEP3_NKD1_chr16-50467999-50468362
NKD1
TOX3_MAGI3_chr1-113307944-113308086
TSHZ2_EPB41L2_chr6-131101990-131102047
IKZF2_CADPS2_chr7-122304512-122304867
JUN_PCSK5_chr9-75964655-75964742
ZNF704_LGR5_chr12-71439684-71439769
LEF1_CHN1_chr2-174778605-174778642
ZNF148_TSHZ2_chr20-52972239-52972406
HIVEP3_NOTUM_chr17-82000531-82000836
HMGB2_SPTLC3_chr20-13068382-13068552
KLF5_GMDS_chr6-1609697-1609971
DACH1_PLXND1_chr3-129593900-129594010
NR5A2_IQGAP2_chr5-76476188-76476212
FOS_EMP1_chr12-13098498-13098556
ETV4_TGM2_chr20-38134797-38134813
TGM2
ZNF519_DOCK11_chrX-118495696-118495711
TOX2_TSHZ2_chr20-53162364-53162397
ZNF704_DOCK11_chrX-118495756-118495897
ZFHX3_NKD1_chr16-50617301-50617580
ZNF267_EGR3_chr8-22695229-22695259
ZSCAN30_NKD1_chr16-50592120-50592207
BCL11A_LTBP1_chr2-33069945-33070272
ZNF605_PLAU_chr10-73887413-73887667
JDP2_CEP55_chr10-93458905-93459019
ZNF444_KLK6_chr19-51019789-51019819
JUND_CASC19_chr8-127294778-127295008
ZNF740_RPL3_chr22-39283631-39283732
N_H3K27ac T_H3K27ac DER_DIFF DER_SCL PT1-3 PT8-10

**F**

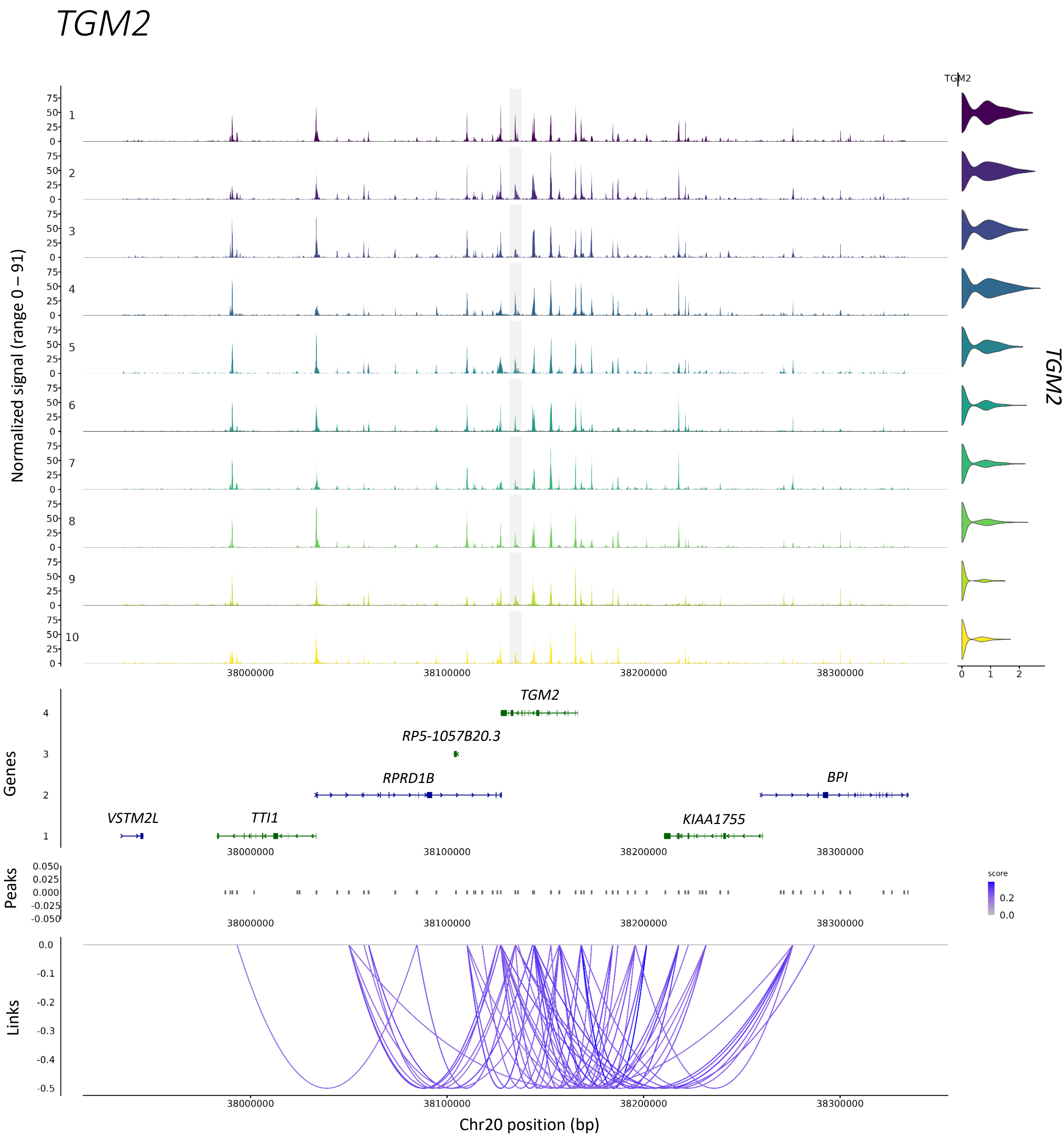

TGM2
Normalized signal (range 0 – 91)
1
2
3
4
5
6
7
8
9
10
0
25
50
75
TGM2
0
1
2
TGM2
38000000
38100000
38200000
38300000
Genes
TGM2
RP5-1057B20.3
RPRD1B
BPI
VSTM2L
TTI1
KIAA1755
Peaks
0.050
0.025
0.000
-0.025
-0.050
score
0.2
0.0
Links
0.0
-0.1
-0.2
-0.3
-0.4
-0.5
Chr20 position (bp)

**G**

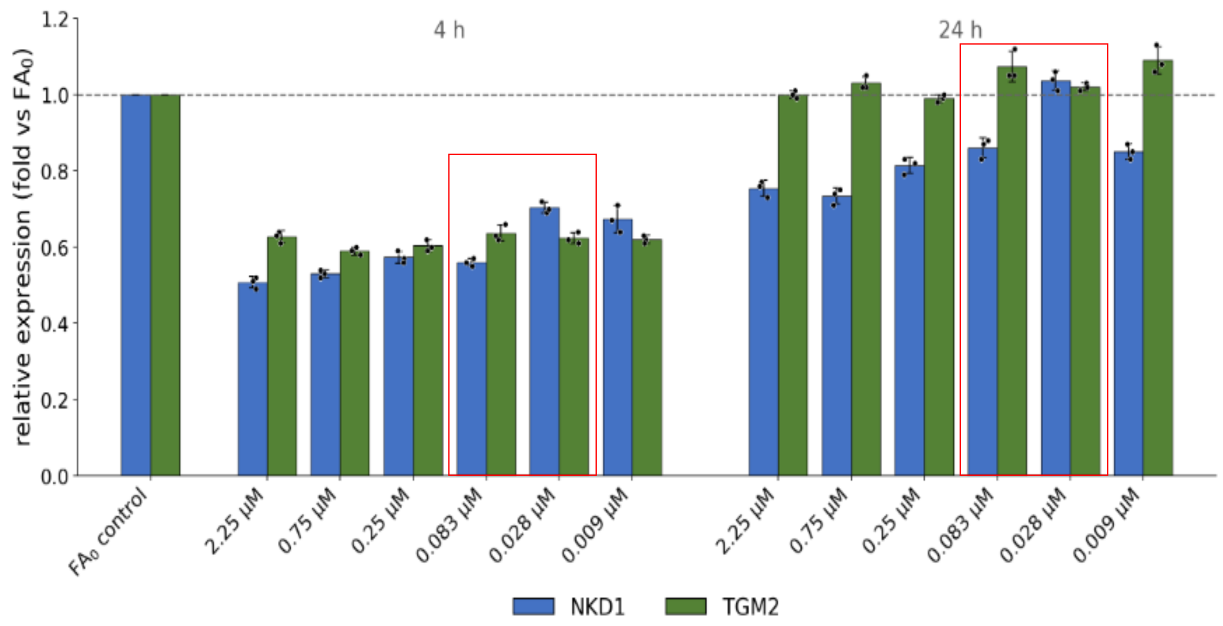
relative expression (fold vs FA0)
4 h
24 h
1.2
1.0
0.8
0.6
0.4
0.2
0.0
FA0 control
2.25 µM
0.75 µM
0.25 µM
0.083 µM
0.028 µM
0.009 µM
2.25 µM
0.75 µM
0.25 µM
0.083 µM
0.028 µM
0.009 µM
NKD1
TGM2

**H**

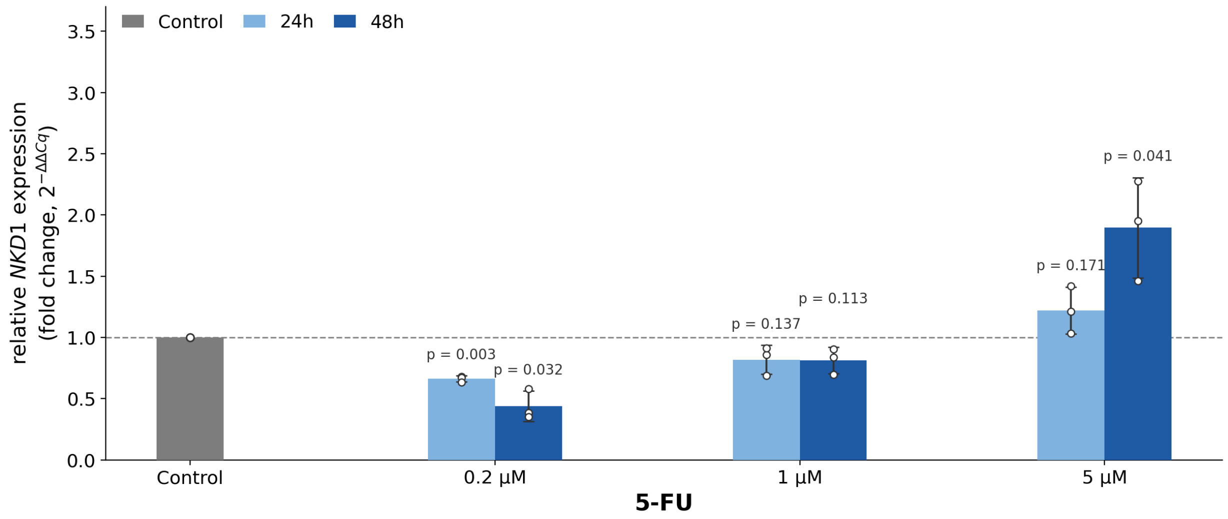
Control
24h
48h
relative NKD1 expression
(fold change, 2−ΔΔCq)
3.5
3.0
2.5
2.0
1.5
1.0
0.5
0.0
p = 0.003
p = 0.032
p = 0.137
p = 0.113
p = 0.171
p = 0.041
Control
0.2 µM
1 µM
5 µM
5-FU

**I**

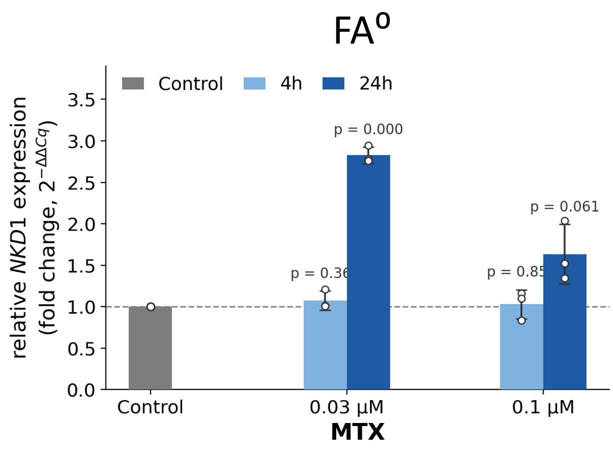
FA0
Control
4h
24h
relative NKD1 expression
(fold change, 2−ΔΔCq)
3.5
3.0
2.5
2.0
1.5
1.0
0.5
0.0
p = 0.000
p = 0.061
Control
0.03 µM
0.1 µM
MTX

**J**

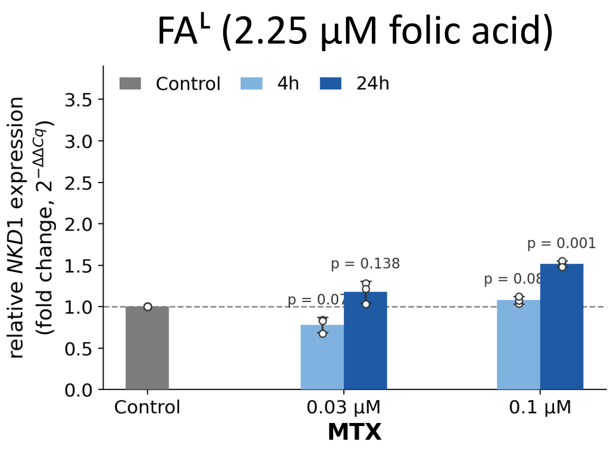
FAL (2.25 µM folic acid)
Control
4h
24h
relative NKD1 expression
(fold change, 2−ΔΔCq)
3.5
3.0
2.5
2.0
1.5
1.0
0.5
0.0
p = 0.138
p = 0.001
Control
0.03 µM
0.1 µM
MTX

**A,** Aggregate chromatin-accessibility profiles and heatmaps from bulk ATAC-seq of DER-sorted $GFP^+$ differentiated (DIFF) and $mKate2^+$ stem-like (SCL) populations, shown for two independent replicates per state across DIFF-up and SCL-up open chromatin regions.

**B,** Heatmap of COAD-specific trajectory-opening regulatory regions used for the Kaplan–Meier analysis in **Fig. 3I**. Samples were partitioned by k-means clustering into high- and low-accessibility groups comprising 30 and 6 patients, respectively.

**C,** Heatmap and Kaplan–Meier overall survival analysis using all trajectory-opening regulatory regions without COAD-specific filtering. Samples were partitioned using the same k-means approach into high- and low-accessibility groups comprising 31 and 5 patients, respectively. Statistical significance was determined by log-rank test.

**D,** Integrated heatmap summarizing folate-responsive gene regulatory network (GRN) loci across trajectory pseudotime, DER differential ATAC peaks, histone-associated chromatin marks, transcription factor ChIP-seq datasets and public Cistrome annotations. Columns summarize accessibility dynamics, DER state specificity and external regulatory evidence. *NKD1* and *TGM2*, prioritized by the ranking in **Fig. 3K**, are indicated.

**E,** Normalized RPKM signal comparison across GRN loci integrating colorectal tissue H3K27ac ChIP-seq, DER ATAC-seq and trajectory-resolved multiome accessibility states. Signal distributions are shown, in order, for normal colon H3K27ac (N_H3K27ac; light blue), colorectal cancer H3K27ac (T_H3K27ac; purple), DIFF-DER (ATAC) (green), SCL-DER (ATAC) (red), early pseudotime states (PT1–PT3, trajectory start; dark blue) and late pseudotime states (PT8–PT10, trajectory end; orange). *NKD1* and *TGM2*, prioritized by the ranking in **Fig. 3K**, are indicated.

**F,** Multiome pseudobulk visualization of the *TGM2* locus across pseudotime, including chromatin accessibility, gene expression and inferred peak-to-gene co-accessibility links.

**G,** *NKD1* (blue) and *TGM2* (green) expression across a graded folate concentration series (2.25, 0.75, 0.25, 0.083, 0.028 and 0.009 μM) at 4 h and 24 h, relative to the $FA^0$ control. Red boxes indicate concentrations within the physiological range of human plasma folate.

**H,** Relative *NKD1* expression in HT29 cells following 5-FU treatment (0.2, 1 and 5 μM) at 24 h and 48 h, relative to the untreated control. Statistical significance was determined by two-sided Student's *t*-test against the untreated control; exact *p* values are indicated.

**I,** Relative *NKD1* expression in HT29 cells cultured under $FA^0$ following methotrexate (MTX) treatment (0.03 and 0.1 μM) at 4 h and 24 h, relative to the untreated control. Statistical significance was determined by two-sided Student's *t*-test against the untreated control; exact *p* values are indicated.

**J,** Relative *NKD1* expression in HT29 cells cultured under $FA^L$ following methotrexate (MTX) treatment (0.03 and 0.1 μM) at 4 h and 24 h, relative to the untreated control. Statistical significance was determined by two-sided Student's *t*-test against the untreated control; exact *p* values are indicated.

Expression in panels G–J was measured by qRT–PCR, normalized to *ACTB* and expressed as $2^{-\Delta\Delta Cq}$ fold change relative to the indicated control. Data are presented as mean ± SD (*n* = 3 biological replicates), with dots representing individual values.

### Supplementary Figure S4.

**Folate-associated transcription factor rewiring and WNT attenuation.**
*Related to Figure 4.*

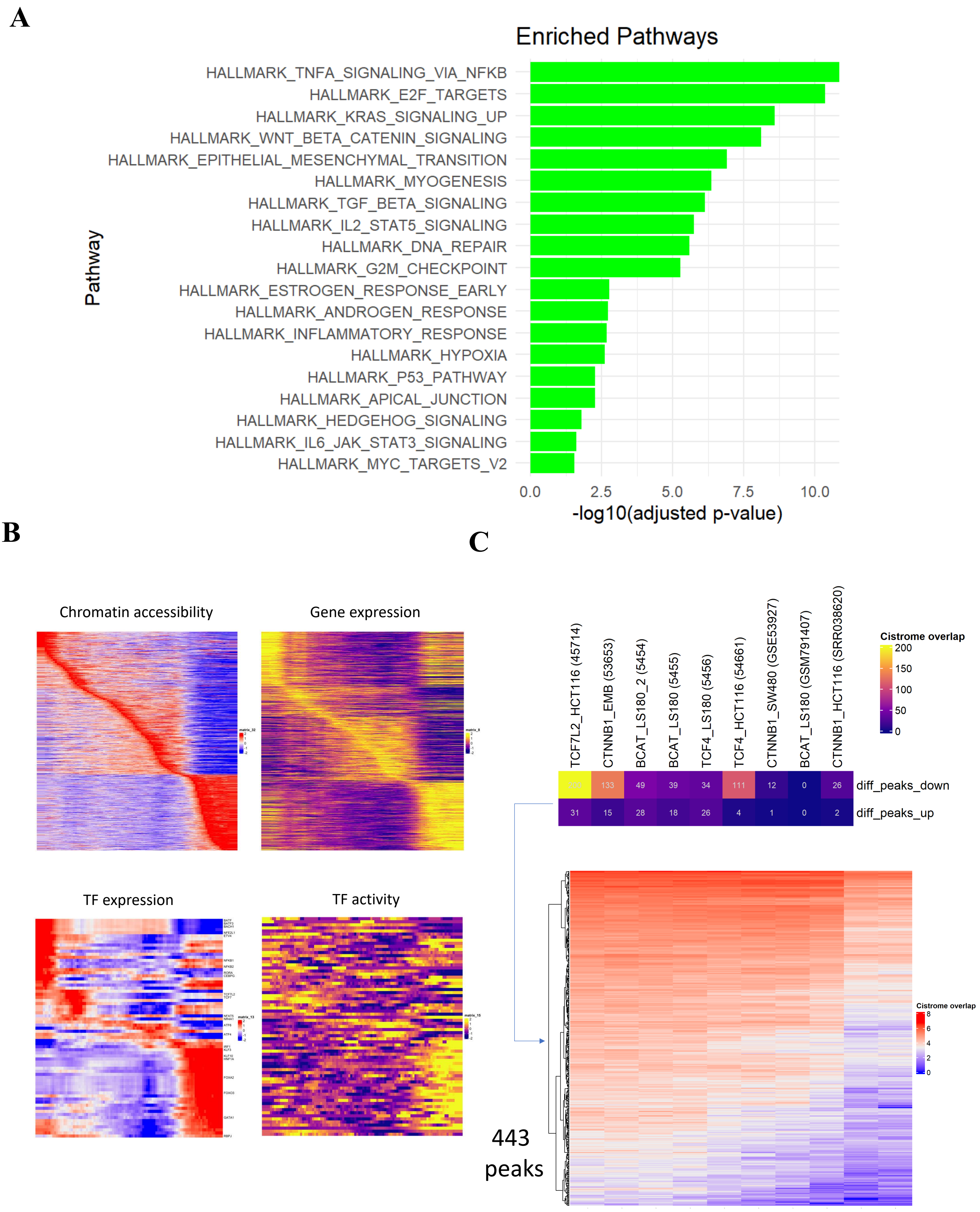
A
Enriched Pathways
Pathway
HALLMARK_TNFA_SIGNALING_VIA_NFKB
HALLMARK_E2F_TARGETS
HALLMARK_KRAS_SIGNALING_UP
HALLMARK_WNT_BETA_CATENIN_SIGNALING
HALLMARK_EPITHELIAL_MESENCHYMAL_TRANSITION
HALLMARK_MYOGENESIS
HALLMARK_TGF_BETA_SIGNALING
HALLMARK_IL2_STAT5_SIGNALING
HALLMARK_DNA_REPAIR
HALLMARK_G2M_CHECKPOINT
HALLMARK_ESTROGEN_RESPONSE_EARLY
HALLMARK_ANDROGEN_RESPONSE
HALLMARK_INFLAMMATORY_RESPONSE
HALLMARK_HYPOXIA
HALLMARK_P53_PATHWAY
HALLMARK_APICAL_JUNCTION
HALLMARK_HEDGEHOG_SIGNALING
HALLMARK_IL6_JAK_STAT3_SIGNALING
HALLMARK_MYC_TARGETS_V2
0.0
2.5
5.0
7.5
10.0
-log10(adjusted p-value)
B
Chromatin accessibility
Gene expression
TF expression
TF activity
C
TCF7L2_HCT116 (45714)
CTNNB1_EMB (53653)
BCAT_LS180_2 (5454)
BCAT_LS180 (5455)
TCF4_LS180 (5456)
TCF4_HCT116 (54661)
CTNNB1_SW480 (GSE53927)
BCAT_LS180 (GSM791407)
CTNNB1_HCT116 (SRR038620)
Cistrome overlap
200
150
100
50
0
200 133 49 39 34 111 12 0 26 diff_peaks_down
31 15 28 18 26 4 1 0 2 diff_peaks_up
Cistrome overlap
8
6
4
2
0
443 peaks

**D**

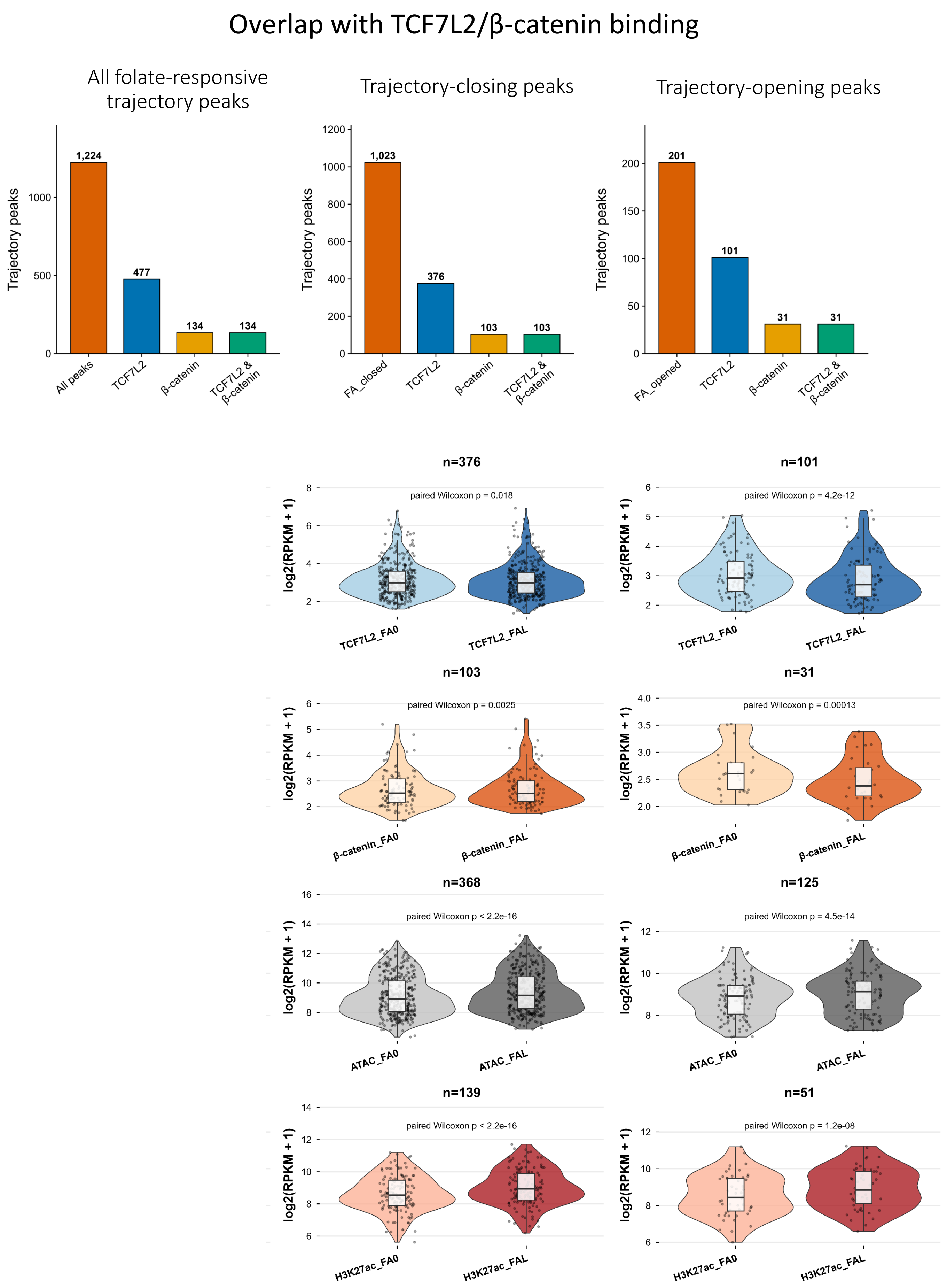

Overlap with TCF7L2/β-catenin binding
All folate-responsive trajectory peaks
Trajectory-closing peaks
Trajectory-opening peaks
Trajectory peaks
1,224
477
134
134
All peaks
TCF7L2
β-catenin
TCF7L2 & β-catenin
1,023
376
103
103
FA_closed
201
101
31
31
FA_opened
n=376
paired Wilcoxon p = 0.018
n=101
paired Wilcoxon p = 4.2e-12
log2(RPKM + 1)
TCF7L2_FA0
TCF7L2_FAL
n=103
paired Wilcoxon p = 0.0025
n=31
paired Wilcoxon p = 0.00013
β-catenin_FA0
β-catenin_FAL
n=368
paired Wilcoxon p < 2.2e-16
n=125
paired Wilcoxon p = 4.5e-14
ATAC_FA0
ATAC_FAL
n=139
paired Wilcoxon p < 2.2e-16
n=51
paired Wilcoxon p = 1.2e-08
H3K27ac_FA0
H3K27ac_FAL

**E**

**β-catenin localization (IHC)**

**$FA^0$** **$FA^L$**

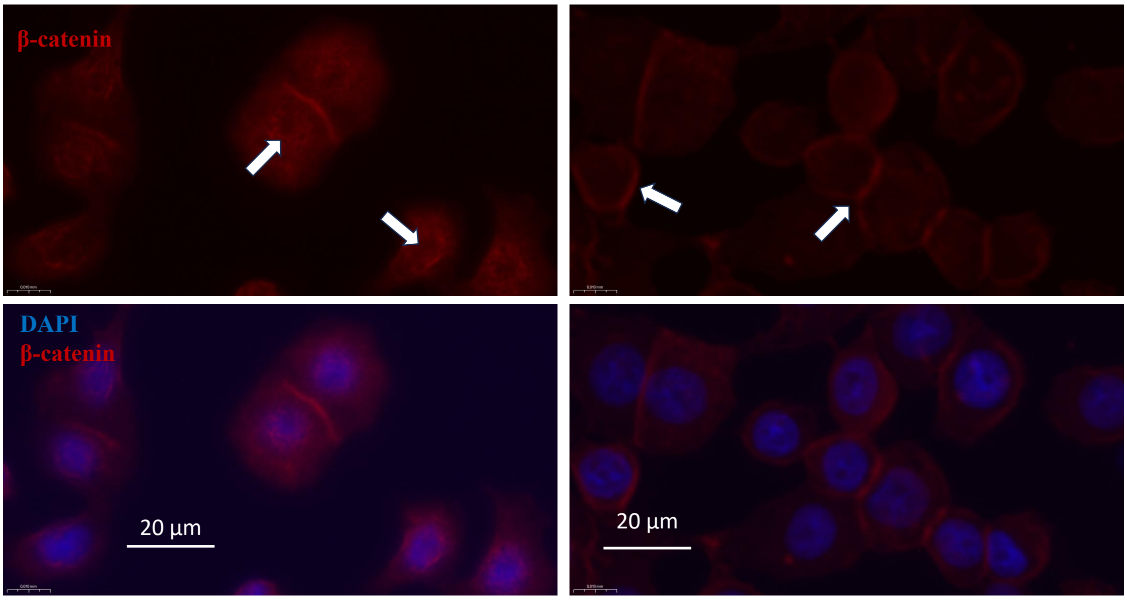


**F**

β-catenin localization (PLA)

$FA^0$ $FA^L$

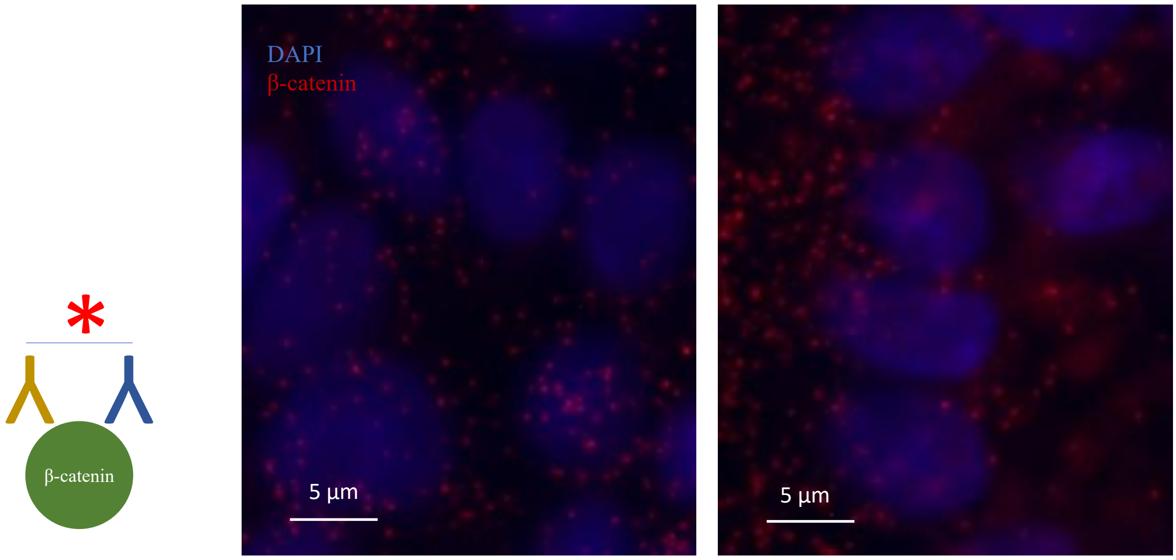


**G**

β-catenin-TCF7L2 co-localization (PLA)

$FA^0$ $FA^L$

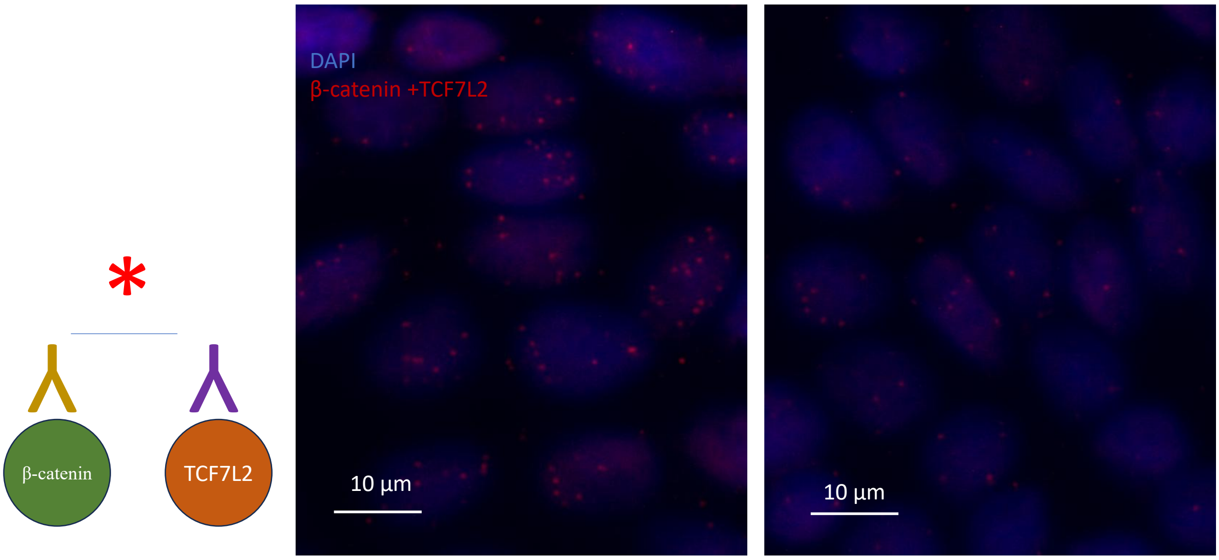

**A,** Hallmark pathway enrichment analysis of transcription factor-associated regulatory programs identified by chromVAR. Bars indicate significantly enriched MSigDB Hallmark gene sets ranked by $-\log_{10}$(adjusted *p* value).

**B,** Integrated pseudotime heatmaps of chromatin accessibility, gene expression, transcription factor expression, and inferred transcription factor activity across trajectory-ordered epithelial cells.

**C,** Integrated Cistrome overlap analysis of trajectory-associated differential chromatin accessibility regions with public TCF7L2 and β-catenin ChIP-seq datasets across colorectal cancer cell lines. Upper panel summarizes overlap counts across datasets; lower panel shows the recurrent subset of 443 overlapping chromatin accessibility regions.

**D,** Quantification of overlap between folate-responsive trajectory-associated chromatin accessibility regions and TCF7L2/β-catenin occupancy. Upper panels summarize overlap counts for all folate-responsive, trajectory-closing and trajectory-opening regions. Lower panels compare TCF7L2 ChIP-seq, β-catenin ChIP-seq, chromatin accessibility (ATAC-seq) and H3K27ac signal across $FA^L$ and $FA^0$ conditions within trajectory-closing and trajectory-opening regions. Statistical significance was determined using a paired Wilcoxon signed-rank test; exact *p* values are indicated.

**E,** Representative immunofluorescence images showing β-catenin localization in HT29 cells cultured under $FA^L$ and $FA^0$ conditions. White arrows indicate membrane-associated β-catenin. Nuclei were counterstained with DAPI. Scale bars, 20 µm. Images are representative of two independent experiments.

**F,** Representative proximity ligation assay (PLA) images detecting β-catenin in HT29 cells cultured under $FA^L$ and $FA^0$ conditions. Nuclei were counterstained with DAPI. Scale bars, 5 µm. Images are representative of two independent experiments.

**G,** Representative proximity ligation assay (PLA) images detecting β-catenin–TCF7L2 interactions in HT29 cells cultured under $FA^L$ and $FA^0$ conditions. Nuclei were counterstained with DAPI. Scale bars, 10 µm. Images are representative of two independent experiments.

### Supplementary Figure S5.

**Targeted metabolomics and chromatin alterations associated with folate perturbation.**
*Related to Figure 5.*

**A**

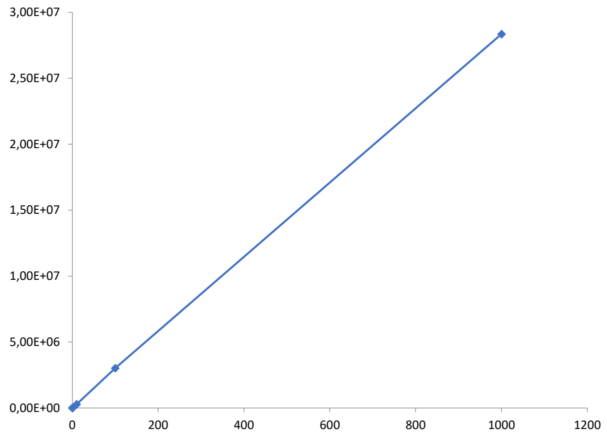


**B**

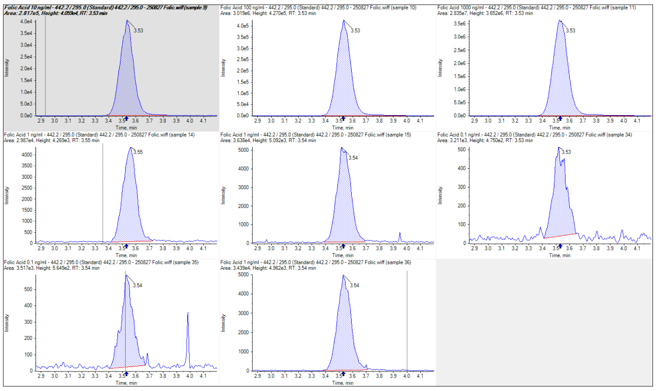

**C**

| Sample Name | Sample Type | Q1/Q3 | Actual Concentration | Area | Retention Time | Signal / Noise | Calculated Concentration | Accuracy |
|---|---|---|---|---|---|---|---|---|
| Folic Acid 0.1 ng/ml | Standard | 442.2 / 295.0 | 0.1 | 3.51E+03 | 3.53 | 36.3 | 0.08 | 82.77 |
| Folic Acid 0.1 ng/ml | Standard | 442.2 / 295.0 | 0.1 | 3.58E+03 | 3.54 | 46.30 | 0.09 | 85.15 |
| Folic Acid 1 ng/ml | Standard | 442.2 / 295.0 | 1 | 3.44E+04 | 3.54 | 433 | 1.17 | 116.53 |
| Folic Acid 1 ng/ml | Standard | 442.2 / 295.0 | 1 | 2.99E+04 | 3.55 | 259.4 | 1.01 | 100.7 |
| Folic Acid 1 ng/ml | Standard | 442.2 / 295.0 | 1 | 3.29E+04 | 3.54 | 289.7 | 1.11 | 111.28 |
| Folic Acid 10 ng/ml | Standard | 442.2 / 295.0 | 10 | 2.82E+05 | 3.53 | 2052.9 | 9.84 | 98.37 |
| Folic Acid 100 ng/ml | Standard | 442.2 / 295.0 | 100 | 3.02E+06 | 3.53 | 6880.3 | 105.79 | 105.79 |
| Folic Acid 1000 ng/ml | Standard | 442.2 / 295.0 | 1000 | 2.84E+07 | 3.53 | 13877.8 | 994.12 | 99.41 |

**D**

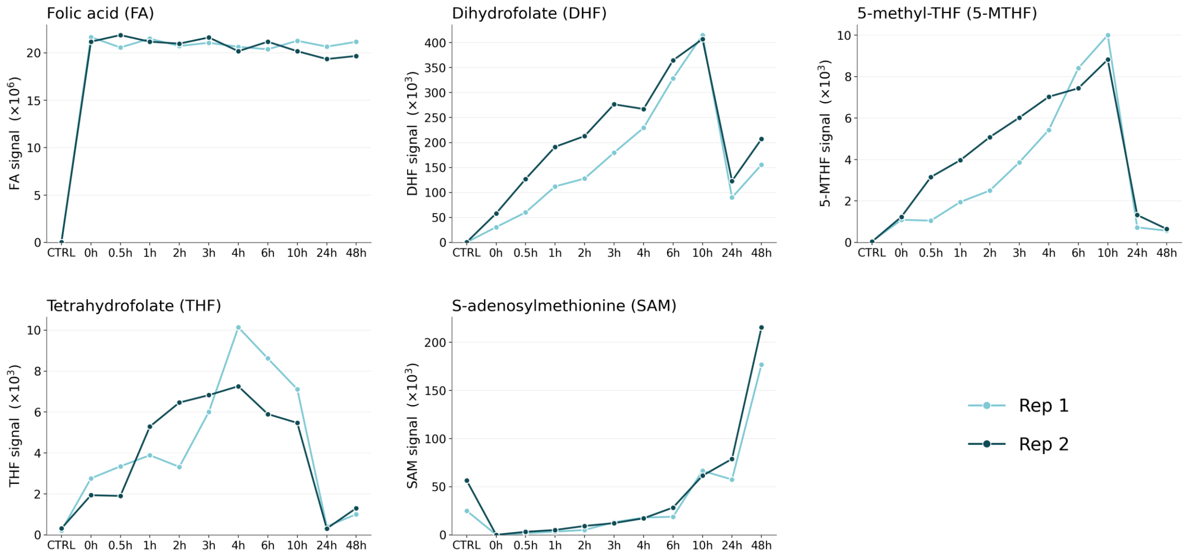


**E**

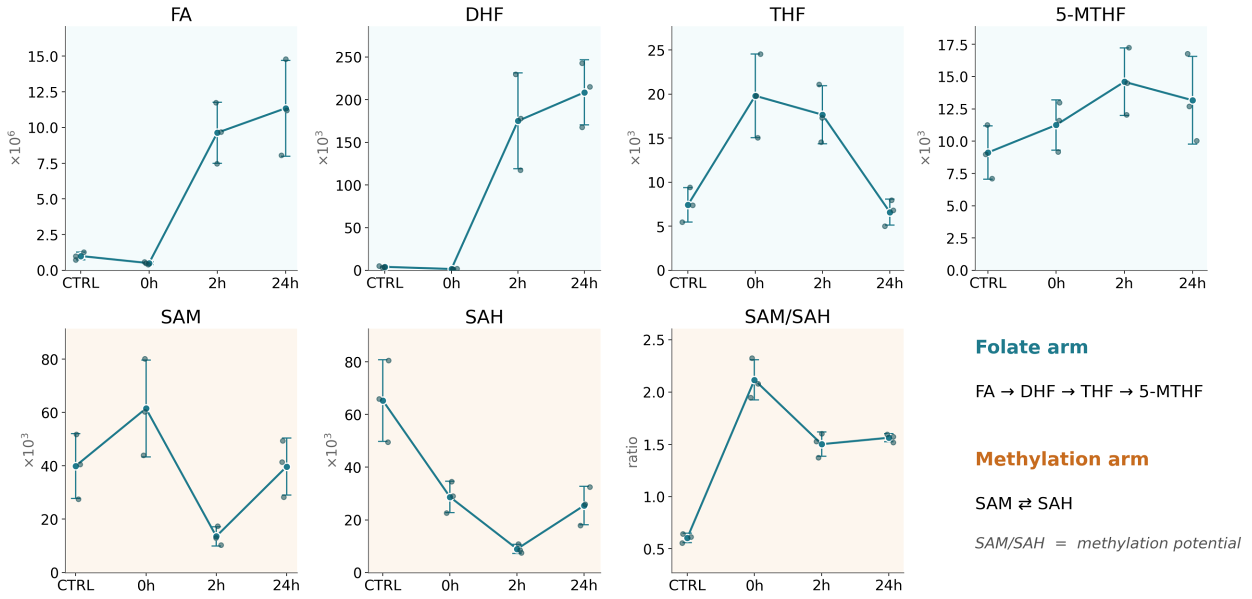

**F**

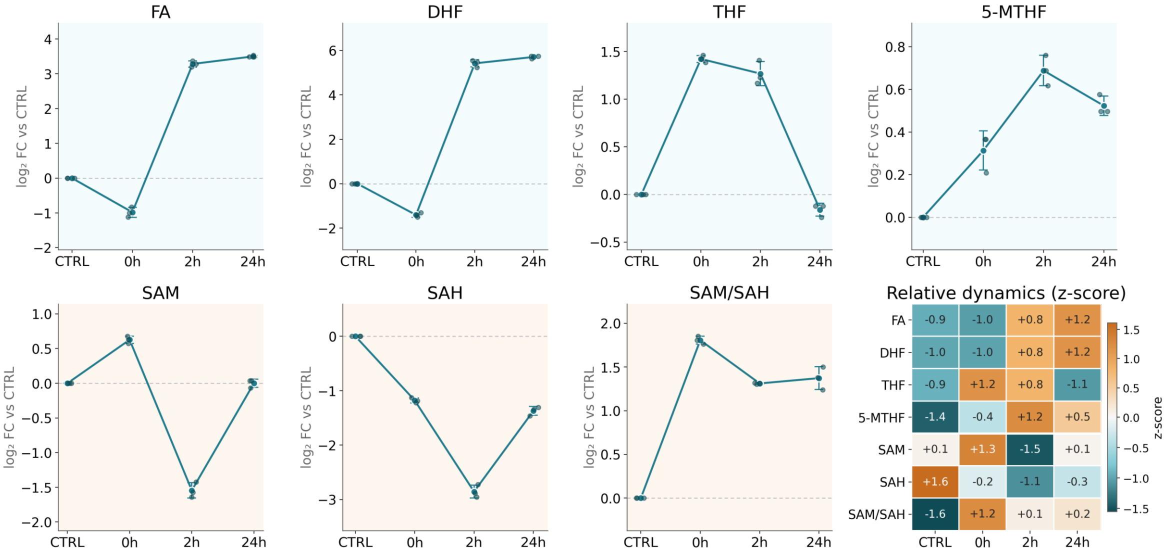

FA
DHF
THF
5-MTHF
SAM
SAH
SAM/SAH
log₂ FC vs CTRL
CTRL
0h
2h
24h
Relative dynamics (z-score)
FA -0.9 -1.0 +0.8 +1.2
DHF -1.0 -1.0 +0.8 +1.2
THF -0.9 +1.2 +0.8 -1.1
5-MTHF -1.4 -0.4 +1.2 +0.5
SAM +0.1 +1.3 -1.5 +0.1
SAH +1.6 -0.2 -1.1 -0.3
SAM/SAH -1.6 +1.2 +0.1 +0.2
z-score


**G**

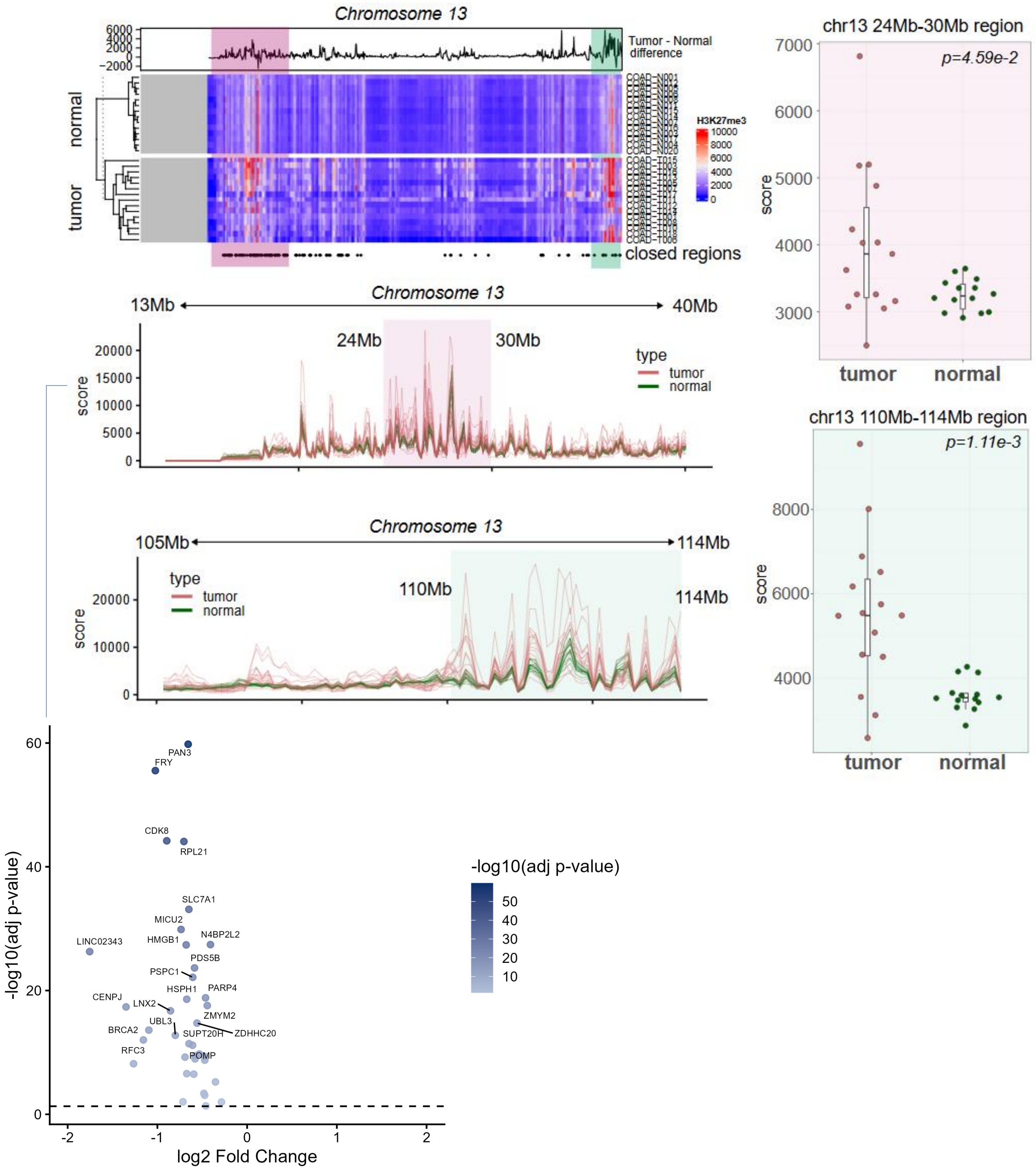

Chromosome 13
Tumor - Normal difference
normal
tumor
H3K27me3
closed regions
13Mb
40Mb
24Mb
30Mb
105Mb
114Mb
110Mb
type
tumor
normal
score
chr13 24Mb-30Mb region
p=4.59e-2
chr13 110Mb-114Mb region
p=1.11e-3
PAN3
FRY
CDK8
RPL21
SLC7A1
MICU2
N4BP2L2
LINC02343
HMGB1
PDS5B
PSPC1
HSPH1
PARP4
CENPJ
LNX2
ZMYM2
UBL3
ZDHHC20
BRCA2
SUPT20H
RFC3
POMP
-log10(adj p-value)
log2 Fold Change

**H**

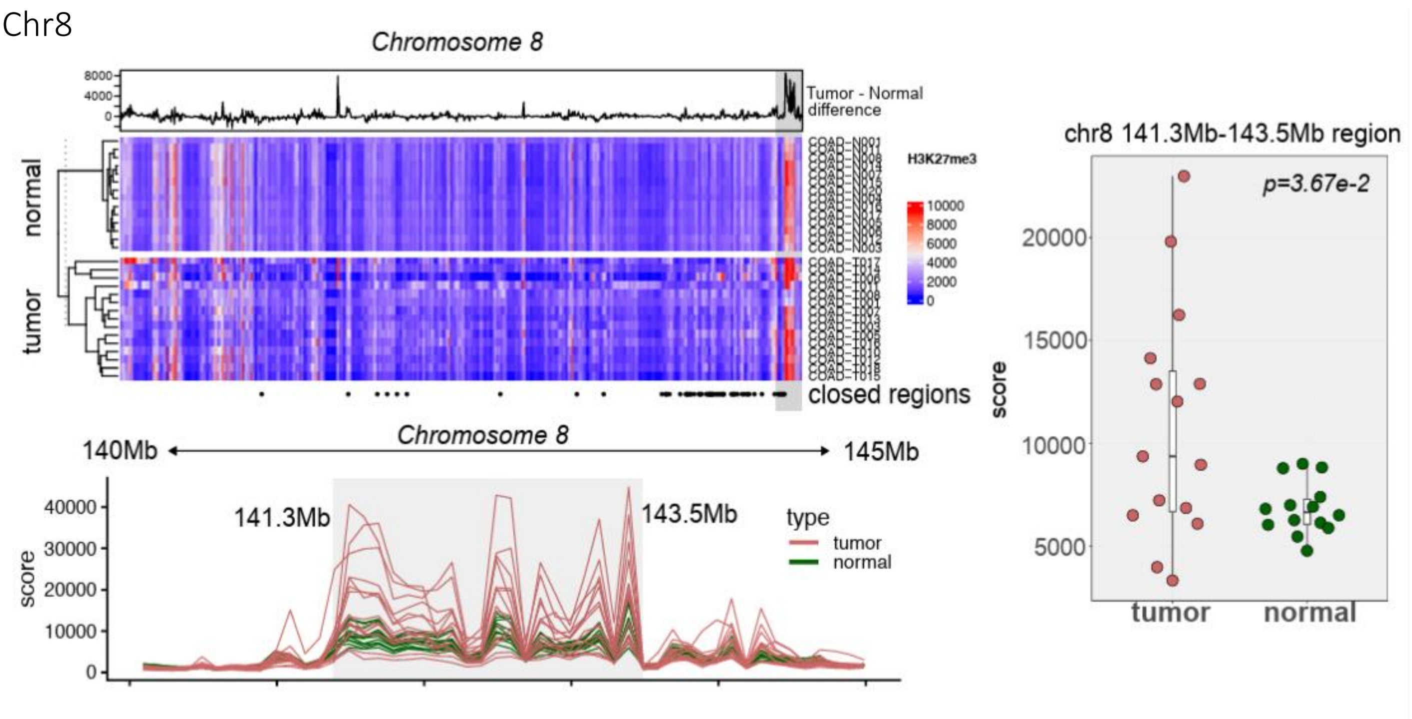

Chr8
Chromosome 8
Tumor - Normal difference
H3K27me3
normal
tumor
closed regions
140Mb
145Mb
141.3Mb
143.5Mb
type
tumor
normal
score
chr8 141.3Mb-143.5Mb region
p=3.67e-2


**I**

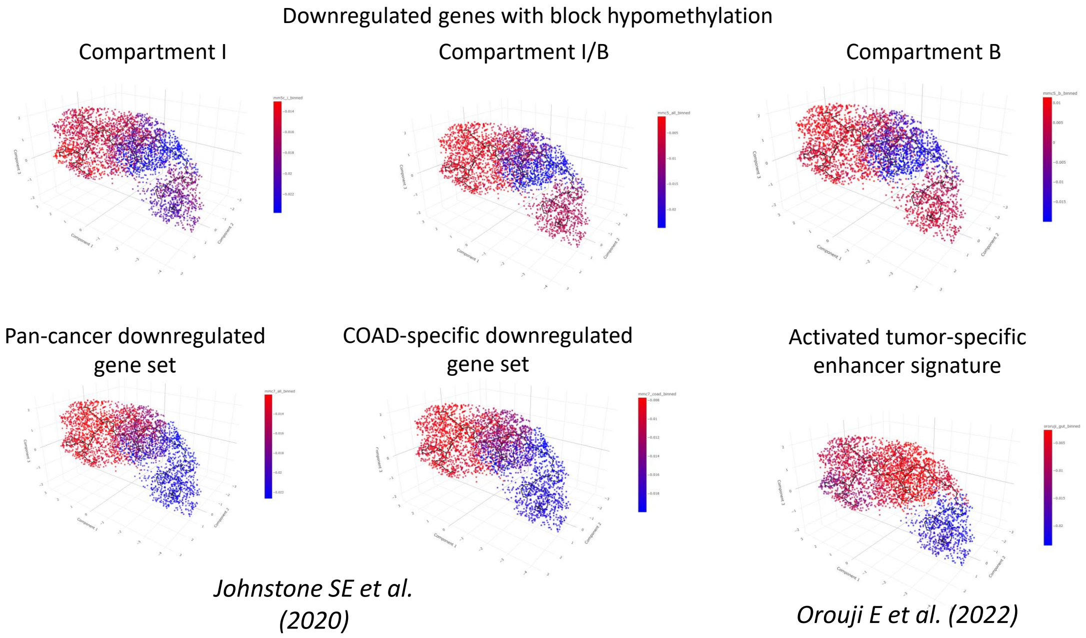

Downregulated genes with block hypomethylation
Compartment I
Compartment I/B
Compartment B
Pan-cancer downregulated gene set
COAD-specific downregulated gene set
Activated tumor-specific enhancer signature
Johnstone SE et al. (2020)
Orouji E et al. (2022)

**A,** Folic acid calibration curve for targeted LC–MS/MS quantification, showing peak area as a function of standard concentration across the analyzed range.

**B,** Representative extracted ion chromatograms (Q1/Q3 442.2/295.0) of folic acid standards spanning the calibration range, demonstrating consistent retention time and peak detection.

**C,** Calibration table for folic acid standards (0.1–1000 ng/ml), including Q1/Q3 transition, peak area, retention time, signal-to-noise ratio, calculated concentration, and quantification accuracy.

**D,** Time-resolved targeted LC–MS/MS quantification of extracellular metabolites in culture supernatants following folate restoration across ten time points (CTRL, 0, 0.5, 1, 2, 3, 4, 6, 10, 24 and 48 h). Signal intensities of folic acid (FA), dihydrofolate (DHF), tetrahydrofolate (THF), 5-methyltetrahydrofolate (5-MTHF) and S-adenosylmethionine (SAM) are shown for two independent biological replicates ($n$ = 2).

**E,** Relative intracellular levels of folate-cycle metabolites (FA, DHF, THF, and 5-MTHF) and methylation-cycle metabolites (SAM, SAH, and the SAM/SAH ratio) measured by targeted LC–MS/MS in HT29 cell lysates following folate restoration after folate starvation. Samples were collected at baseline (CTRL) and at 0 h, 2 h, and 24 h following folate restoration. Values are presented as mean ± SD ($n$ = 3 biological replicates).

**F,** Relative intracellular dynamics of folate-cycle metabolites (FA, DHF, THF, and 5-MTHF) and methylation-cycle metabolites (SAM, SAH, and the SAM/SAH ratio), displayed as metabolite-wise z-scores. Line plots show the normalized response of individual metabolites, whereas the accompanying heatmap provides an integrated overview of relative metabolite dynamics across the indicated conditions.

**G,** Tumor-enriched H3K27me3 chromatin domains on chromosome 13 and differential expression of resident genes. H3K27me3 profiles were obtained from GSE136888 (43). H3K27me3 enrichment heatmap across chromosome 13 in normal and tumor tissue samples (hierarchically clustered), together with the tumor–normal difference track and annotated trajectory-associated closed regions. Regional H3K27me3 profiles are shown across the 13–40 Mb interval (24–30 Mb domain highlighted) and the 105–114 Mb interval (110–114 Mb domain highlighted; tumor, red; normal, green). Right, per-sample quantification of H3K27me3 signal within the chr13 24–30 Mb ($p = 4.59 \times 10^{-2}$) and chr13 110–114 Mb ($p = 1.11 \times 10^{-3}$) regions in 15 tumor and 14 normal samples. Bottom, volcano plot of differential gene expression for genes located within the chr13 24–30 Mb H3K27me3-enriched domain following folate restoration in HT29 single-cell multiome data ($\log_2$ fold change versus $-\log_{10}$ adjusted p value), with significantly downregulated genes labeled. Statistical significance was determined by Wilcoxon rank-sum test.

**H,** Tumor-enriched H3K27me3 chromatin domains on chromosome 8, based on reanalysis of the same profiles (43; GSE136888). H3K27me3 enrichment heatmap across chromosome 8 in normal and tumor tissue samples (hierarchically clustered), together with the corresponding tumor–normal difference track and annotated trajectory-associated closed regions. Lower panel, regional H3K27me3 profiles across the 140–145 Mb interval (141.3–143.5 Mb domain highlighted; tumor, red; normal, green). Right, per-sample quantification of H3K27me3 signal within the chr8 141.3–143.5 Mb region in 15 tumor and 14 normal samples ($p = 3.67 \times 10^{-2}$). Statistical significance was determined by Wilcoxon rank-sum test.

**I,** Projection of previously published (39, 44) colorectal cancer-associated chromatin-state and enhancer signatures onto the folate-responsive epithelial trajectory. Cells are colored according to aggregated signature score (red, high; blue, low). Shown are compartment I-, compartment I/B-, and compartment B-associated block hypomethylation gene sets together with pan-cancer downregulated, COAD-specific downregulated, and activated tumor-specific enhancer signature.

### Supplementary Figure S6.

**Quantitative characterization and tissue validation of polarized epithelial trajectory remodeling.**
*Related to Figure 6.*

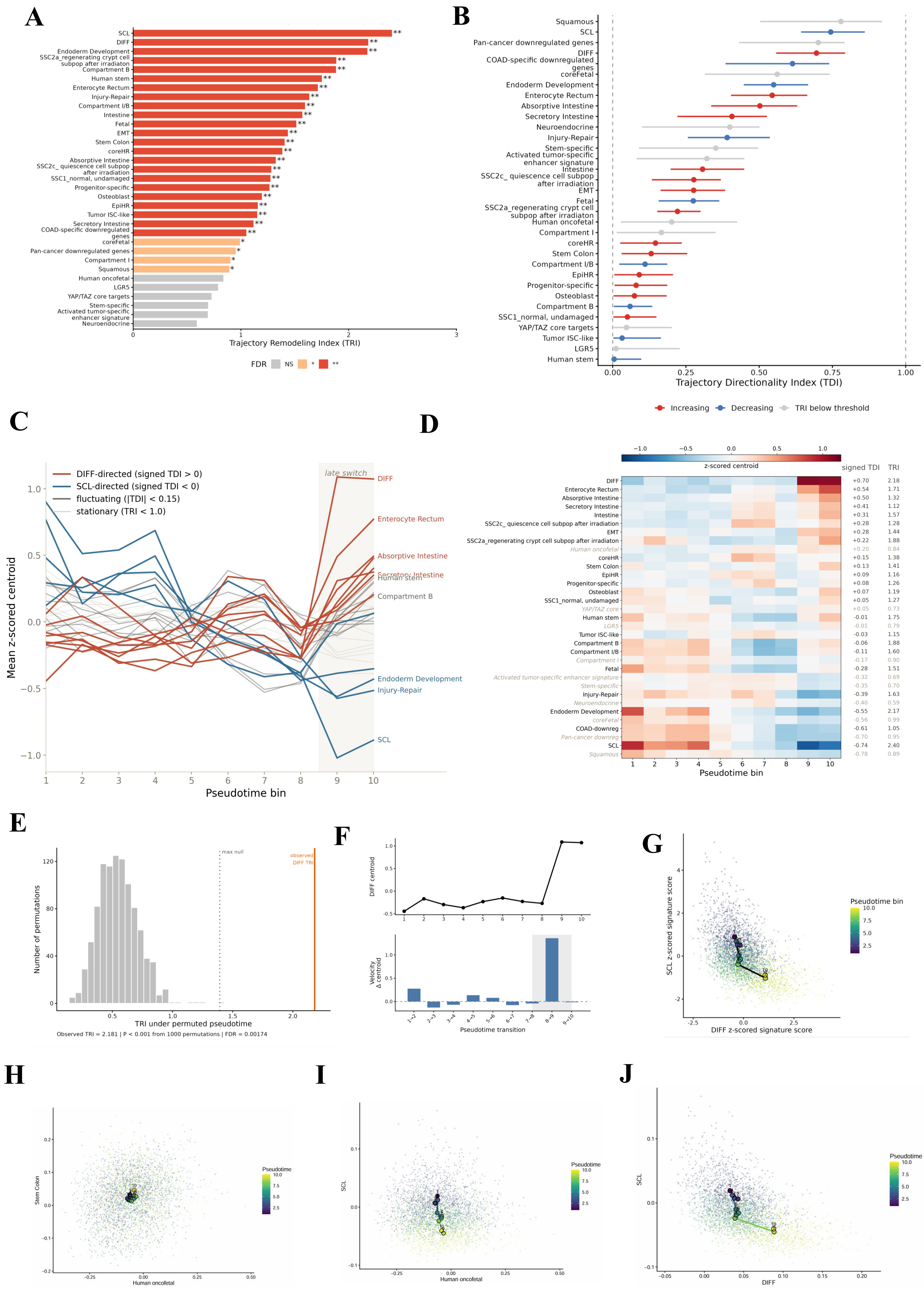
A
B
C
D
E
F
G
H
I
J
Trajectory Remodeling Index (TRI)
FDR NS * **
SCL
DIFF
Endoderm Development
SSC2a_regenerating crypt cell subpop after irradiation
Compartment B
Human stem
Enterocyte Rectum
Injury-Repair
Compartment I/B
Intestine
Fetal
EMT
Stem Colon
coreHR
Absorptive Intestine
SSC2c_ quiescence cell subpop after irradiation
SSC1_normal, undamaged
Progenitor-specific
Osteoblast
EpiHR
Tumor ISC-like
Secretory Intestine
COAD-specific downregulated genes
coreFetal
Pan-cancer downregulated genes
Compartment I
Squamous
Human oncofetal
LGR5
YAP/TAZ core targets
Stem-specific
Activated tumor-specific enhancer signature
Neuroendocrine
Trajectory Directionality Index (TDI)
Increasing
Decreasing
TRI below threshold
Mean z-scored centroid
Pseudotime bin
DIFF-directed (signed TDI > 0)
SCL-directed (signed TDI < 0)
fluctuating (|TDI| < 0.15)
stationary (TRI < 1.0)
late switch
z-scored centroid
signed TDI
TRI
Number of permutations
TRI under permuted pseudotime
max null
observed DIFF TRI
Observed TRI = 2.181 | P < 0.001 from 1000 permutations | FDR = 0.00174
DIFF centroid
Velocity Δ centroid
Pseudotime transition
SCL z-scored signature score
DIFF z-scored signature score
Pseudotime bin
Stem Colon
Human oncofetal
SCL
DIFF
Pseudotime

K

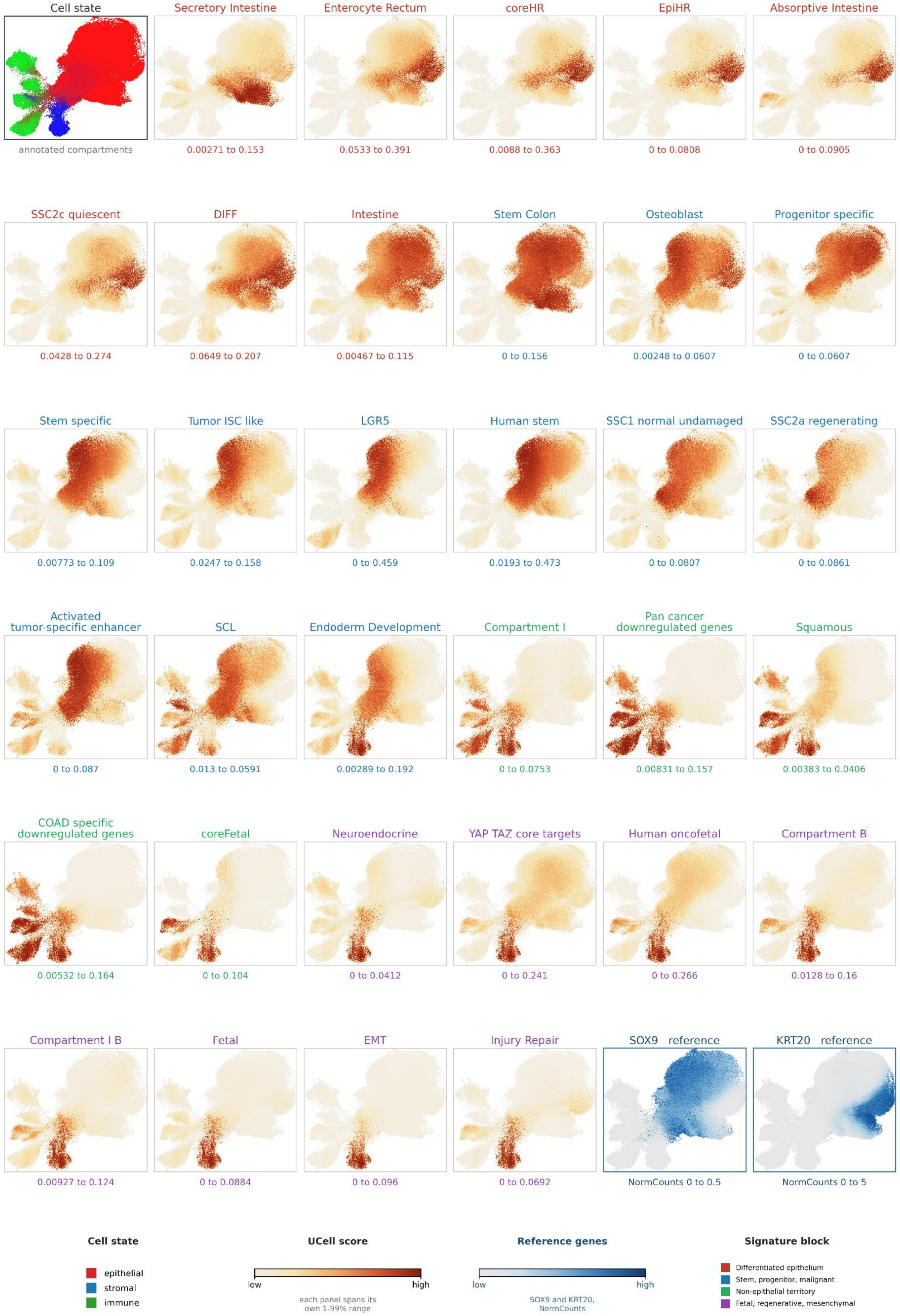
Cell state
annotated compartments
Secretory Intestine
0.00271 to 0.153
Enterocyte Rectum
0.0533 to 0.391
coreHR
0.0088 to 0.363
EpiHR
0 to 0.0808
Absorptive Intestine
0 to 0.0905
SSC2c quiescent
0.0428 to 0.274
DIFF
0.0649 to 0.207
Intestine
0.00467 to 0.115
Stem Colon
0 to 0.156
Osteoblast
0.00248 to 0.0607
Progenitor specific
0 to 0.0607
Stem specific
0.00773 to 0.109
Tumor ISC like
0.0247 to 0.158
LGR5
0 to 0.459
Human stem
0.0193 to 0.473
SSC1 normal undamaged
0 to 0.0807
SSC2a regenerating
0 to 0.0861
Activated tumor-specific enhancer
0 to 0.087
SCL
0.013 to 0.0591
Endoderm Development
0.00289 to 0.192
Compartment I
0 to 0.0753
Pan cancer downregulated genes
0.00831 to 0.157
Squamous
0.00383 to 0.0406
COAD specific downregulated genes
0.00532 to 0.164
coreFetal
0 to 0.104
Neuroendocrine
0 to 0.0412
YAP TAZ core targets
0 to 0.241
Human oncofetal
0 to 0.266
Compartment B
0.0128 to 0.16
Compartment I B
0.00927 to 0.124
Fetal
0 to 0.0884
EMT
0 to 0.096
Injury Repair
0 to 0.0692
SOX9 reference
NormCounts 0 to 0.5
KRT20 reference
NormCounts 0 to 5
Cell state
epithelial
stromal
immune
UCell score
low
high
each panel spans its own 1-99% range
Reference genes
low
high
SOX9 and KRT20, NormCounts
Signature block
Differentiated epithelium
Stem, progenitor, malignant
Non-epithelial territory
Fetal, regenerative, mesenchymal

**Epithelial-filtered UCell activity of 33 programmes across Normal–Adenoma–CRC**

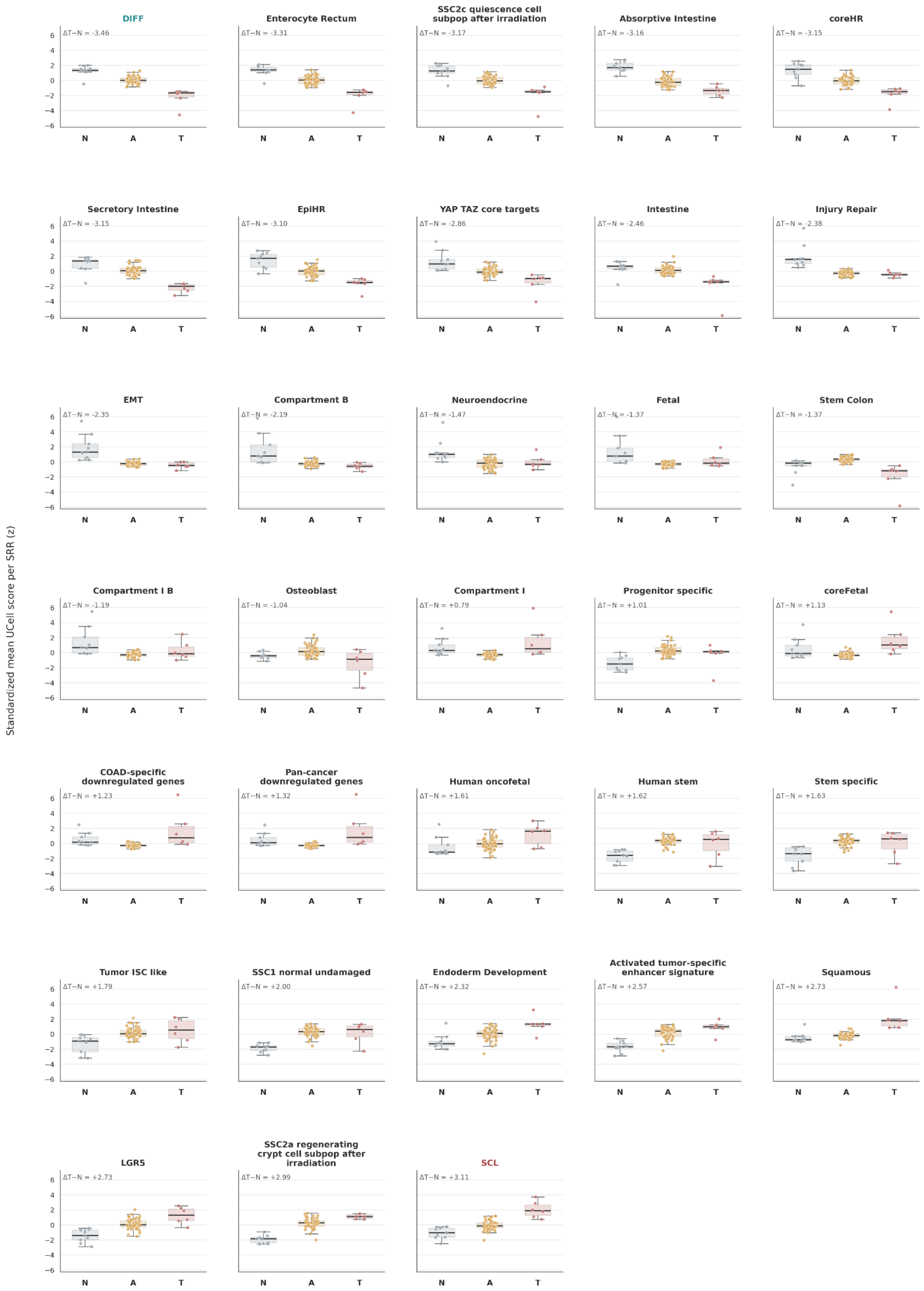


Epithelium and malignant epithelium; unaffected tissue excluded. Scores were standardized within programme across SRR means; all panels share the same y-axis. No inferential P values are shown.

M

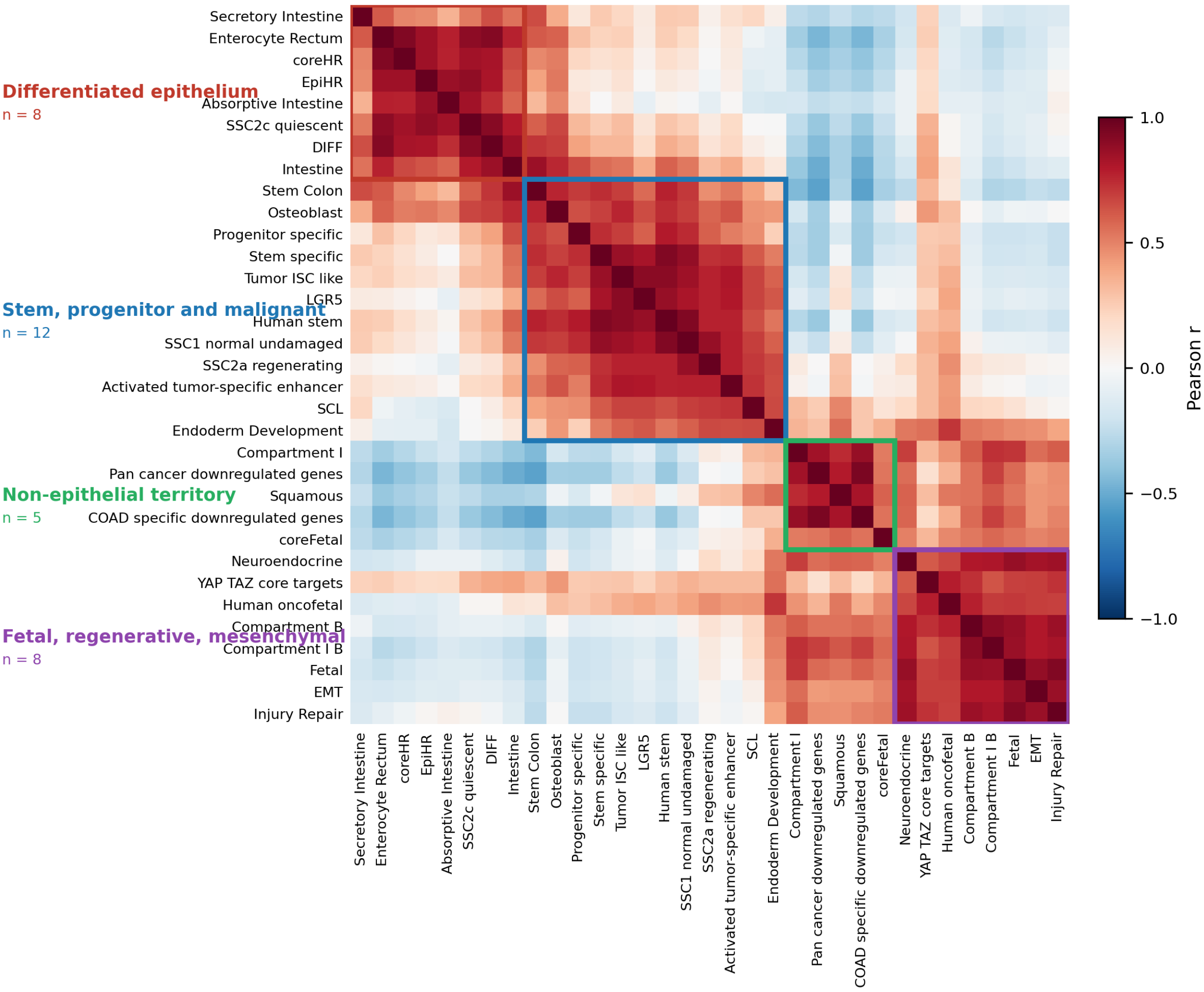
Differentiated epithelium
n = 8
Stem, progenitor and malignant
n = 12
Non-epithelial territory
n = 5
Fetal, regenerative, mesenchymal
n = 8
Secretory Intestine
Enterocyte Rectum
coreHR
EpiHR
Absorptive Intestine
SSC2c quiescent
DIFF
Intestine
Stem Colon
Osteoblast
Progenitor specific
Stem specific
Tumor ISC like
LGR5
Human stem
SSC1 normal undamaged
SSC2a regenerating
Activated tumor-specific enhancer
SCL
Endoderm Development
Compartment I
Pan cancer downregulated genes
Squamous
COAD specific downregulated genes
coreFetal
Neuroendocrine
YAP TAZ core targets
Human oncofetal
Compartment B
Compartment I B
Fetal
EMT
Injury Repair
Pearson r
1.0
0.5
0.0
−0.5
−1.0

N

O

P

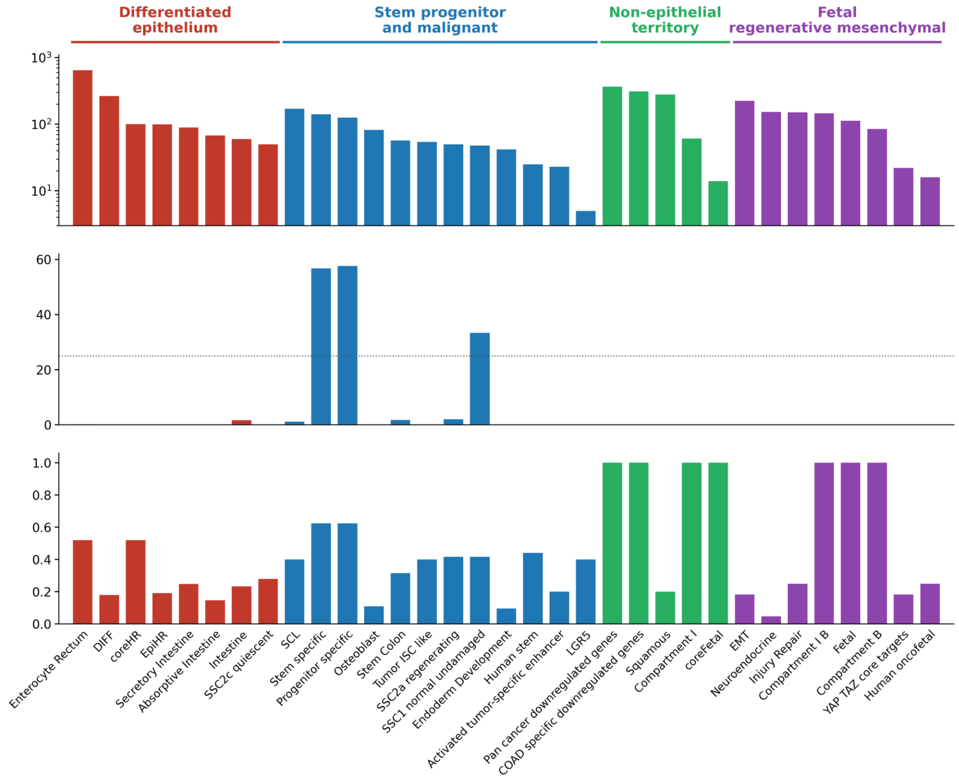
Differentiated epithelium
Stem progenitor and malignant
Non-epithelial territory
Fetal regenerative mesenchymal
$10^3$
$10^2$
$10^1$
60
40
20
0
1.0
0.8
0.6
0.4
0.2
0.0
Enterocyte Rectum
DIFF
coreHR
EpiHR
Secretory Intestine
Absorptive Intestine
Intestine
SSC2c quiescent
SCL
Stem specific
Progenitor specific
Osteoblast
Stem Colon
Tumor ISC like
SSC2a regenerating
SSC1 normal undamaged
Endoderm Development
Human stem
Activated tumor-specific enhancer
LGR5
Pan cancer downregulated genes
COAD specific downregulated genes
Squamous
Compartment I
coreFetal
EMT
Neuroendocrine
Injury Repair
Compartment I B
Fetal
Compartment B
YAP TAZ core targets
Human oncofetal

Q

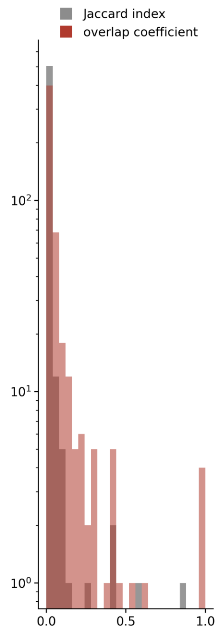
Jaccard index
overlap coefficient
$10^2$
$10^1$
$10^0$
0.0
0.5
1.0

**A,** TRI ranking of the 33 cell-state programs. TRI represents the cumulative path length travelled by each program centroid across ten pseudotime bins in standardized signature-score space. Programs are ordered by decreasing TRI. Colors and asterisks indicate empirical significance determined from 1,000 pseudotime permutations followed by Benjamini–Hochberg correction; grey, not significant; light orange, FDR < 0.05; red, FDR < 0.01.

**B,** Directionality of program remodeling. Points show TDI estimates and horizontal lines show bootstrap 95% confidence intervals obtained from 1,000 within-bin resamples. Red indicates programs that increase along pseudotime and blue indicates programs that decrease. Grey denotes programs below the predefined TRI ≥ 1 threshold, for which TDI was not interpreted.

**C,** Standardized centroid trajectories of all 33 programs across ten pseudotime bins. Programs are classified as DIFF-directed, SCL-directed, fluctuating or stationary according to signed TDI and TRI. The shaded region highlights the coordinated late transition between bins 8 and 9.

**D,** Heatmap of mean standardized program centroids across pseudotime. Programs are ordered by signed TDI. Corresponding signed TDI and TRI values are shown to the right. Programs below the TRI threshold are indicated in grey italics.

**E,** Permutation-based validation of DIFF remodeling. The histogram shows the null distribution of DIFF TRI values obtained from 1,000 pseudotime permutations. The orange line indicates the observed DIFF TRI of 2.181 and the dotted line indicates the maximum permuted value. Empirical P < 0.001; Benjamini–Hochberg-adjusted FDR = 0.00174.

**F,** Late-switch behavior of the DIFF program. The upper plot shows the standardized DIFF centroid across pseudotime bins. The lower plot shows the change in centroid value between consecutive bins, identifying the 8-to-9 transition as the dominant remodeling step.

**G,** Joint DIFF–SCL state-space trajectory. Individual cells are positioned by standardized DIFF and SCL scores and colored by pseudotime bin. Large outlined points represent bin centroids connected in pseudotime order. The cumulative two-dimensional centroid path length was 3.536.

**H–J,** Pairwise program-state spaces used to distinguish reciprocal remodeling from correlated or stationary behavior. Individual cells are colored by pseudotime and large outlined points show bin centroids. (**Fig. S6H**) Human oncofetal versus stem colon, representing weak positive association without marked directional displacement (Pearson $r = 0.186$; Spearman $\rho = 0.185$). (**Fig. S6I**) Human oncofetal versus SCL, representing minimal association (Pearson $r = -0.070$; Spearman $\rho = -0.071$). (**Fig. S6J**) DIFF versus SCL, demonstrating reciprocal remodeling (Pearson $r = -0.493$; Spearman $\rho = -0.483$). Each analysis includes 3,424 cells.

**K**, Projection of all 33 UCell programs onto the published single-nucleus atlas of normal colorectal mucosa, adenoma and CRC (39). Samples were restricted to a filtered subset with sufficient epithelial representation. The cell-state map is shown at upper left, followed by program projections grouped as differentiated epithelium; stem, progenitor and malignant; non-epithelial territory; or fetal, regenerative and mesenchymal programs, with program titles colored by block. SOX9 and KRT20 expression maps are shown last as epithelial reference markers. Values beneath each projection indicate the displayed UCell-score or expression range, which is scaled independently for each panel.

**L,** Epithelial-filtered UCell activity of all 33 programs across normal mucosa (N), adenoma (A) and CRC (T). Analysis was restricted to epithelial and malignant epithelial cells and excluded unaffected tissue. Program scores were averaged within tissue samples and standardized separately for each program across sample means. Each point represents one tissue sample; boxes show the median and interquartile range, and whiskers extend to 1.5 times the interquartile range. All panels share the same y-axis and are ordered from the strongest CRC-minus-normal decrease to the strongest increase. The analysis included 9 normal, 52 adenoma and 6 carcinoma samples from 2, 9 and 6 donors, respectively. DIFF and SCL define the opposing extremes of the ranking, with $\Delta T-N = -3.46$ and $+3.11$, respectively. No inferential P values are shown because the normal samples include multiple regions originating from two donors.

**M,** Pearson correlation matrix of program activities across the colorectal atlas. Hierarchical organization resolves four activity blocks, differentiated epithelium (n = 8); stem, progenitor and malignant programs (n = 12); non-epithelial territory (n = 5); and fetal, regenerative and mesenchymal programs (n = 8).

**N–P,** Gene-content characteristics of the 33 programs, ordered according to the four activity blocks. (**Fig. S6N**) Number of genes in each program, shown on a logarithmic scale. (**Fig. S6O**) Maximum number of shared genes between each program and any other program in the collection; the dashed line marks 25 shared genes. (**Fig. S6P**) Maximum pairwise overlap coefficient for each program.

**Q,** Distribution of pairwise Jaccard indices and overlap coefficients across all 528 program pairs, with counts shown on a logarithmic scale. Most program pairs show little or no gene overlap, 306 of 528 pairs contain no shared genes, and the SCL and DIFF programs are completely disjoint. These analyses show that the activity blocks and the reciprocal SCL–DIFF relationship cannot be explained by direct gene-set overlap alone.
